# Predicting substrates for orphan Solute Carrier Proteins using multi-omics datasets

**DOI:** 10.1101/2024.06.24.600425

**Authors:** Y. Zhang, S. Newstead, P. Sarkies

## Abstract

Solute carriers (SLC) are integral membrane proteins responsible for transporting a wide variety of metabolites, signaling molecules and drugs across cellular membranes. Despite key roles in metabolism, signaling and pharmacology, around one third of SLC proteins are ‘orphans’ whose substrates are unknown. Experimental determination of SLC substrates is technically challenging given the wide range of possible physiological candidates. Here, we develop a predictive algorithm to identify correlations between SLC expression levels and intracellular metabolite concentrations by leveraging existing cancer multi-omics datasets. Our predictions recovered known SLC-substrate pairs with high sensitivity and specificity compared to simulated random pairs. CRISPR loss-of-function screen data and metabolic pathway adjacency data further improved the performance of our algorithm. In parallel, we combined drug sensitivity data with SLC expression profiles to predict new SLC-drug interactions. Together, we provide a novel bioinformatic pipeline to predict new substrate predictions for SLCs, offering new opportunities to de-orphanise SLCs with important implications for understanding their roles in health and disease.

## Introduction

Solute Carriers (SLCs) represent the second largest family of membrane proteins in the human genome after G-protein Coupled Receptors (GPCRs). According to the latest classification database a total of 456 protein-coding genes are classified as a SLC (Ferrada & Superti-Furga, 2022). SLC proteins are localised throughout the cell and regulate the flux of many different classes of small molecules including ions, sugars, amino acids, peptides, vitamins and nucleotides across the cell and organelle membranes (Hediger et al., 2013; Meixner et al., 2020).

SLCs are alpha-helical integral membrane proteins and operate under the influence of ion or metabolite gradients to transport molecules across membranes using an alternating access cycle. Several versions of the alternating access cycle have evolved, however, they all share the same fundamental process: an outward open state, wherein a central binding site is available to the non-cytoplasmic side of the membrane; an occluded state, where ions and/or metabolites are trapped inside the transporter; and an inward-facing state, where the transporter has opened its binding site to the cytoplasm.

The breadth of substrate specificities and subcellular localization of SLCs gives them critical roles in regulating cellular metabolism (Song et al., 2020), energy production (Aulakh et al., 2022; Kunji et al., 2016; Majd et al., 2018), signal transduction (Nimmanon et al., 2017), and the maintenance of physical characteristics such as cell volume (Okada, 2004). Inherited mutations in SLCs have been linked to at least 100 monogenic disorders (Lin et al., 2015). Beyond their physiological substrates, SLCs also transport drug molecules and are thus of crucial importance in pharmacokinetics, drug sensitivity and therapy outcomes (Koepsell & Endou, 2004; Mikkaichi et al., 2004; Winter et al., 2014).

The key roles played by SLCs in regulating cellular metabolism and their potential to control the efficacy of drug treatments mean that characterisation of both physiological and non-physiological substrates of SLCs is of great importance. However, many SLCs have no substrate annotated and are described as ‘Orphan Transporters’. A recent survey reported that around 28% of SLCs had no experimentally determined substrate (Meixner et al., 2020). The key challenge to de-orphanising SLCs is the inability to predict substrates based on sequence or structure alone, as many transporters within the same family recognise chemically diverse molecules (Ferrada & Superti-Furga, 2022). For example, members of the SLC13 family share 40% - 50% amino acid identity, but separate into two different groups based on function: whilst SLC13A1 and SLC13A4 transport sulfates, SLC13A2, SLC13A3 and SLC13A5 recognise carboxylates (Bergeron et al., 2013). Therefore, it is currently not possible to accurately infer SLC substrates from amino acid sequence, and experimental determination of substrates is required. However, screening enough substrates to cover all the possibilities is not practical. Thus, methods that could produce predictions for the likely substrate of any given SLC would be extremely valuable in narrowing down the list of substrates to test in functional assays as well as suggesting insights into cellular functions of uncharacterised SLCs.

Here, we describe a new method to use existing multi-omics high throughput datasets to predict SLC substrates. We demonstrate that our method carries predictive power in recovering known SLC-substrate pairs when applied to multiple major cell line panels. We use our method to generate predictions for orphan SLCs. In parallel, we develop our method to produce new predictions for the effects of SLC expression on sensitivity to drugs. We hope that our predictions act to generate new hypotheses for the substrates, cellular roles, and therapeutic implications of uncharacterised SLC proteins.

## Results

### Correlation analysis between metabolomics and expression datasets successfully predicts SLC substrates

We set out to de-orphanise SLC proteins by investigating the potential effects their expression might have on cellular metabolite concentrations. We reasoned that the transporter activity of SLCs might result in correlations between SLC expression level and the intracellular concentrations of their corresponding substrates (Figure 1A). We used a major cancer cell panel profiling 225 metabolites with liquid chromatography-mass spectrometry (LC-MS) across almost a thousand cancer cell lines from the Cancer Cell Line Encyclopedia 2019 (CCLE2019) (Li et al., 2019). We selected a list of SLC and SLC-like genes (S1 Table) from previous curations (Gyimesi & Hediger, 2022; Meixner et al., 2020). For each SLC or SLC-like gene in the list, we related their normalised transcript levels to metabolite concentration in 913 cell lines, and calculated the Z-score of the absolute values of the correlation coefficients (Spearman’s ρ) for each metabolite to account for varying degrees of correlation strength across different metabolites (Methods). Upon inspection of this set we observed many cases where expression of an SLC correlated most strongly with its known substrate. For example, SLC6A6, a Na /Cl -dependent β-alanine and taurine transporter (Ramamoorthy + - et al., 1994), correlated most strongly to β-alanine and taurine, whilst SLC6A8, a Na /Cl -dependent creatine transporter (Skelton et al., 2011), correlated most strongly to creatine (Figure 1B). Notably, expression of SLC6A8 also strongly correlated with two other metabolites, phosphocreatine and creatinine, which are direct derivatives of creatine (Taegtmeyer & Ingwall, 2013; Wyss & Kaddurah-Daouk, 2000).

**Figure 1.**
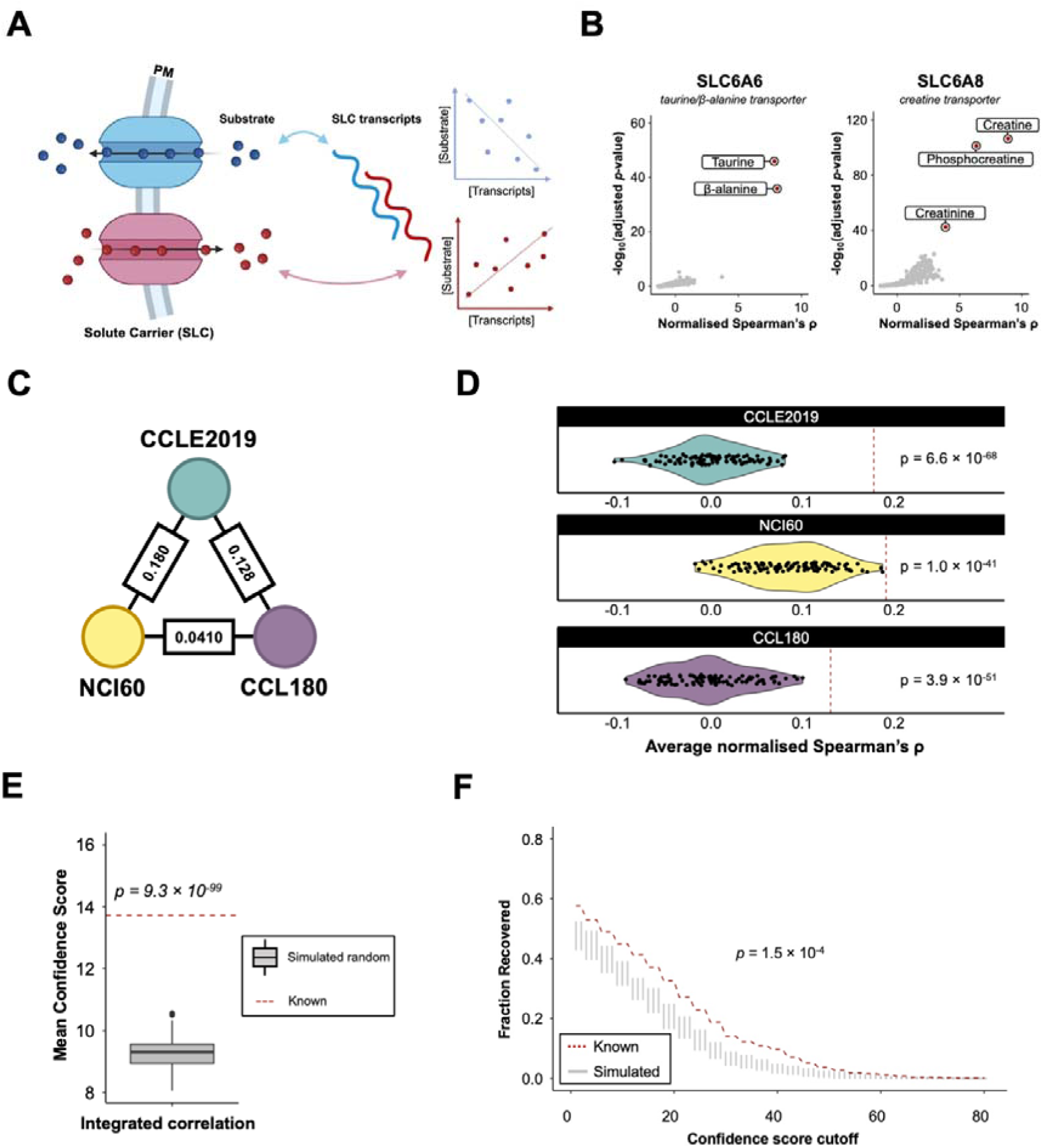
| Correlating SLC transcript levels to substrate concentrations reveals known substrates of SLC. A) Schematic representation shows the principle of correlation analysis. SLC expression levels are hypothesised to correlate with the intracellular concentration of their corresponding substrates. Blue, SLC exporting substrate; red, SLC importing substrate. Figure created with elements from BioRender. B) Scatter plot shows normalised Spearman’s ρ and adjusted p-value for 225 metabolites correlating with the expression level of SLC6A6 and SLC6A8 across 913 cell lines from CCLE. Key substrates are labelled. C) Mutual concordance of SLC-metabolite correlation outcomes from different datasets, each processed with the pipeline specified in Methods. Nodes, datasets, edge, concordance parameter measured in -293 -221 Spearman’s ρ. CCLE2019 and NCI60, p = 4.30 x 10 -24 CCL180, p = 2.38 x 10 . ; CCLE2019 and CCL180, p = 1.50 x 10 ; NCI60 and D) Violin plot shows the mean normalised Spearman’s ρ in known SLC-metabolite set compared to 100 simulated random sets in each dataset. Colored violin and dots, mean normalised Spearman’s ρ of 100 simulated random sets; Red dashed line, mean normalised Spearman’s ρ of known sets; p-value was derived from one-tailed t-test against the null hypothesis that average normalised Spearman’s ρ of known set is not higher than its corresponding 100 simulated random sets. E) Boxplot shows the mean confidence score of known SLC-metabolite sets compared to that of 100 simulated random sets. p-value was derived from one-tailed t-test against the null hypothesis that mean normalised Spearman’s ρ of known set is not higher than its corresponding 100 simulated random sets. F) Across confidence cutoff selected, fraction of interactions recovered (having confidence score better than cutoff) in known set compared to simulated random set. Red dashed line, recovered fraction value in known set; grey violin distribution, recovered fraction value in simulated random set; p-value was derived from one-tailed Wilcoxon test against the null hypothesis that fractions recovered in known set is not higher than its corresponding 100 simulated random sets.

These examples indicated that the expression and metabolite level variation across cancer cell lines might be more generally predictive of the functions of SLCs. To test this, we first expanded the correlation analysis to two other major cell line panels, NCI-60 (Shoemaker, 2006) and CCL180 (Cherkaoui et al., 2022) (Table S2-S4). These three datasets were generated using different methodologies to measure metabolite levels and gene expression; nevertheless, our analysis demonstrated significant concordance in SLC-metabolite correlations between them, suggesting that our method generated robust predictions (Figure 1C). We next investigated how well our correlation method was able to capture the known substrates of SLCs. We updated and expanded the database of SLC annotations based on a previous report (Meixner et al., 2020), selected a list of known substrates that can be found in the metabolomics we used (Table S5; see also Table S1 for references). Since the annotated names for the same metabolites was often different across the databases, we manually examined the metabolomic annotations in the three metabolomics we used to ensure consistent nomenclature across them (Table S6). We tested what fraction of the known substrates were recovered by our correlation analysis and compared this to a random expectation generated by shuffling the SLCs and substrates (“simulated random pairs”). All three datasets had a mean Z-score normalised Spearman’s ρ of known SLC-substrate pairs (Table S5; see also Table S1 for references) significantly higher than that of simulated random pairs (Figure 1D), indicating that correlation analysis was able to successfully predict known interactions.

In order to use our method to generate prediction for novel SLC-substrate pairs, we sought to define a cutoff for the correlation strength indicative of a strong prediction (Methods). To do this we systematically varied the normalized Spearman’s ρ and attempted to maximise the fractional difference between true positive (fraction of the set of known pairs above the cutoff) and false negatives (fraction of the set of simulated random pairs above the cutoff). We defined this threshold separately for each of the three datasets (Figure S1A-C). To unify the correlations from the three datasets we assigned each metabolite/SLC pair with a score such that a higher score corresponded to a stronger prediction. To assign a score for a particular metabolite/SLC pair we first normalized the absolute value of the correlation coefficient to give a Z-score. We then compared the Z-score to the Z-scores of the correlation coefficients for all the experimentally determined metabolite/SLC pairs. To give an example of how the score is calculated, the correlation between SLC6A8 and creatine has a Z-score of 3.08 in the NCI60 panel, 8.89 in the CCLE2019 panel and is not represented in the CCL180 panel because creatine was not measured. 3.08 is within the top 10% of known metabolite/SLC pairs in the NCI60, and is therefore assigned a score of 10; 8.89 is in fact the best correlation within the known set for the CCLE and so is assigned a score of 11. These scores are multiplied by 3 to give a total score of 63 for the creatine/SLC6A8 pair. When compared against the mean confidence score of the known SLC-substrate pairs our score had good predictive power compared with the simulated random sets (Figure 1E) and worked across a range of confidence score cutoffs (Figure 1F). Taken together, these results indicate that correlation analysis is able to provide an accurate indication of SLC-substrate pairs and thus could be a potential method to predict new substrates for orphan SLCs.

### Data from gene dependency screens improves SLC substrate predictions

We next considered whether data from genome-wide gene dependency screens (Bock et al., 2022) could be incorporated into our predictions of SLC substrates. We reasoned that cell growth may be dependent on a specific SLC if the cells grow more slowly after the expression of that SLC is depleted, due to loss of that particular metabolite(s) (Figure 2A). Metabolites whose concentrations are significantly different between dependent and non-dependent cell lines might therefore be candidate substrates for the SLC in question. We used the CRISPR-Cas9 dependency screen (Tsherniak et al., 2017) recording cell growth in over a thousand cell lines upon CRISPR knockdown, of which 625 cell lines overlap with CCLE2019 metabolomics profiling. To infer the dependency of cell lines on specific genes we used the gene effect score as defined previously (Meyers et al., 2017). Genes annotated with negative gene effect scores indicated that cells exhibit reduced proliferation upon their deletion compared to normal cells. For each SLC gene in the curated list, we ranked the cell lines based on their corresponding gene effect scores, excluding positive scores where growth was improved by loss of the SLC. We computed a p-value following multiple test corrections for the difference in each metabolite between cell lines with the top 20% of negative scores (more dependent) and bottom 20% of negative scores (less dependent) (Table S7). Across a range of p-value cutoffs the fraction of significant pairs recovered from the known set is higher than simulated random pairs. The fractional difference is maximised when the p-value cutoff for significance is set to 0.077 (Figure S1D). For p-values smaller than the cutoff, a confidence score is assigned based on its position within a similarly calculated decile-based quantile of known pairs, scaled to 0.1 (Figure S1E). Thus, incorporating the CRISPR-Cas9 dependency screen as an additional data source improved the prediction performance, as the confidence score difference between known pairs and simulated random pairs increased by 7 % (Figure 2C).

**Figure 2.**
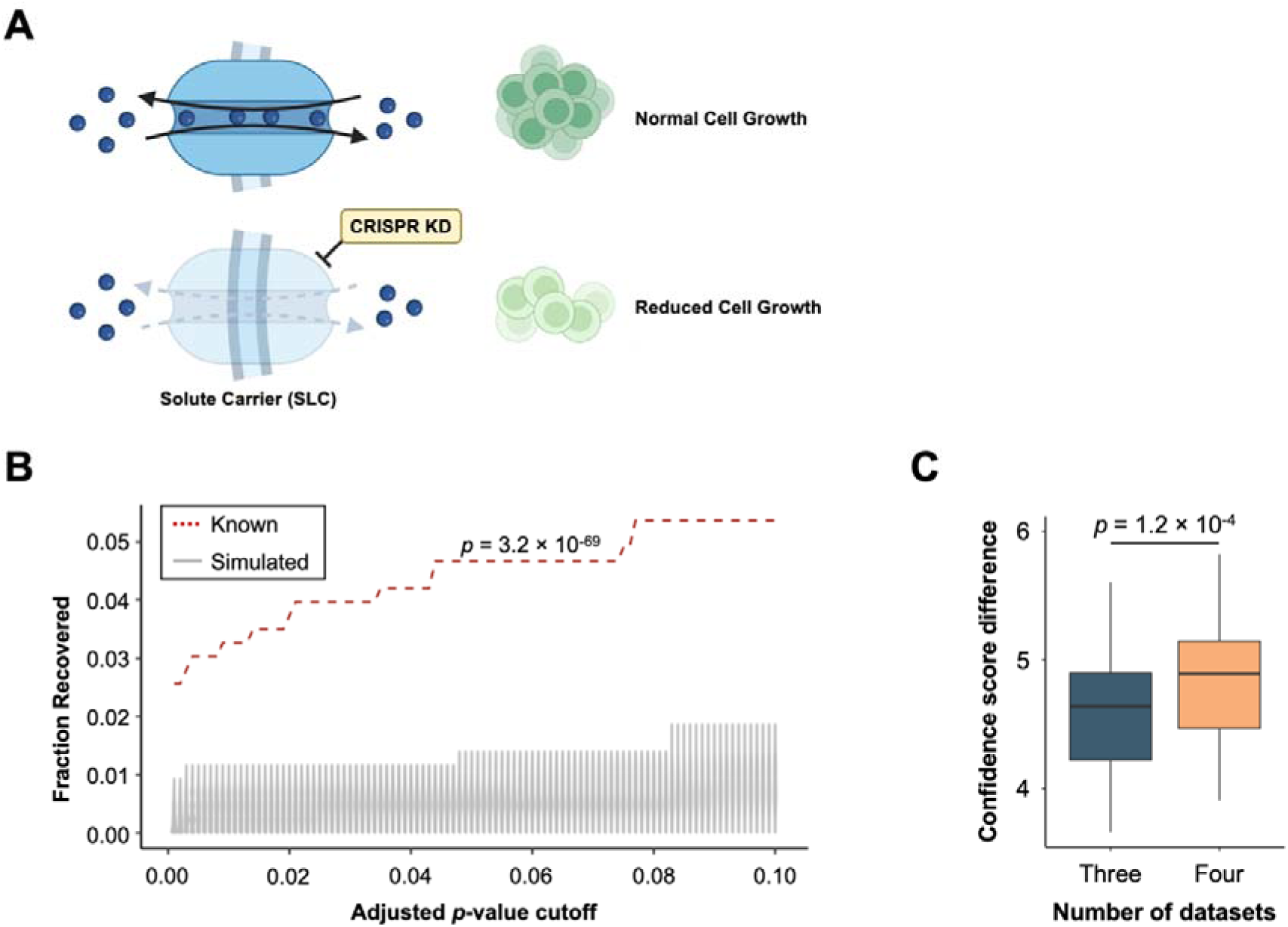
| CRISPR loss-of-function screen provides alternative source for SLC prediction. **A)** Schematic representation shows the principle of CRISPR loss-of-function data analysis to predict SLC substrates. Figure created with elements from BioRender. **B)** Violin plot shows the recovered fractions in known sets compared to simulated random sets across different cutoff selected for resulting adjusted p-values. The p-value comparing the fraction distribution between known and simulated random was derived from one-tailed Wilcoxon test against the null hypothesis that fractions recovered in known set is not higher than its corresponding 100 simulated random sets. **C)** Boxplot showing the effect of incorporation of CRISPR loss-of-function screen into prediction algorithm on the confidence score difference between true positive and false positives. The p-value comparing the confidence score difference was derived from one-tailed Wilcoxon test against the null hypothesis that confidence score difference calculated using both correlation analysis and CRISPR loss-of-function screen (“Four”) is not better than only using correlation analysis (“Three”).

### Inclusion of adjacent metabolites improves substrate prediction for SLCs

Previous results consolidated that the correlation analysis and CRISPR-Cas9 dependency carry predictive power towards recovering known SLC-substrate pairs (references?). However, substrates might easily dissipate into downstream derivatives, leading to poor correlation and reduced predictive power. Furthermore, the prediction of substrates will be reinforced if the SLC correlates with the derivatives of the substrates as well. To address these ideas, we created a metabolite adjacency matrix (Table S8) from annotated KEGG metabolic pathways. This was done by extracting the number of conversion steps required for one metabolite node to reach another, with each unit of adjacency representing a conversion edge between two metabolite nodes. We reasoned that the expression of the SLC that transports the substrate molecule may correlate with its proximal derivatives, while for its distant derivatives, the distribution of correlation strength will be more random and thus more similar to metabolites that are not related to the substrate of the SLC (Figure 3A). To validate this hypothesis, we generated adjacency tables containing the derivatives that represent different steps of conversion away from the original SLC-substrate table, from proximal to distant. For each SLC-derivative pair in the table, we measured the similarity between the correlation of SLC-derivative pairs and the original SLC-substrate pairs by calculating their Spearman’s ρ difference. Subsequently, these differences were compared against the control tables containing randomly sampled non-adjacent metabolites. Non-adjacent metabolites cannot be linked to the substrate node via any continuous path. We performed one-tailed Wilcoxon tests to compare the Spearman’s ρ differences between the adjacent tables and non-adjacent controls, testing the null hypothesis that the differences in adjacent tables are not smaller than non-adjacent controls. Additionally, for each non-adjacent control in the previous comparison, 100 non-adjacent controls were compared with in one-to-one manner to ensure robustness. Our results demonstrate that proximal derivatives (those requiring fewer conversion steps) showed greater similarity to the original substrates compared to distant derivatives (Figure 3B) and that the difference between derivatives and non-adjacent molecules decreased with increasing distance from the substrate (Figure S2). We next determined the optimal correlation coefficient and the increase in confidence score to be added to maximise the difference between known SLC-metabolite pairs and simulated random pairs (Figure S1F-I). This improved the fractional recovery of known SLC-metabolite pairs by 90% (Figure 3D), indicating that metabolite adjacency information bolstered the accuracy of our prediction algorithm.

**Figure 3.**
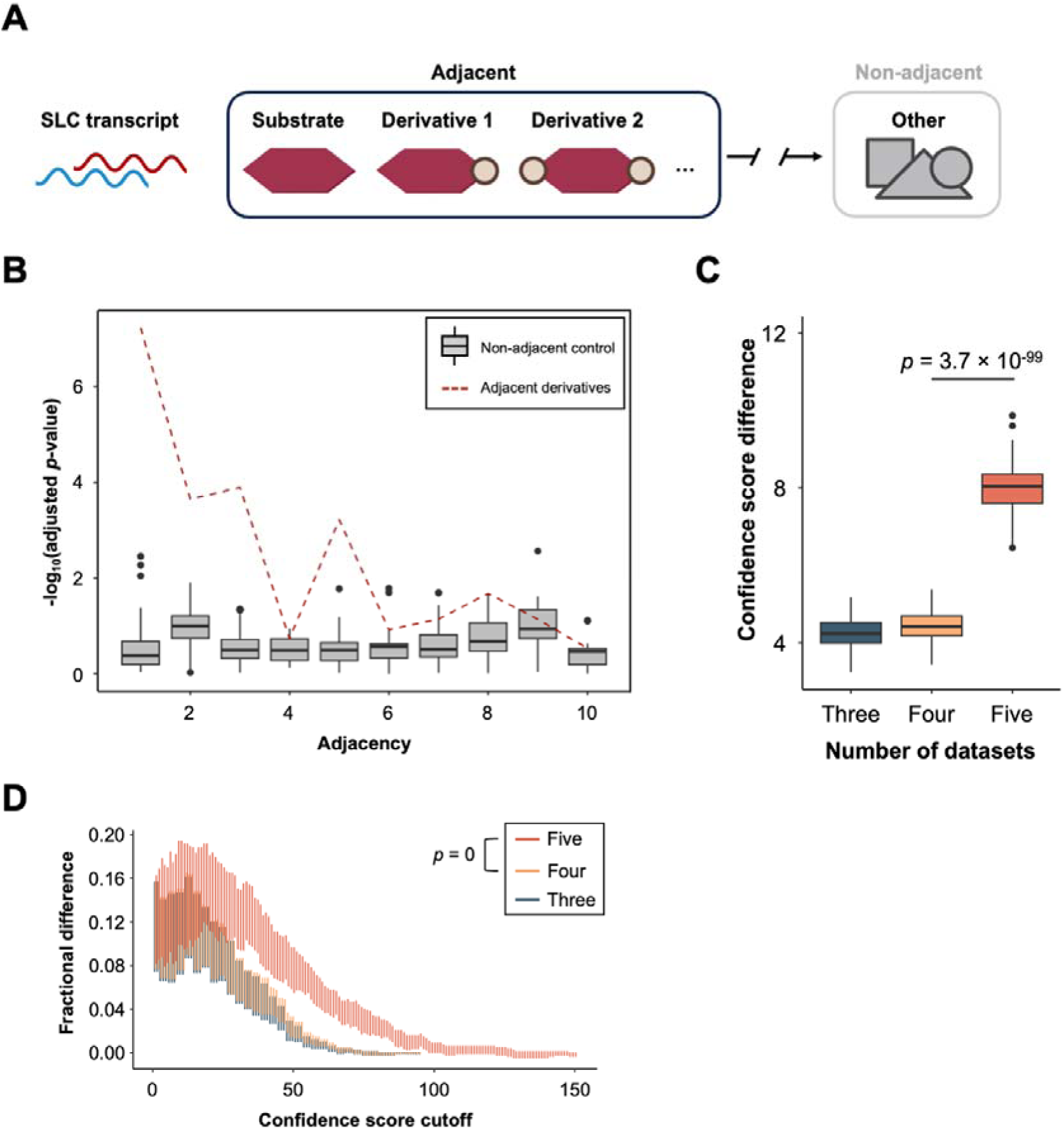
| Inclusion of metabolite adjacency to the prediction substantially differentiates known interaction from simulated random interaction. **A)** Schematic representation shows the relationship between adjacent metabolites (i.e., derivatives of the substrate) and non-adjacent metabolites (i.e., not derivatives of the substrate). Pipeline details specified in **Method**. Figure created with elements from BioRender. **B)** Boxplot shows the general similarity of Spearman’s ρ between derivative across unit of adjacency. Red dashed line, Spearman’s ρ differences between adjacent derivatives and substrates are compared to non-adjacent controls across unit of adjacency. Grey boxes, Spearman’s ρ differences between non-adjacent controls are compared to another 100 non-adjacent controls across unit of adjacency. **C)** Boxplot shows the effect of the incorporation of metabolite adjacency into the prediction algorithm (“Five” vs “Four”). The p-value comparing the confidence score difference was derived from one-tailed Wilcoxon test against the null hypothesis that confidence score difference calculated adding metabolite adjacency (“Five”) is not better than only using correlation analysis (“Four”). **D)** Violin plot shows the fractional difference between true positive and false positives calculated with the inclusion of metabolite adjacency compared to the algorithm without including metabolite adjacency over a range of confidence score cutoffs. The p-value comparing the confidence score difference was derived from one-tailed Wilcoxon test against the null hypothesis that confidence score difference calculated adding metabolite adjacency (“Five”) is not better than only using correlation analysis (“Four”).

### Predicted substrates for Orphan SLCs

Our method thus confirms that true SLC-substrate pairs tend to appear in a higher position compared to simulated random pairs when pairs for each SLC were ranked according to their confidence scores, measured with median rank (Figure 4A). We further demonstrated that the fractional difference peaked if we only considered predictions ranked ≤ 178, with the fraction of true positives reaching 50% (Figure 4B). However, in order to generate a number of predictions for orphan SLCs that could be reasonably tested experimentally, we sought to reduce the number of predictions further. We reasoned that we could improve predictive power for a smaller number of possible substrates by simultaneously identifying over-represented metabolite pathways within the set. We curated a list of 623 metabolites across the three metabolomics datasets that could be linked to 57 metabolic pathways (Table S9). Using the known SLC-metabolite pairs we showed that 20 metabolite predictions were optimal to successfully predict enriched metabolic pathways containing the known substrate (Figure 4C).

**Figure 4.**
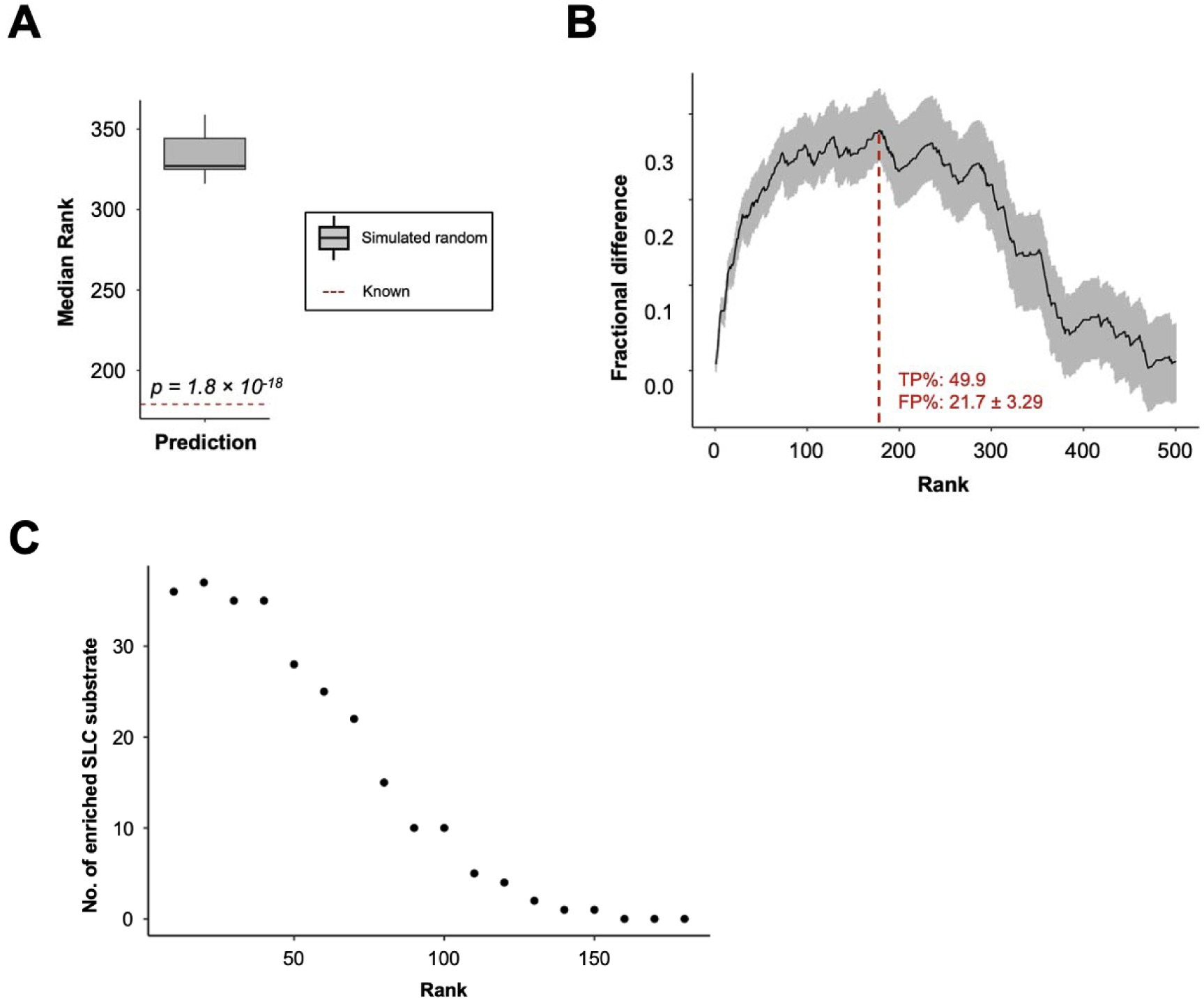
| Top-ranking predictions of known transport activity. **A)** Boxplot shows the median rank of known SLC-substrate pairs compared to simulated random pairs. The p-value comparing the median rank was derived from one-tailed Wilcoxon test against the null hypothesis that the median rank in the known set is not closer to top than in simulated random set. **B)** Fractional difference between true positive (TP%) and false positive (FP%) when only rank above the given value is considered as predicted. Black line, median fractional difference; grey, range of fractional difference. **C)** Point plot shows the number of SLC with substrate converged with the pathway enriched from metabolites ranked above the given rank value in the prediction list.

On this basis we used our prediction algorithm to create a list of substrate predictions with high confidence scores for 128 orphan SLCs (Table S10). We identified many predictions that are in line with experimental data. For example, we found strong associations between the orphan SLC CLN3 and several glycerol phosphate related metabolites (e.g. phosphatidylcholine, alpha-glycerophosphocholine, alpha-glycerophosphate, glycerylphosphorylethanolamine), agreeing with recent research indicating that CLN3 mutant in zebrafish leads to glycerophosphodiesters (GPDs) accumulation in early development (Heins-Marroquin et al., 2024). We predicted that MTCH1 could be associated with metabolites involved in glutathione synthesis (glycine, glutathione, glutamate, pyroglutamate, NADPH), which aligns with the recent observation that MTCH1-deficiency correlates with NAD+ depletion in mitochondria (Wang et al., 2023). Moreover, our results converge with a previous attempt to predict SLC substrate predictions that used sequence information (Meixner et al., 2020). In this publication, SLC25A45, SLC22A25 and SLC35E2B were all predicted to have nucleobase-containing substrates, and our algorithm also predicted a variety of nucleobases as substrates for these transporters (Table S10). Together, our predictions could be used to generate plausible hypotheses for novel SLC substrates, which can be used to narrow down sebsets of metabolites for downstream experimental verification and leading to faster de-orphanisation.

### Leveraging drug repurposing panels provides new predicted SLC-drug interactions

Solute carriers are known to play an important role in determining drug pharmacokinetics, safety and efficacy profiles (Alam et al., 2023). A key goal of the recently established International Transporter Consortium is to identify key transporters involved in drug transport and highlight potential issues around adverse drug-drug interactions involving transporters during clinical trials. Therefore, in parallel to the prediction of physiologically relevant substrates, we investigated whether interrogation of omics datasets could be used to identify drug molecules that are substrates for specific SLC proteins. We reasoned that expression of SLCs might affect drug efficacy, thus alter the shape of the dose-response curve reporting the relationship between viability and drug concentration. For example, when considering anticancer drugs, if cell death is improved or attenuated with higher SLC expression levels, one possible indication is that the drug is a substrate for transport by the SLC in question (Figure 5A). We investigated our hypothesis using the cancer repurposing screen profiling 1448 active drugs against 578 cancer cell lines across 8 doses (Corsello et al., 2020). The cancer cell lines screened were ranked based on each SLC’s transcript levels in CCLE2019, with the highest and lowest 20% marked with “high expression” and “low expression”, respectively. We fitted non-linear regression models into the annotated screen data, and compared the curves between high expression and low expression cell lines. Our analysis captured the difference with accuracy, as it revealed consistency with previously validated results. For example, SLC35F2 expression sensitized cells to the drug YM-155, a known substrate imported by this SLC (Winter et al., 2014). SLC19A2 encodes a plasma membrane thiamine transporter (Dutta et al., 1999), but thiamine uptake is not a dose-dependent factor impacting cell viability (Figure 5B).

**Figure 5.**
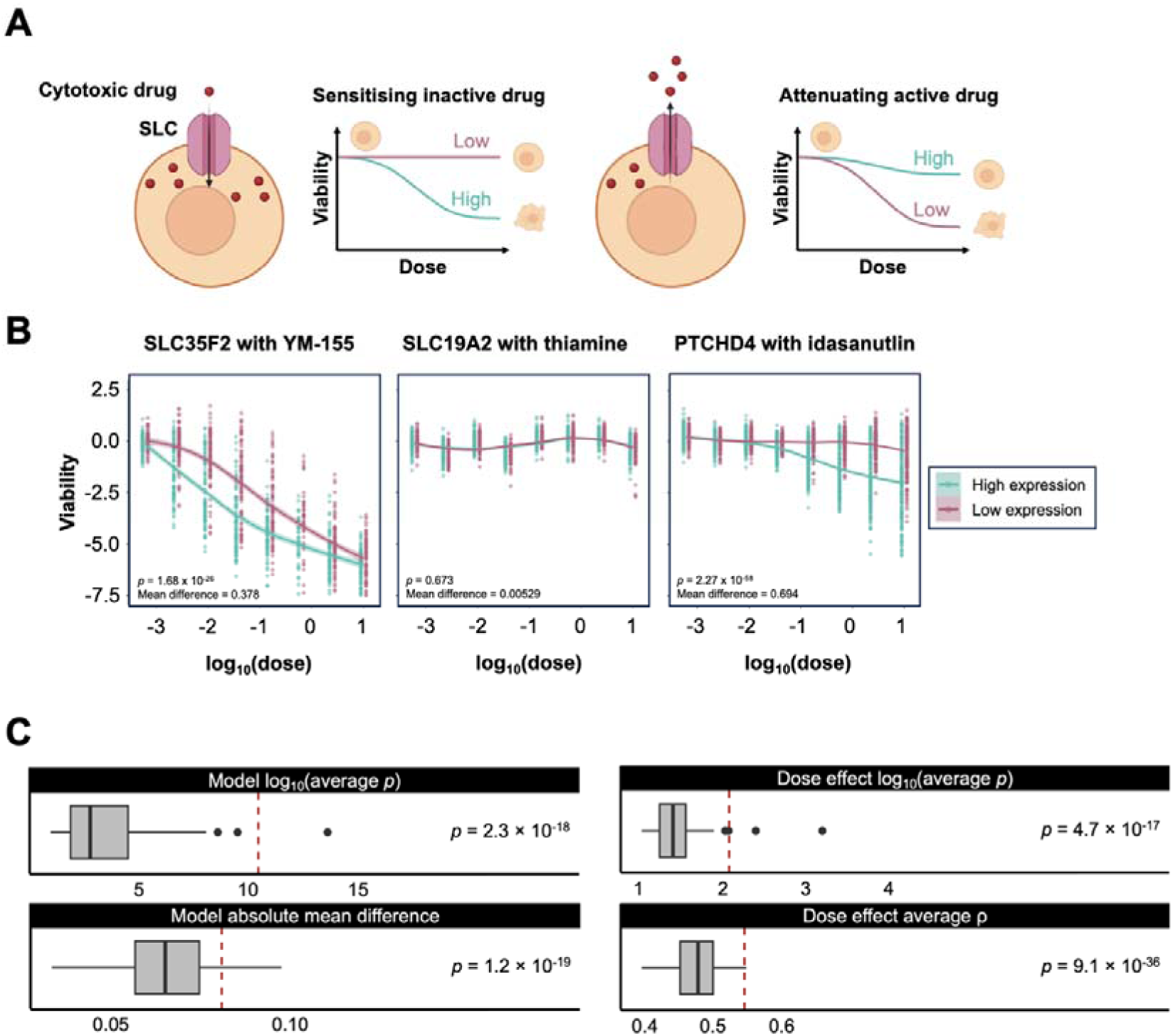
| Combining drug sensitivities and SLC expression profile reveals valuable associations between SLC and drug efficacy. A) Schematic representation shows the principle of leveraging the drug dose response curve to predict SLC-drug associations. If a cytotoxic drug compound is transported by a SLC, expression of the SLC may either enhance or attenuate its killing effect. Figure created with elements from BioRender. B) Non-linear regression shows an example of how SLC35F2 expression affects the killing efficacy of the known substrate drug YM-155 (Left); an example of how SLC19A2 expression does not affect sensitivity to thiamine (Middle); a prediction example of idasanutlin efficacy might associate with PTCHD4, which is not an identified link. Statistics were computed based on paired t-test of model prediction capturing the shape of fitted regression. C) Boxplots show comparison between simulated random interactions and known interactions either through fitting non-linear regression models (Model, left) or performing correlation analysis between dosage and cellular viability (Dose effect, right); p-values derive from one-tailed Wilcoxon test (left) and one-tailed t-test (right) against a null distribution that statistics of the known set is not better than those of the simulated random.

To test if the analysis shows systematic predictive power in the drug repurposing screen, we selected a list of known transport activity of drug molecule by SLCs (Table S11), and used this to benchmark our predictions. Known pairs showed a higher difference between cell lines marked with high and low expression levels in a dosage dependent manner, compared to simulated random pairs (Figure 5C), with optimal p-value cutoff maximising the fractional difference at 0.17 (Figure S1H).

To remove the general impact of drug properties, we calculated a drug-specific significance threshold. For every drug, we randomly picked 20% of cell lines and separated these into high and low expression for 100 times and compared their model predictions. To filter out insignificant pairs we used a drug-specific significant threshold two standard deviations away from the mean of log-transformed p-value (Table S12). Subsequently, drug predictions were listed as we ranked pairs with absolute mean difference and dose-dependent strength, and removed any drug targeting for specific mutations. We then selected the top 50 predictions (Table S13). The SLC-drug pair with the best prediction statistics was an experimentally validated transport activity of YM-155 by SLC35F2 (Winter et al., 2014). Our algorithm also predicted previously unknown links; for example, we predicted an interaction between the orphan SLC Patched Domain Containing 4 (PTCHD4) and the drug molecule idasanutlin, which acts as a small molecule antagonist of p53 activity suppressor Mouse double minute 2 homolog (MDM2) (Figure 5B; Ding et al., 2013). We also noticed a group of SLCs (SLC3A1, SLC7A7, SLC16A4, SLC23A1, SLC37A1, SLC37A2, SLC41A2, NPC1L1, CLN3) that interact with the small molecule inhibitor RITA, which leads to induction of cell apoptosis by (re)activating wild-type or mutant p53 (Wiegering et al., 2017). SLC3A1 associates with attenuation while the other associate with sensitisation of drug killing effect (Figure S3A). Importantly, our predictions worked across a range of cell lines (Figure S3B), demonstrating…. In summary, our work provided a possible route to predict SLC-drug interactions in parallel to physiological substrate determination, aiding the process of exploring SLC as a therapeutic target reservoir or alerting drug discovery teams to potential downstream issues with cell toxicity or adverse impacts on drug pharmacokinetics.

## Discussion

Deorphanisation of SLCs has been a large collective effort in the community (César-Razquin et al., 2015; Superti-Furga et al., 2020). However, experimental substrate determination is hindered by the technical difficulties in expressing and purifying functional membrane proteins and the huge range of potential compounds that could be tested even if the SLC is isolated for functional study. Therefore, accurate prediction of SLC substrates would be an important development for the field. Here we developed a new method to predict SLC substrates, and demonstrated we can recover known interactions, indicating its potential for deorphanisation of SLCs. Below, we discuss how the algorithm compares to previous methods, its strengths and limitations, and prospects for future improvement.

An obvious approach to attempt to predict SLC substrates is to utilise structures of transporters with known substrates to generate a set of rules that could be subsequently used to predict substrates for orphan SLCs. This approach was attempted by Meixner et al. (2020), who trained a machine learning algorithm with systematic SLC substrate annotations and structural features, such as sequence and topological domains. Probabilities were produced for 115 orphan SLCs against 18 selected substrate terms. Over the subsequent 3 years, substrates have been experimentally defined for 28 of these orphan SLCs, offering the opportunity to evaluate this method (Table S14). Prediction of the substrates of 4 SLCs aligned with experimental results (SLC39A11, TMEM165, SLC16A6, SLC16A17). For example, TMEM165, now characterised as a lysosomal Ca^2+^ importer (Zajac et al., 2024), was predicted to have high probability in transporting divalent metal cations. SLC6A17, demonstrated to transport glutamine in mice synaptic vesicles (Jia et al., 2023), was predicted to transport L-amino acids. However, for the remainder of SLCs where substrates were determined, the algorithm either did not provide any predictions (16/28), or the predictions produced do not match experimental determination completely (8/28). For example, ANKH was predicted to have high probability of transporting metal ions, but turned out to mediate ATP and citrate export (Szeri et al., 2020). This evaluation showed that the prediction based on structural features has limitations.

Our method disregarded SLC’s structural features entirely, and focused on the how varying levels of SLC expression affects metabolite levels. Our approach is completely agnostic about known biochemical properties, which means that it has the potential to predict surprising or unexpected interactions, even if an experimentally defined substrate has been annotated. This could be important as many substrates are defined experimentally using a limited range of compounds and using in vitro assays, therefore may not capture the full spectrum of activities for an SLC in vivo. For example, inositol was found to have great correlation with recently characterised facilitative taurine transporter SLC16A6 (Higuchi et al., 2022) in all three datasets, but not with other SLC16 family members, potentially indicating of that this may be an additional substrate of SLC16A6.

The biochemical naivety of our model also presents limitations. As we rely on metabolomics data sets, our method is limited to metabolites measured and prevalent in these experiments. Most notably, ion concentrations are not measured, therefore we cannot predict ion transporter activity, an obvious drawback since many SLCs transport small ions either exclusively or as counterions to drive transport of another metabolite (Pizzagalli et al., 2021). Moreover, many of the correlations with metabolites may reflect secondary effects downstream of the primary substrate which is either not measured or itself rapidly metabolised to other substrates. Our inclusion of adjacency information (Figure X) may go some way towards addressing this; nevertheless it means that predicted substrates should be taken as an indication of the group of possible substrates rather than a clearly defined single substrate. Thus, at present we would suggest that our method offers an opportunity to generate predictions which would still have to be validated with experiments in vitro with purified proteins. Our predictions thus present a resource to the community to expedite experimental substrate identification.

In addition to identifying physiological substrates for SLCs, we also used omics data to identify potential drugs that might be substrates. A previous, experimental, approach to this, screened the impact of knocking out specific SLCs on 60 representative cytotoxic drugs, highlighting the broad role of SLCs in drug efficacy (Girardi et al., 2020). Of the 201 prominent associations (47 drugs, 101 SLCs) they reported, 39 drugs and 97 SLCs are also included in our analysis. Two-thirds (26/39) of these drugs were found to have associated SLCs predicted in our study, with five associated SLCs ranked highly in our predictions: SLC1A4 & Triptolide (3^rd^ out of 92), SLC19A1 & Methotrexate (6^th^ out of 72), SLC2A1 & Idarubicin (6^th^ out of 58), SLC15A1 & 6-Mercaptopurine (7^th^ out of 61), and SLC12A4 & 5-Azacitidine (11 out of 86). The remaining 13 drug associations were eliminated as they fell outside the range of drug-specific thresholds; however, half of these would have associated SLCs found in the top 10% of predictions for each drug, indicating concordance and predictive power.

Our analysis could be a good complement to this previous screen by including a much larger dataset of cell lines and types (469 cell lines compared to 1 in the previous study). Additionally, we examined the effect of individual SLC expression, which might be masked by severe knockout phenotypes. By disregarding structural considerations in our analysis, we allow for the emergence of unexpected drug associations. However, this approach might lead to predicted drug associations not directly related to SLC transport of the drug itself, such as the metabolic environment of the cell and downstream events of SLC activity. Nevertheless, our analysis could aid in characterizing unknown SLC-drug interactions. The potential for polymorphisms in SLCs within human populations has been demonstrated to be a promising angle for personalized medicine(Giacomini et al., 2022); our drug interaction results potentially extend this to include differences in expression as a method to predict the sensitivity of specific tumors to particular drugs in personalized medicine.

## Methods

### Data acquisition for CCLE2019 dataset

CCLE2019 RNA-Seq and metabolomics data were downloaded from the DepMap Portal (depmap.org, CCLE 2019 omics) as read counts (file “CCLE_RNAseq_genes_counts_20180929.gct.gz”) and mean concentration levels (file “CCLE_metabolomics_20190502.csv”).

### Data acquisition for NCI60 dataset

NCI60 RNA-Seq data was derived from alignment and normalisation as performed previously (Perez & Sarkies, 2023). NCI-60 metabolomics data were downloaded from the NCI DTP Data Portal (wiki.nci.nih.gov) as mean concentration levels (file “WEB_DATA_METABOLON.ZIP”).

### Data acquisition for CCL180 dataset

CCL180 metabolomics data were downloaded from the ETH Research Data Collection (https://doi.org/10.3929/ethz-b-000511784) as concentration levels and annotations (file “primary analysis (metabolomics)”).

### Data acquisition for CRISPR-Cas9 dependency screen

CRISPR-Cas9 dependency screen dataset was downloaded from DepMap portal (depmap.org, DepMap Public 23Q2 omics) as gene effect scores (file “CRISPRGeneEffect.csv”).

### Data acquisition for drug repurposing screen

Drug repurposing screen dataset was downloaded from Cancer Dependency Map Portal (https://depmap.org/repurposing), including cell line annotation (file “secondary-screen-cell-line-info.csv”), treatment metadata (“secondary-screen-replicate-collapsed-treatment-info.csv”) and viability log-fold (“secondary-screen-replicate-collapsed-logfold-change.csv”).

### Correlating SLC expression levels to metabolites

Prior to all analysis, RNA-Seq read counts were normalised with Median Ratio Normalisation (MRN) by ‘DESeq2’ package in R to account for gene expression difference across different tissue types and cancer cell lines. Normalisations were applied to both CCLE 2019 and NCI60 raw counts across all cell lines. The data was first converted to a DESeqDataSet (dds) object using ‘DESeqDataSetFromMatrix()’ function, and the sum of gene reads in each cell line was calculated and filtered if lower than 10. The resulting dds object was normalised by applying ‘estimateSizeFactors()’ function, and the normalised pseudocounts were extracted by ‘counts()’ function with argument ‘normalized = TRUE’. All subsequent analyses used the resulting normalised pseudocounts.

Correlation analysis was applied between CCLE 2019 pseudocounts and CCLE 2019 metabolomics (“CCLE2019”), CCLE 2019 pseudocounts and CCL180 metabolomics (“CCL180”), NCI-60 pseudocounts and NCI-60 metabolomics (“NCI60”). Spearman’s correlations were computed across mutually overlapping cell lines between pseudocounts and metabolite levels using the ‘cor.test()’ function in R with argument ‘method = “spearman”’.

Resultingcorrelation p-values were adjusted for each gene using Benjamini-Hochberg Procedure using the ‘p.adjust()’ function with ‘method = “BH”’. Correlation coefficients (ρ) might not be able to present correlation strength accurately across dataset due to the change of correlation distributions. Therefore, for metabolite a and SLC z, the normalised ρ coefficient 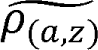 is computed from the following formula:

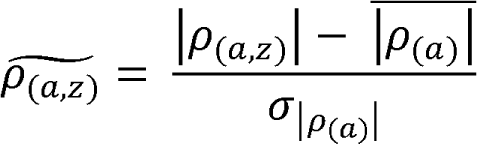

where each *p_(a,z)_* will be transformed to represent the number of absolute standard deviation 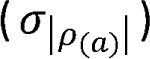 away from the absolute mean 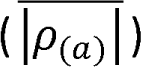 based on the correlation distribution of metabolite a to every SLC, and thus represent only correlation strength of the pair.

### Concordance assessments

Between datasets, only mutually overlapping SLC and metabolite terms were assessed. The resulting raw ρ values for each overlapping SLC and metabolite were taken and correlated using the ‘cor.test()’ function in R with argument ‘method = “spearman”’.

### Benchmarking

Known pair tables (Table S5, S11) were manually extracted from the SLC ontology annotation (Table S1) for overlapping SLC and metabolite terms. Metabolites or drug molecules appearing in the known pair tables were shuffled and randomly assigned to SLCsthat are not known to transport it, while keeping the SLC column unchanged, generating 100 simulated random pair tables. Mean statistics (e.g. normalised ρ, adjusted p, confidence score) were calculated per table to measure predictive power. Across threshold of discovery for corresponding statistics, fractional difference was calculated as the difference between true positive fraction (fraction left in known pair tables) and false positive fraction (fraction left in simulated random pair tables) to measure the validity of threshold chosen.

To benchmark the drug repurposing screen predictions, the difference between “high expression” and “low expression” cell lines for each SLC were compared to the difference between 100 sets of the same numbers of randomly selected cell lines. A value that two standard deviation away from the mean -log_10_(p-value) and absolute mean difference was taken per drug as the filtering threshold.

### Metabolite adjacency

Metabolite adjacency table (Table S8) was generated from human KEGG pathway by ‘MetaboSignal’ package in R (Rodriguez-Martinez et al., 2016). Specifically, all human metabolic pathways were extracted and subsetted using ‘MS_getPathIds()’ function with argument ‘organism_code = “hsa”’. Reaction network was built based on metabolic pathways using ‘MS_reactionNetwork()’ function. Subsequently, node distance was calculated using ‘MS_nodeBW()’ function with argument ‘node = “out”’ and ‘normalized = TRUE’.

### Metabolite prediction algorithm

For every SLC-metabolite pair i, the prediction algorithm computes a confidence score by evaluating its correlation statistics across the three datasets (A_i_), resulted adjusted p-value of annotated CRISPR-Cas9 screen (B_i_), and adjacent metabolites (c_i_). Threshold of discovery and score reward per discovery is specified in Figure S1, validified with fractional difference between true positive and false positive. For every SLC-metabolite pair, if value implicated in A_i_, B_i_, or c_i_ is smaller than its respective threshold of discovery or the metabolite is not measured in the dataset not exist, a confidence score of 0 was assigned. Otherwise, the confidence score will be measured with respect to the decile-based percentile of known SLC-substrate tables, such that a decile of 10 corresponds to the top 10% and a decile of 1 the lowest 10%; correlations greater than or equal to the top correlation (i.e. greater than decile 10) were assigned a score of 11. The adjusted p-value of B_i_was measured in log-transformed format. The decile number was multiplied by 3 for A_i_ , 1.1 for B_i_, and 0.9 for c_i_.

### Drug prediction algorithm

For each pair between drug molecule a and SLC z, the dose responses were only calculated for cell lines annotated based on the highest and lowest 20% of SLC z expression. Local polynomial regression models were fitted to the two expression types using ‘loess()’ function, viability against log transformed dose (−3.21 to 1). 421 data points, ranging from - 3.21 to 1 and separated by 0.01, predicted by model using ‘predict()’ function was generated to capture the shape information of fitted models. Curve shapes were compared with a paired t-test. The resulted p-value and mean differences were recorded.

### Code availability

All the code for processing data, as well as generating the figures and tables in the manuscripts, is available as supplemental material and will be uploaded to GitHub upon publication.

## Supporting information

Code for plotting results

Code for generating results

## Acknowledgements

We thank the members of the EpiEvo group and other colleagues at the Department of Biochemistry, University of Oxford for helpful discussion and comments on the project. We thank Dr Marcos Francisco Perez for aligning and normalising raw NCI-60 transcriptomics data. We also thank Dr Louise Fets (London Institute of Medical Sciences) for helpful discussions.

---

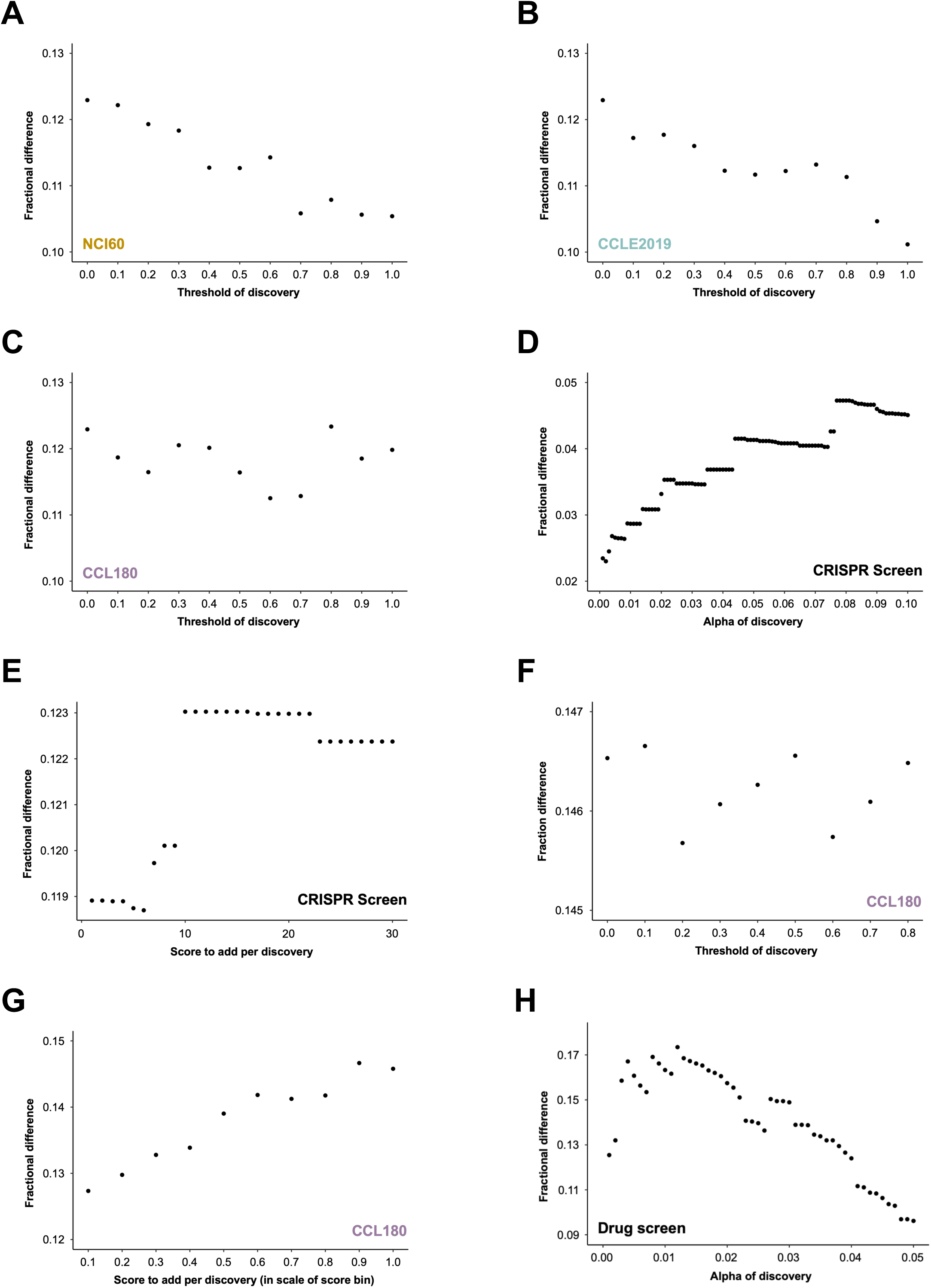
Figure S1

**Figure S2.**
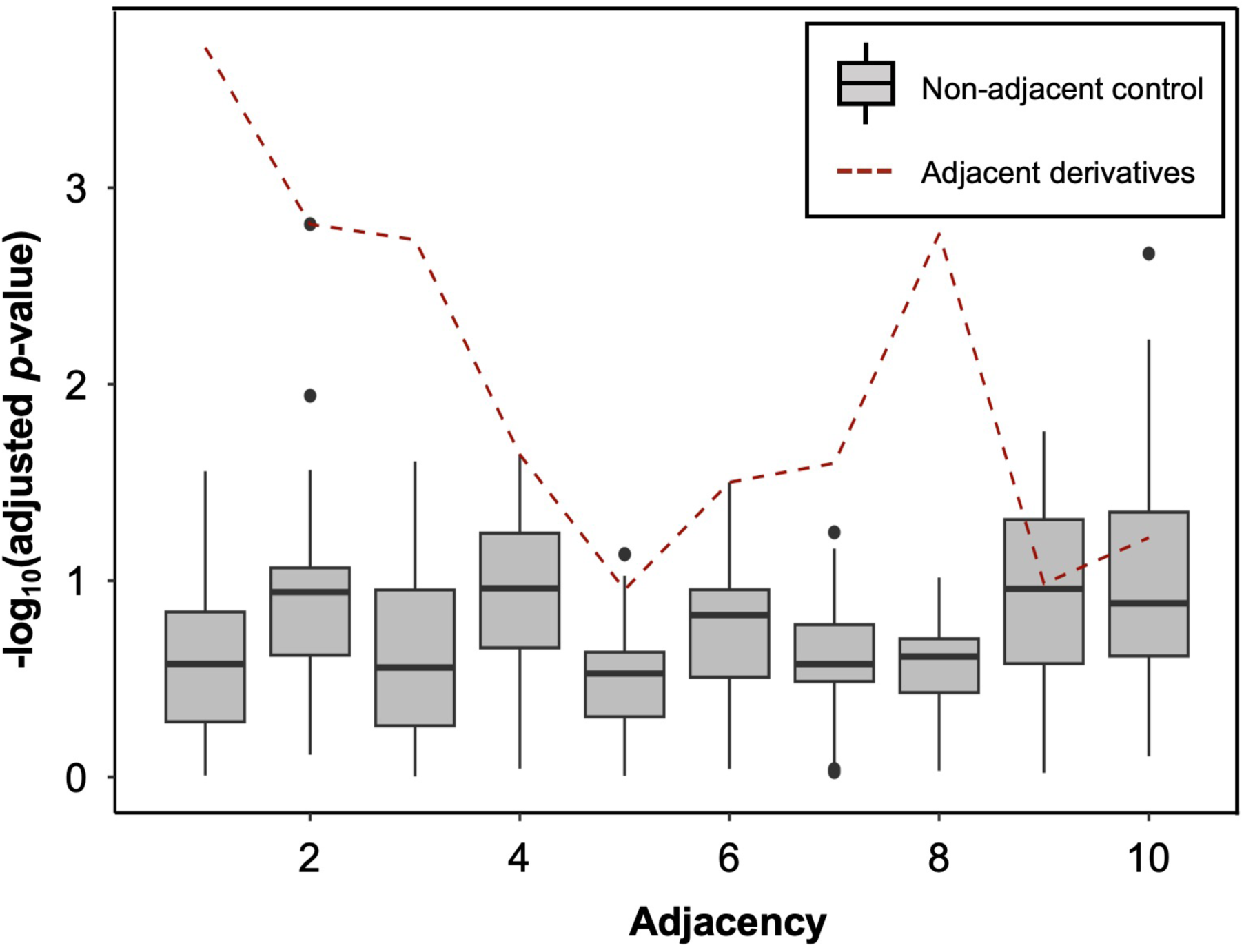
Adjacency normalised rho

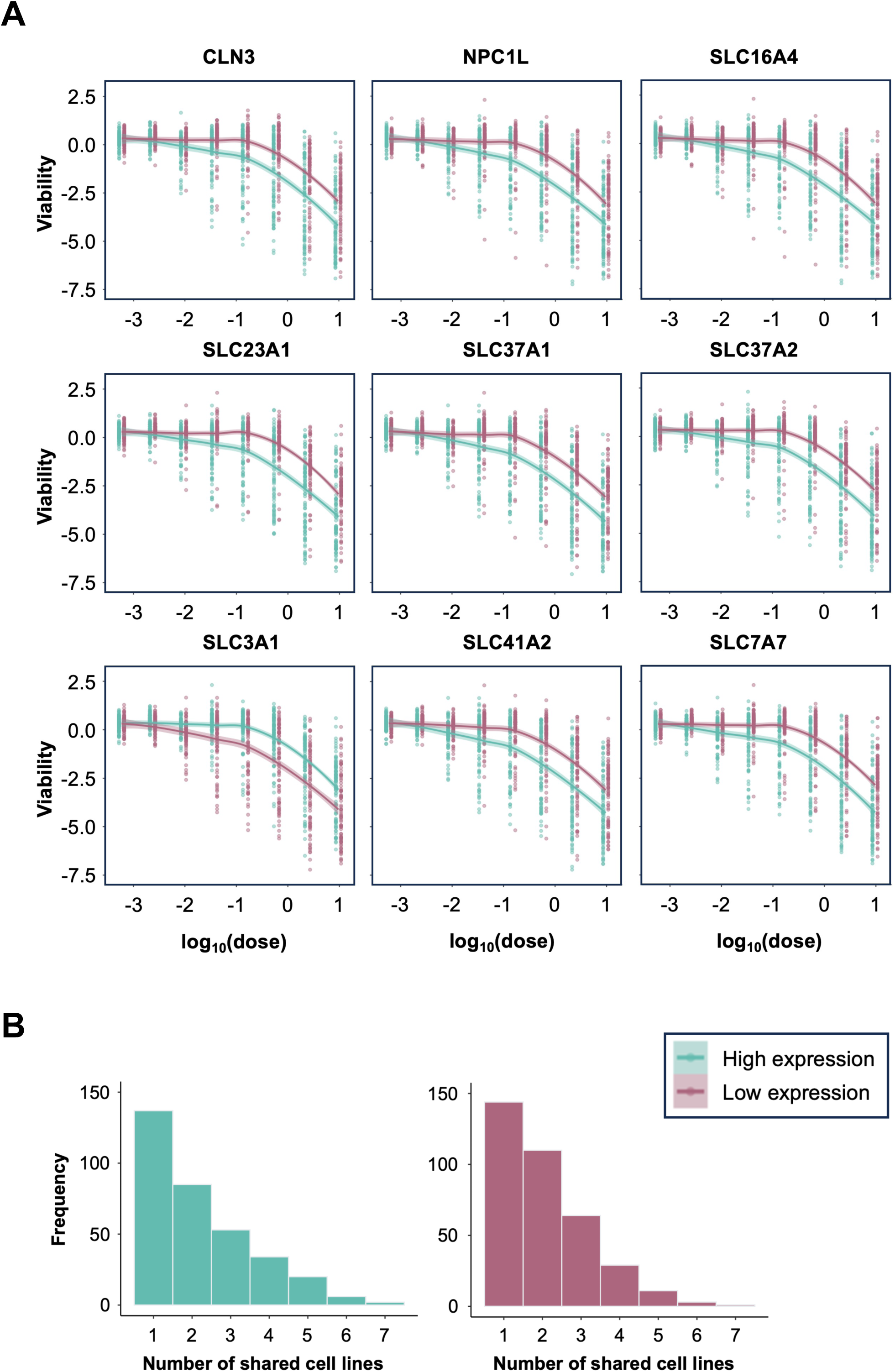
Figure S3

| Table | Description |
| --- | --- |
| S1 | Comprehensive SLC review |
| S2 | CCLE2019 correlation table & normalised table |
| S3 | NCI60 correlation table & normalised table |
| S4 | CCL180 correlation table & normalised table |
| S5 | Extracted SLC known interaction |
| S6 | Metabolite annotation standardisation |
| S7 | CRISPR result table |
| S8 | Adjacency matrix |
| S9 | KEGG metabolic pathway definition |
| S10 | Algorithm interaction prediction |
| S11 | Extracted SLC-drug known interaction |
| S12 | Pseudo-SLC expression simulation threshold |
| S13 | Algorithm interaction prediction (drug) |
| S14 | Recently de-orphanised SLC |

## References

1. Alam, S., Doherty, E., Ortega-Prieto, P., Arizanova, J., & Fets, L. (2023). Membrane transporters in cell physiology, cancer metabolism and drug response. Disease Models & Mechanisms, 16(11). 10.1242/dmm.050404

2. Aulakh, S. K., Varma, S. J., & Ralser, M. (2022). Metal ion availability and homeostasis as drivers of metabolic evolution and enzyme function. Current Opinion in Genetics & Development, 77, 101987. 10.1016/j.gde.2022.101987

3. Bergeron, M. J., Clémençon, B., Hediger, M. A., & Markovich, D. (2013). SLC13 family of Na+-coupled di- and tri-carboxylate/sulfate transporters. Molecular Aspects of Medicine, 34(2–3), 299–312. 10.1016/j.mam.2012.12.001

4. Bock, C., Datlinger, P., Chardon, F., Coelho, M. A., Dong, M. B., Lawson, K. A., Lu, T., Maroc, L., Norman, T. M., Song, B., Stanley, G., Chen, S., Garnett, M., Li, W., Moffat, J., Qi, L. S., Shapiro, R. S., Shendure, J., Weissman, J. S., & Zhuang, X. (2022). High-content CRISPR screening. In Nature Reviews Methods Primers (Vol. 2, Issue 1). Springer Nature. 10.1038/s43586-021-00093-4

5. César-Razquin, A., Snijder, B., Frappier-Brinton, T., Isserlin, R., Gyimesi, G., Bai, X., Reithmeier, R. A., Hepworth, D., Hediger, M. A., Edwards, A. M., & Superti-Furga, G. (2015). A Call for Systematic Research on Solute Carriers. Cell, 162(3), 478–487. 10.1016/j.cell.2015.07.022

6. Cherkaoui, S., Durot, S., Bradley, J., Critchlow, S. E., Dubuis, S., Masiero, M., Wegmann, R., Snijder, B., Othman, A., Bendtsen, C., & Zamboni, N. (2022). A functional analysis of 180 cancer cell lines reveals conserved intrinsic metabolic programs. Molecular Systems Biology, 18(11). 10.15252/msb.202211033

7. Corsello, S. M., Nagari, R. T., Spangler, R. D., Rossen, J., Kocak, M., Bryan, J. G., Humeidi, R., Peck, D., Wu, X., Tang, A. A., Wang, V. M., Bender, S. A., Lemire, E., Narayan, R., Montgomery, P., Ben-David, U., Garvie, C. W., Chen, Y., Rees, M. G., … Golub, T. R. (2020). Discovering the anticancer potential of non-oncology drugs by systematic viability profiling. Nature Cancer, 1, 235–248. 10.1038/s43018-019-0018-6

8. Dutta, B., Huang, W., Molero, M., Kekuda, R., Leibach, F. H., Devoe, L. D., Ganapathy, V., & Prasad, P. D. (1999). Cloning of the Human Thiamine Transporter, a Member of the Folate Transporter Family *. Journal of Biological Chemistry, 274(45), 31925–31929. 10.1074/jbc.274.45.31925

9. Ferrada, E., & Superti-Furga, G. (2022). A structure and evolutionary-based classification of solute carriers. IScience, 25(10), 105096. 10.1016/j.isci.2022.105096

10. Giacomini, K. M., Yee, S. W., Koleske, M. L., Zou, L., Matsson, P., Chen, E. C., Kroetz, D. L., Miller, M. A., Gozalpour, E., & Chu, X. (2022). New and Emerging Research on Solute Carrier and ATP Binding Cassette Transporters in Drug Discovery and Development: Outlook From the International Transporter Consortium. Clinical Pharmacology & Therapeutics, 112(3), 540–561. 10.1002/CPT.2627

11. Girardi, E., César-Razquin, A., Lindinger, S., Papakostas, K., Konecka, J., Hemmerich, J., Kickinger, S., Kartnig, F., Gürtl, B., Klavins, K., Sedlyarov, V., Ingles-Prieto, A., Fiume, G., Koren, A., Lardeau, C.-H., Kumaran Kandasamy, R., Kubicek, S., Ecker, G. F., & Superti-Furga, G. (2020). A widespread role for SLC transmembrane transporters in resistance to cytotoxic drugs. Nature Chemical Biology, 16(4), 469–478. 10.1038/s41589-020-0483-3

12. Gyimesi, G., & Hediger, M. A. (2022). Systematic in silico discovery of novel solute carrier-like proteins from proteomes. PLOS ONE, 17(7), e0271062–e0271062. 10.1371/journal.pone.0271062

13. Hediger, M. A., Clémençon, B., Burrier, R. E., & Bruford, E. A. (2013). The ABCs of membrane transporters in health and disease (SLC series): Introduction. Molecular Aspects of Medicine, 34(2–3), 95–107. 10.1016/J.MAM.2012.12.009

14. Heins-Marroquin, U., Singh, R. R., Perathoner, S., Gavotto, F., Ruiz, C. M., Patraskaki, M., Gomez-Giro, G., Borgmann, F. K., Meyer, M., Carpentier, A., Warmoes, M. O., Jäger, C., Mittelbronn, M., Schwamborn, J. C., Cordero-Maldonado, M. L., Crawford, A. D., Schymanski, E. L., & Linster, C. L. (2024). CLN3 deficiency leads to neurological and metabolic perturbations during early development. Life Science Alliance, 7(3), e202302057–e202302057. 10.26508/lsa.202302057

15. Higuchi, K., Sugiyama, K., Tomabechi, R., Kishimoto, H., & Inoue, K. (2022). Mammalian monocarboxylate transporter 7 (MCT7/Slc16a6) is a novel facilitative taurine transporter. The Journal of Biological Chemistry, 298(4), 101800. 10.1016/j.jbc.2022.101800

16. Jia, X., Zhu, J., Bian, X., Liu, S., Yu, S., Liang, W., Jiang, L., Mao, R., Zhang, W., & Rao, Y. (2023). Importance of glutamine in synaptic vesicles revealed by functional studies of SLC6A17 and its mutations pathogenic for intellectual disability. ELife, 12, RP86972. 10.7554/eLife.86972

17. Koepsell, H., & Endou, H. (2004). The SLC22 drug transporter family. Pflügers Archiv: European Journal of Physiology, 447(5), 666–676. 10.1007/s00424-003-1089-9

18. Kunji, E. R. S., Aleksandrova, A., King, M. S., Majd, H., Ashton, V. L., Cerson, E., Springett, R., Kibalchenko, M., Tavoulari, S., Crichton, P. G., & Ruprecht, J. J. (2016). The transport mechanism of the mitochondrial ADP/ATP carrier. Biochimica et Biophysica Acta (BBA) - Molecular Cell Research, 1863(10), 2379–2393. 10.1016/j.bbamcr.2016.03.015

19. Li, H., Ning, S., Ghandi, M., Kryukov, G. V, Gopal, S., Deik, A., Souza, A., Pierce, K., Keskula, P., Hernandez, D., Ann, J., Shkoza, D., Apfel, V., Zou, Y., Vazquez, F., Barretina, J., Pagliarini, R. A., Galli, G. G., Root, D. E., … Sellers, W. R. (2019). The landscape of cancer cell line metabolism. Nature Medicine, 25(5), 850–860. 10.1038/s41591-019-0404-8

20. Lin, L., Yee, S. W., Kim, R. B., & Giacomini, K. M. (2015). SLC transporters as therapeutic targets: Emerging opportunities. In Nature Reviews Drug Discovery (Vol. 14, Issue 8, pp. 543–560). Nature Publishing Group. 10.1038/nrd4626

21. Majd, H., King, M. S., Smith, A. C., & Kunji, E. R. S. (2018). Pathogenic mutations of the human mitochondrial citrate carrier SLC25A1 lead to impaired citrate export required for lipid, dolichol, ubiquinone and sterol synthesis. Biochimica et Biophysica Acta (BBA) - Bioenergetics, 1859(1), 1–7. 10.1016/j.bbabio.2017.10.002

22. Meixner, E., Goldmann, U., Sedlyarov, V., Scorzoni, S., Rebsamen, M., Girardi, E., & Superti-Furga, G. (2020). A substrate-based ontology for human solute carriers. Molecular Systems Biology, 16(7). 10.15252/msb.20209652

23. Meyers, R. M., Bryan, J. G., McFarland, J. M., Weir, B. A., Sizemore, A. E., Xu, H., Dharia, N. V, Montgomery, P. G., Cowley, G. S., Pantel, S., Goodale, A., Lee, Y., Ali, L. D., Jiang, G., Lubonja, R., Harrington, W. F., Strickland, M., Wu, T., Hawes, D. C., … Tsherniak, A. (2017). Computational correction of copy number effect improves specificity of CRISPR– Cas9 essentiality screens in cancer cells. Nature Genetics, 49(12), 1779–1784. 10.1038/ng.3984

24. Mikkaichi, T., Suzuki, T., Onogawa, T., Tanemoto, M., Mizutamari, H., Okada, M., Chaki, T., Masuda, S., Tokui, T., Eto, N., Abe, M., Satoh, F., Unno, M., Hishinuma, T., Inui, K. I., Ito, S., Goto, J., & Abe, T. (2004). Isolation and characterization of a digoxin transporter and its rat homologue expressed in the kidney. Proc. Natl. Acad. Sci. U. S. A., 101(10), 3569– 3574. 10.1073/pnas.0304987101

25. Nimmanon, T., Ziliotto, S., Morris, S., Flanagan, L., & Taylor, K. M. (2017). Phosphorylation of zinc channel ZIP7 drives MAPK, PI3K and mTOR growth and proliferation signalling. Metallomics, 9(5), 471–481. 10.1039/c6mt00286b

26. Okada, Y. (2004). Ion Channels and Transporters Involved in Cell Volume Regulation and Sensor Mechanisms. Cell Biochemistry and Biophysics, 41(2), 233–258. 10.1385/cbb:41:2:233

27. Perez, M. F., & Sarkies, P. (2023). Histone methyltransferase activity affects metabolism in human cells independently of transcriptional regulation. PLoS Biology, 21(10 October). 10.1371/journal.pbio.3002354

28. Pizzagalli, M. D., Bensimon, A., & Superti-Furga, G. (2021). A guide to plasma membrane solute carrier proteins. In FEBS Journal (Vol. 288, Issue 9, pp. 2784–2835). John Wiley and Sons Inc. 10.1111/febs.15531

29. Ramamoorthy, S., Leibach, F. H., Mahesh, V. B., Han, H., Yang-Feng, T., Blakely, R. D., & Ganapathy, V. (1994). Functional characterization and chromosomal localization of a cloned taurine transporter from human placenta. Biochemical Journal, 300(3), 893–900. 10.1042/bj3000893

30. Rodriguez-Martinez, A., Ayala, R., Posma, J. M., Neves, A. L., Gauguier, D., Nicholson, J. K., & Dumas, M.-E. (2016). MetaboSignal: a network-based approach for topological analysis of metabotype regulationviametabolic and signaling pathways. Bioinformatics, 33(5), btw697. 10.1093/bioinformatics/btw697

31. Shoemaker, R. H. (2006). The NCI60 human tumour cell line anticancer drug screen. Nature Reviews Cancer, 6(10), 813–823. 10.1038/nrc1951

32. Skelton, M. R., Schaefer, T. L., Graham, D. L., deGrauw, T. J., Clark, J. F., Williams, M. T., & Vorhees, C. V. (2011). Creatine Transporter (CrT; Slc6a8) Knockout Mice as a Model of Human CrT Deficiency. PLoS ONE, 6(1), e16187. 10.1371/journal.pone.0016187

33. Song, W., Li, D., Tao, L., Luo, Q., & Chen, L. (2020). Solute carrier transporters: the metabolic gatekeepers of immune cells. Acta Pharmaceutica Sinica B, 10(1), 61–78. 10.1016/j.apsb.2019.12.006

34. Superti-Furga, G., Lackner, D., Wiedmer, T., Ingles-Prieto, A., Barbosa, B., Girardi, E., Goldmann, U., Gürtl, B., Klavins, K., Klimek, C., Lindinger, S., Liñeiro-Retes, E., Müller, A. C., Onstein, S., Redinger, G., Reil, D., Sedlyarov, V., Wolf, G., Crawford, M., … Steppan, C. M. (2020). The RESOLUTE consortium: unlocking SLC transporters for drug discovery. Nature Reviews Drug Discovery, 19(7), 429–430. 10.1038/d41573-020-00056-6

35. Szeri, F., Lundkvist, S., Donnelly, S., Engelke, U., Rhee, K., Williams, C. J., Sundberg, J. P., Wevers, R. A., Tomlinson, R. E., Jansen, R. S., & Wetering, K. (2020). The membrane protein ANKH is crucial for bone mechanical performance by mediating cellular export of citrate and ATP. PLOS Genetics, 16(7), e1008884–e1008884. 10.1371/journal.pgen.1008884

36. Taegtmeyer, H., & Ingwall, J. S. (2013). Creatine—A Dispensable Metabolite? Circulation Research, 112(6), 878–880. 10.1161/circresaha.113.300974

37. Tsherniak, A., Vazquez, F., Montgomery, P. G., Weir, B. A., Kryukov, G., Cowley, G. S., Gill, S., Harrington, W. F., Pantel, S., Krill-Burger, J. M., Meyers, R. M., Ali, L., Goodale, A., Lee, Y., Jiang, G., Hsiao, J., Gerath, W. F. J., Howell, S., Merkel, E., … Hahn, W. C. (2017). Defining a Cancer Dependency Map. Cell, 170(3), 564–576.e16. 10.1016/j.cell.2017.06.010

38. Wang, X., Ji, Y., Qi, J., Zhou, S., Wan, S., Fan, C., Gu, Z., An, P., Luo, Y., & Luo, J. (2023). Mitochondrial carrier 1 (MTCH1) governs ferroptosis by triggering the FoxO1-GPX4 axis-mediated retrograde signaling in cervical cancer cells. Cell Death and Disease, 14(8). 10.1038/s41419-023-06033-2

39. Wiegering, A., Matthes, N., Mühling, B., Koospal, M., Quenzer, A., Peter, S., Germer, C.-T., Linnebacher, M., & Otto, C. (2017). Reactivating p53 and Inducing Tumor Apoptosis (RITA) Enhances the Response of RITA-Sensitive Colorectal Cancer Cells to Chemotherapeutic Agents 5-Fluorouracil and Oxaliplatin. Neoplasia, 19(4), 301–309. 10.1016/j.neo.2017.01.007

40. Winter, G. E., Radic, B., Mayor-Ruiz, C., Blomen, V. A., Trefzer, C., Kandasamy, R. K., Huber, K. V. M., Gridling, M., Chen, D., Klampfl, T., Kralovics, R., Kubicek, S., Fernandez-Capetillo, O., Brummelkamp, T. R., & Superti-Furga, G. (2014). The solute carrier SLC35F2 enables YM155-mediated DNA damage toxicity. Nature Chemical Biology, 10(9), 768–773. 10.1038/nchembio.1590

41. Wyss, M., & Kaddurah-Daouk, R. (2000). Creatine and Creatinine Metabolism. Physiological Reviews, 80(3), 1107–1213. 10.1152/physrev.2000.80.3.1107

42. Zajac, M., Mukherjee, S., Anees, P., Oettinger, D., Henn, K., Srikumar, J., Zou, J., Saminathan, A., & Krishnan, Y. (2024). A mechanism of lysosomal calcium entry. Science Advances, 10(7). 10.1126/sciadv.adk2317

