## Supplemental Figure Legends for "Predicting substrates for orphan Solute Carrier Proteins using multi-omics datasets"

**Figure 1 | Correlating SLC transcript levels to substrate concentrations reveals known substrates of SLC.**

**A)** Schematic representation shows the principle of correlation analysis. SLC expression levels are hypothesised to correlate with the intracellular concentration of their corresponding substrates. Blue, SLC exporting substrate; red, SLC importing substrate. Figure created with elements from BioRender.

**B)** Scatter plot shows normalised Spearman’s ρ and adjusted *p*-value for 225 metabolites correlating with the expression level of SLC6A6 and SLC6A8 across 913 cell lines from CCLE. Key substrates are labelled.

**C)** Mutual concordance of SLC-metabolite correlation outcomes from different datasets, each processed with the pipeline specified in **Methods**. Nodes, datasets, edge, concordance parameter measured in Spearman’s ρ. CCLE2019 and NCI60, *p* = 4.30 x 10^-293^; CCLE2019 and CCL180, *p* = 1.50 x 10^-221^; NCI60 and CCL180, *p* = 2.38 x 10^-24^.

**Figure S1 | Rationale for threshold and score setting of the prediction algorithm.**

**A)** The selection of threshold of discovery for NCI60 dataset. 0.0 was selected.

**B)** The selection of threshold of discovery for CCLE2019 dataset. 0.0 was selected.

**C)** The selection of threshold of discovery for CCL180 dataset. 0.8 was selected.

**D)** The selection of threshold of discovery for CRISPR screen dataset. 0.077 was selected.

**E)** The selection of score reward for each discovery in CRISPR screen dataset. 10 was selected.

**F)** The selection of threshold of discovery for adjacent metabolites in CCL180 dataset. 0.1 was selected.

**G)** The selection of score reward (in proportion of score reward in correlation analysis) for each discovery of adjacent metabolites in CCL180n dataset. 0.9 was selected.

**H)** The selection of alpha for for drug screen dataset. 0.012 was selected.

**Figure S2 | The general similarity of normalised Spearman’s ρ between derivative across unit of adjacency.**

Boxplot shows the general similarity of normalised Spearman’s ρ between derivative across unit of adjacency. Red dashed line, Spearman’s ρ differences between adjacent derivatives and substrates are compared to non-adjacent controls across unit of adjacency. Grey boxes, Spearman’s ρ differences between non-adjacent controls are compared to another 100 non-adjacent controls across unit of adjacency.

**Figure S3 | A group of SLCs interacts with small molecule drug RITA**

**A)** Non-linear regression shows the dose-response of 9 SLCs interact to RITA.

**B)** Histograms show the frequency of number of cell lines that are shared across SLC presented in above. Green, cell lines marked with high SLC expression; Red, cell lines marked with low SLC expression.
