## Supplementary material for "Predicting substrates for orphan Solute Carrier Proteins using multi-omics datasets": Code for plotting results: Figure 1B.R

```
##### 0 load libraries #####
library(data.table)
library(tidyverse)

##### 1 Predictive score example plot SLC6A6 #####
SLC6A6 = fread("Desktop/Figure plots/Fig1/1B/SLC6A6 (example plot).csv")
ggplot() +
  geom_point(SLC6A6[order(ps, decreasing=TRUE)][3:nrow(SLC6A6)], mapping = aes(x = ps, y = -log10(padj)), color = "grey") +
  geom_point(SLC6A6[order(ps, decreasing=TRUE)][1:2], mapping = aes(x = ps, y = -log10(padj)), color = "#990000") +
  expand_limits(x = 10) +
  scale_x_continuous(breaks = seq(0,10,5)) +
  scale_y_continuous(limits = c(0,60), breaks = seq(0,60,20)) +
  theme(
    legend.position="none", plot.title = element_text(size=11),
    panel.grid.major = element_blank(), panel.grid.minor = element_blank(),
    panel.background = element_blank(), axis.line = element_line(colour = "black"))

##### 2 Predictive score example plot SLC6A8 #####
SLC6A8 = fread("Desktop/Figure plots/Fig1/1C/SLC6A8 (example plot).csv")
ggplot() +
  geom_point(SLC6A8[order(ps, decreasing=TRUE)][4:nrow(SLC6A8)], mapping = aes(x = ps, y = -log10(padj)), color = "grey") +
  geom_point(SLC6A8[order(ps, decreasing=TRUE)][1:3], mapping = aes(x = ps, y = -log10(padj)), color = "#990000") +
  expand_limits(x = 10) +
  scale_x_continuous(breaks = seq(0,10,5)) +
  scale_y_continuous(limits = c(0,120), breaks = seq(0,120,40)) +
  theme(
    legend.position="none", plot.title = element_text(size=11),
    panel.grid.major = element_blank(), panel.grid.minor = element_blank(),
    panel.background = element_blank(), axis.line = element_line(colour = "black"))
```
