## Supplementary material for "Predicting substrates for orphan Solute Carrier Proteins using multi-omics datasets": Code for plotting results: Figure 1C.R

```
##### 0 Load libraries #####
library(data.table)
library(tidyverse)

##### 1 Read files #####
et = fread("Desktop/Figure plots/Fig1/1C/Evidence_Table_in_rho.csv")

##### 2 CCLE2019 and NCI60 #####
et_1 <- et[rowSums(is.na(et[,3:4])) == 0][, !5:6]
length(unique(et_1$Formula))
length(unique(et_1$SYMBOL))
cor.test(et_1$CCLE2019, et_1$NCI60, method = "spearman", exact = F)$p.value
cor.test(et_1$CCLE2019, et_1$NCI60, method = "spearman", exact = F)$estimate

##### 3 CCLE2019 and CCL180 #####
et_2 = et[rowSums(is.na(et[,4:5])) == 0][, !c(3,6)]
length(unique(et_2$Formula))
length(unique(et_2$SYMBOL))
cor.test(et_2$CCLE2019, et_2$CCL180, method = "spearman", exact = F)$p.value
cor.test(et_2$CCLE2019, et_2$CCL180, method = "spearman", exact = F)$estimate

##### 4 NCI60 and CCL180 #####
et_3 = et[rowSums(is.na(et[,c(3,5)])) == 0][, !c(4,6)]
length(unique(et_3$Formula))
length(unique(et_3$SYMBOL))
cor.test(et_3$NCI60, et_3$CCL180, method = "spearman", exact = F)$p.value
cor.test(et_3$NCI60, et_3$CCL180, method = "spearman", exact = F)$estimate
```
