## Supplementary material for "Predicting substrates for orphan Solute Carrier Proteins using multi-omics datasets": Code for plotting results: Figure 1D.R

```
##### 0 load libraries #####
library(data.table)
library(tidyverse)
library(hrbrthemes)
library(viridis)

##### 1 read files #####
score = fread("Desktop/Figure plots/Fig1/1D/Correlation analysis benchmark report (new).csv")

##### 2 violin plot #####
score |>
  filter(Data == "CCLE2019" & Type == "Simulated") |>
  ggplot(aes(x = Data, y = Mean, fill = Data)) + 
  geom_violin(alpha = 1) +
  geom_point(position = position_jitter(seed = 1, width = 0.1)) +
  scale_fill_manual(values=c("#8ABFBD")) +
  ylim(c(-0.15, 0.2)) +
  theme_ipsum() +
  theme(
    legend.position="none",
    plot.title = element_text(size=11)
  ) +
  geom_hline(aes(yintercept=score[Data == "CCLE2019" & Type == "Known", Mean]), colour="#990000", linetype="dashed") +
  coord_flip() +
  theme(panel.grid.major = element_blank(), panel.grid.minor = element_blank(),
        panel.background = element_blank(), axis.line = element_line(colour = "black"),
        axis.text.y = element_blank())

shapiro.test(score[Data == "CCLE2019" & Type == "Simulated", Mean]) # normal distribution
t.test(score[Data == "CCLE2019" & Type == "Simulated", Mean], mu = score[Data == "CCLE2019" & Type == "Known", Mean])$p.value

score |>
  filter(Data == "NCI60" & Type == "Simulated") |>
  ggplot(aes(x = Data, y = Mean, fill = Data)) + 
  geom_violin(alpha = 1) +
  geom_point(position = position_jitter(seed = 1, width = 0.1)) +
  scale_fill_manual(values=c("#FDF18A")) +
  #ylim(c(-0.15, 0.2)) +
  theme_ipsum() +
  theme(
    legend.position="none",
    plot.title = element_text(size=11)
  ) +
  geom_hline(aes(yintercept=score[Data == "NCI60" & Type == "Known", Mean]), colour="#990000", linetype="dashed") +
  coord_flip() +
  theme(panel.grid.major = element_blank(), panel.grid.minor = element_blank(),
        panel.background = element_blank(), axis.line = element_line(colour = "black"),
        axis.text.y = element_blank())

shapiro.test(score[Data == "NCI60" & Type == "Simulated", Mean]) # normal distribution
t.test(score[Data == "NCI60" & Type == "Simulated", Mean], mu = score[Data == "NCI60" & Type == "Known", Mean])$p.value

score |>
  filter(Data == "CCL180" & Type == "Simulated") |>
  ggplot(aes(x = Data, y = Mean, fill = Data)) + 
  geom_violin(alpha = 1) +
  geom_point(position = position_jitter(seed = 1, width = 0.1)) +
  scale_fill_manual(values=c("#987B9F")) +
  #ylim(c(-0.15, 0.2)) +
  theme_ipsum() +
  theme(
    legend.position="none",
    plot.title = element_text(size=11)
  ) +
  geom_hline(aes(yintercept=score[Data == "CCL180" & Type == "Known", Mean]), colour="#990000", linetype="dashed") +
  coord_flip() +
  theme(panel.grid.major = element_blank(), panel.grid.minor = element_blank(),
        panel.background = element_blank(), axis.line = element_line(colour = "black"),
        axis.text.y = element_blank())

shapiro.test(score[Data == "CCL180" & Type == "Simulated", Mean]) # normal distribution
t.test(score[Data == "CCL180" & Type == "Simulated", Mean], mu = score[Data == "CCL180" & Type == "Known", Mean])$p.value


# Most basic violin chart (not in use)
p = ggplot(score_random_normalised, aes(x=source, y=score, fill=source)) + # fill=name allow to automatically dedicate a color for each group
  geom_boxplot() +
  ylim(c(-0.15, 0.2)) +
  scale_fill_viridis(discrete = TRUE, alpha=0.6, option="D") +
  theme_ipsum() +
  theme(
    legend.position="none",
    plot.title = element_text(size=11)
  ) +
  geom_hline(aes(yintercept=0.2), colour="#990000", linetype="dashed") +
  coord_flip() +
  theme(panel.grid.major = element_blank(), panel.grid.minor = element_blank(),
        panel.background = element_blank(), axis.line = element_line(colour = "black"))

p
```
