## Supplementary material for "Predicting substrates for orphan Solute Carrier Proteins using multi-omics datasets": Code for plotting results: Figure 1E.R

```
##### 0 load library #####
library(data.table)
library(tidyverse)
library(hrbrthemes)
library(viridis)

##### 1 read files #####
ScoreResult = fread("Desktop/Figure plots/Fig1/1E/Factor_A_ScoreResult.csv")

##### 2 plot #####
ggplot(ScoreResult[Class == "random"], aes(x = Class, y = Score)) +
  geom_boxplot(fill = "grey") +
  scale_fill_viridis(discrete = TRUE, alpha=0.6, option="C") +
  #ylim(c(10,18)) +
  theme(
    legend.position="none",
    plot.title = element_text(size=11)
  ) +
  geom_hline(aes(yintercept=ScoreResult[Class == "true", Score]), colour="#990000", linetype="dashed") +
  theme(panel.grid.major = element_blank(), panel.grid.minor = element_blank(),
        panel.background = element_blank(), axis.line = element_line(colour = "black")) +
  scale_y_continuous(breaks=seq(6,16,2)) +
  expand_limits(y = 16)

shapiro.test(ScoreResult[Class == "random", Score]) # normally distributed
t.test(ScoreResult[Class == "random", Score], mu = ScoreResult[Class == "true", Score])$p.value
```
