## Supplementary material for "Predicting substrates for orphan Solute Carrier Proteins using multi-omics datasets": Code for plotting results: Figure 3C.R

```
##### 0 load library #####
library(data.table)
library(tidyverse)
library(hrbrthemes)
library(viridis)

##### 1 read files #####
FractionRecovery = fread("Desktop/Figure plots/Fig3/3C/CRISPRDEP_FractionRecoveryAcrossAlpha.csv")

##### 2 plot #####
ggplot() +
  geom_violin(FractionRecovery[Type == "Simulated"], 
              mapping = aes(x = AlphaCutoff, y = Fraction_recovery, group = AlphaCutoff),
              color = "grey", fill = "grey") +
  geom_line(FractionRecovery[Type == "Known"],
            mapping = aes(x= AlphaCutoff, y= Fraction_recovery),
            color = "#990000",
            linetype="dashed") + 
  theme(
    legend.position="none", plot.title = element_text(size=11),
    panel.grid.major = element_blank(), panel.grid.minor = element_blank(),
    panel.background = element_blank(), axis.line = element_line(colour = "black")) +
  scale_x_continuous(breaks=seq(0,0.1,0.02)) +
  scale_y_continuous(breaks = seq(0,0.06,0.01))

ggplot(FractionRecovery, aes(x = Type, y = Fraction_recovery, fill = Type)) +
  geom_violin(width=1.4) +
  geom_boxplot(width=0.05, color="black", fill = "white") +
  scale_fill_manual(values=c("#990000", "grey")) +
  theme(
    legend.position="none", plot.title = element_text(size=11),
    panel.grid.major = element_blank(), panel.grid.minor = element_blank(),
    panel.background = element_blank(), axis.line = element_line(colour = "black"))

wilcox.test(FractionRecovery[Type == "Known", Fraction_recovery], FractionRecovery[Type == "Simulated", Fraction_recovery], alternative = "greater")$p.value
```
