## Supplementary material for "Predicting substrates for orphan Solute Carrier Proteins using multi-omics datasets": Code for plotting results: Figure 2C.R

```
library(data.table)
library(tidyverse)

dCS = fread("~/Desktop/Figure plots/Fig2/2C/dCS.csv")

ggplot(dCS, aes(x = Factor, y = difference.averageCS, fill = Factor)) +
  geom_boxplot() +
  scale_fill_manual(values=c("#3D5C6F", "#F9AE78")) +
  expand_limits(y = 6) +
  scale_y_continuous(breaks = seq(4,6,1)) +
  theme(
    legend.position="none", plot.title = element_text(size=11),
    panel.grid.major = element_blank(), panel.grid.minor = element_blank(),
    panel.background = element_blank(), axis.line = element_line(colour = "black"))

shapiro.test(dCS[Factor == "A", difference.averageCS])
shapiro.test(dCS[Factor == "AB", difference.averageCS])

t.test(dCS[Factor == "A", difference.averageCS], dCS[Factor == "AB", difference.averageCS], alternative = "l")
mean((ConfAB_dcs - ConfA_dcs)/ConfA_dcs)
```
