## Supplementary material for "Predicting substrates for orphan Solute Carrier Proteins using multi-omics datasets": Code for plotting results: Figure 3B.R

```
##### 0 load libraries #####
library(data.table)
library(tidyverse)

##### 1 read files #####
step_assembly = fread("Desktop/Figure plots/Fig4/4B/proximal and distant (rho).csv")

##### 2 plot #####
step_simulated = step_assembly[class == "simulated"]
step_known = step_assembly[class == "known"]

g = ggplot() +
  # box plot of fecs
  geom_boxplot(data = step_simulated, 
               aes(x = Steps, y= -log10(WT.pvalue), group = Steps), fill = "grey") +
  # points of ecs
  geom_line(data = step_known,
            aes(x= Steps, y= -log10(WT.pvalue)),
            color = "#990000",
            linetype="dashed") + 
  theme(
    legend.position="none",
    plot.title = element_text(size=11)
  ) +
  theme(panel.grid.major = element_blank(), panel.grid.minor = element_blank(),
        panel.background = element_blank(), axis.line = element_line(colour = "black")) +
  scale_x_continuous(breaks=seq(0,10,2))
g
```
