## Supplementary material for "Predicting substrates for orphan Solute Carrier Proteins using multi-omics datasets": Code for plotting results: Figure 3C-D.R

```
library(data.table)
library(tidyverse)

# Figure 3C
dCS = fread("~/Desktop/Figure plots/Fig3/3C-D/ConfidenceScoreDifference.csv")
ggplot(dCS, aes(x = Factor, y = difference.averageCS, fill = Factor)) +
  geom_boxplot() +
  scale_fill_manual(values=c("#3D5C6F", "#F9AE78","#E47159")) +
  expand_limits(y = 12) +
  scale_y_continuous(breaks = seq(4,12,4)) +
  theme(
    legend.position="none", plot.title = element_text(size=11),
    panel.grid.major = element_blank(), panel.grid.minor = element_blank(),
    panel.background = element_blank(), axis.line = element_line(colour = "black"))

shapiro.test(dCS[Factor == "A", difference.averageCS])
shapiro.test(dCS[Factor == "AB", difference.averageCS])
shapiro.test(dCS[Factor == "ABC", difference.averageCS])

t.test(dCS[Factor == "AB", difference.averageCS], dCS[Factor == "ABC", difference.averageCS], alternative = "l")$p.value

# Figure 3D
FractionDifference = fread("~/Desktop/Figure plots/Fig3/3C-D/FractionDifference.csv")
ggplot(FractionDifference, aes(x = X, y = Q, group = interaction(X, Fct), fill = Fct, color = Fct)) +
  geom_violin(alpha = 0.2) +
  scale_color_manual(values=c("#3D5C6F", "#F9AE78","#E47159")) +
  scale_fill_manual(values=c("#3D5C6F", "#F9AE78","#E47159")) +
  theme(
    legend.position="none", plot.title = element_text(size=11),
    panel.grid.major = element_blank(), panel.grid.minor = element_blank(),
    panel.background = element_blank(), axis.line = element_line(colour = "black")) +
  scale_y_continuous(breaks = seq(0, 0.2, 0.04))

wilcox.test(FractionDifference[Fct == "AB", Q], FractionDifference[Fct == "ABC", Q], alternative = "l")$p.value
```
