## Supplementary material for "Predicting substrates for orphan Solute Carrier Proteins using multi-omics datasets": Code for plotting results: Figure 4A.R

```
library(data.table)
library(tidyverse)
library(hrbrthemes)
library(viridis)

median_benchmark = fread("Desktop/Figure plots/Fig4/4A/MedianBenchmark.csv")

ggplot(median_benchmark[Class == "simulated"], aes(x = Class, y = Median)) +
  geom_boxplot(fill = "grey") +
  scale_fill_viridis(discrete = TRUE, alpha=0.6, option="C") +
  #ylim(c()) +
  theme(
    legend.position="none",
    plot.title = element_text(size=11)
  ) +
  geom_hline(aes(yintercept=median_benchmark[Class == "true", Median]), colour="#990000", linetype="dashed") +
  theme(panel.grid.major = element_blank(), panel.grid.minor = element_blank(),
        panel.background = element_blank(), axis.line = element_line(colour = "black"))
#scale_y_continuous(limits = c(150, 400), breaks=seq(150, 400,50)) 
#expand_limits()

shapiro.test(median_benchmark[Class == "simulated", Median]) # not normally distributed
wilcox.test(median_benchmark[Class == "simulated", Median], mu = median_benchmark[Class == "true", Median], alternative = "g")$p.value
```
