## Supplementary material for "Predicting substrates for orphan Solute Carrier Proteins using multi-omics datasets": Code for plotting results: Figure 4B.R

```
library(data.table)
library(tidyverse)
library(hrbrthemes)
library(viridis)

Sweep = fread("Desktop/Figure plots/Fig4/4B/Sweep.csv")

ggplot() +
  geom_violin(Sweep, mapping = aes(x = X, y = Q, group = X, alpha = 0.2), color = "grey", fill = "grey") +
  geom_line(data.table(X = 1:500, Q = Sweep[, median(Q), by = X][, V1]), mapping = aes(x = X, y = Q)) +
  theme(
    legend.position="none", plot.title = element_text(size=11),
    panel.grid.major = element_blank(), panel.grid.minor = element_blank(),
    panel.background = element_blank(), axis.line = element_line(colour = "black"))
```
