## Supplementary material for "Predicting substrates for orphan Solute Carrier Proteins using multi-omics datasets": Code for plotting results: Figure 4C.R

```
library(data.table)
library(tidyverse)

plot_table = fread("Desktop/Figure plots/Fig4/4C/ORA_Plot.csv")

ggplot(plot_table, aes(x = Rank, y = Number)) +
  geom_point() +
  theme(
    legend.position="none", plot.title = element_text(size=11),
    panel.grid.major = element_blank(), panel.grid.minor = element_blank(),
    panel.background = element_blank(), axis.line = element_line(colour = "black"))
```
