## Supplementary material for "Predicting substrates for orphan Solute Carrier Proteins using multi-omics datasets": Code for plotting results: Figure 5B.R

```
##### 0 load libraries #####
library(data.table)
library(tidyverse)

##### 1 read files #####
exampleDT = fread("Desktop/Figure plots/Fig5/5B/SLC35F2 vs YM155.csv") # positive example
exampleDT = fread("Desktop/Figure plots/Fig5/5B/SLC19A2 vs Thiamine.csv") # negative example
exampleDT = fread("Desktop/Figure plots/Fig5/5B/PTCHD4 vs idasanutlin.csv") # prediction example

##### 2 draw plots #####
ggplot(exampleDT, aes(x=log10(dose), y=viability, fill =expression_type, color = expression_type)) + 
  scale_fill_manual(values=c("#63BBB0", "#AC667E")) +
  scale_color_manual(values=c("#63BBB0", "#AC667E")) +
  geom_point(position = position_dodge(0.2), alpha = 0.5) + 
  geom_smooth(method = 'loess') + 
  scale_y_continuous(limits = c(-7.5, 2.5), breaks = seq(-7.5, 2.5, 2.5)) +
  theme(
    legend.position="none", plot.title = element_text(size=11),
    panel.grid.major = element_blank(), panel.grid.minor = element_blank(),
    panel.background = element_blank(), axis.line = element_line(colour = "black"))
```
