## Supplementary material for "Predicting substrates for orphan Solute Carrier Proteins using multi-omics datasets": Code for plotting results: Figure 5C.R

```
##### 0 load libraries #####
library(data.table)
library(ggplot2)
library(hrbrthemes)
library(viridis)

##### 1 read files #####
benchmark = fread("Desktop/Figure plots/Fig5/5C/t-test model prediction benchmark.csv")

##### 2 draw plots #####
ggplot(benchmark[Status == "random"], aes(x = Status, y=log.p)) +
  geom_boxplot(fill = "grey") +
  scale_fill_viridis(discrete = TRUE, alpha=0.6, option="C") +
  expand_limits(y = 20) +
  scale_y_continuous(breaks = seq(5, 15, 5)) +
  theme_ipsum() +
  theme(
    legend.position="none",
    plot.title = element_text(size=11)
  ) +
  geom_hline(aes(yintercept=benchmark[Status == "true", log.p]), colour="#990000", linetype="dashed") +
  theme(panel.grid.major = element_blank(), panel.grid.minor = element_blank(),
        panel.background = element_blank(), axis.line = element_line(colour = "black")) +
  coord_flip()

shapiro.test(benchmark[Status == "random", log.p]) # not normal distribution
wilcox.test(benchmark[Status == "random", log.p], mu = benchmark[Status == "true", log.p], alternative = "l")$p.value

ggplot(benchmark[Status == "random"], aes(x = Status, y=mean.diff)) +
  geom_boxplot(fill = "grey") +
  scale_fill_viridis(discrete = TRUE, alpha=0.6, option="C") +
  expand_limits(y = 0.15) +
  scale_y_continuous(breaks = seq(0.05, 0.10, 0.05)) +
  theme_ipsum() +
  theme(
    legend.position="none",
    plot.title = element_text(size=11)
  ) +
  geom_hline(aes(yintercept=benchmark[Status == "true", mean.diff]), colour="#990000", linetype="dashed") +
  theme(panel.grid.major = element_blank(), panel.grid.minor = element_blank(),
        panel.background = element_blank(), axis.line = element_line(colour = "black")) +
  coord_flip()

shapiro.test(benchmark[Status == "random", mean.diff]) # not normal distribution
t.test(benchmark[Status == "random", mean.diff], mu = benchmark[Status == "true", mean.diff], alternative = "l")$p.value

##### 3 read files #####
benchmark = fread("Desktop/Figure plots/Fig5/5C/check dose effect benchmark.csv")

##### 4 draw plots #####
ggplot(benchmark[Status == "random"], aes(x = Status, y=log.p.mean)) +
  geom_boxplot(fill = "grey") +
  scale_fill_viridis(discrete = TRUE, alpha=0.6, option="C") +
  expand_limits(y = 6) +
  scale_y_continuous(breaks = seq(1, 4, 1)) +
  theme_ipsum() +
  theme(
    legend.position="none",
    plot.title = element_text(size=11)
  ) +
  geom_hline(aes(yintercept=benchmark[Status == "true", log.p.mean]), colour="#990000", linetype="dashed") +
  theme(panel.grid.major = element_blank(), panel.grid.minor = element_blank(),
        panel.background = element_blank(), axis.line = element_line(colour = "black")) +
  coord_flip()

shapiro.test(benchmark[Status == "random", log.p.mean]) # not normal distribution
wilcox.test(benchmark[Status == "random", log.p.mean], mu = benchmark[Status == "true", log.p.mean], alternative = "l")$p.value

ggplot(benchmark[Status == "random"], aes(x = Status, y=rho.mean)) +
  geom_boxplot(fill = "grey") +
  scale_fill_viridis(discrete = TRUE, alpha=0.6, option="C") +
  expand_limits(y = 1) +
  scale_y_continuous(breaks = seq(0.4, 0.6, 0.1)) +
  theme_ipsum() +
  theme(
    legend.position="none",
    plot.title = element_text(size=11)
  ) +
  geom_hline(aes(yintercept=benchmark[Status == "true", rho.mean]), colour="#990000", linetype="dashed") +
  theme(panel.grid.major = element_blank(), panel.grid.minor = element_blank(),
        panel.background = element_blank(), axis.line = element_line(colour = "black")) +
  coord_flip()

shapiro.test(benchmark[Status == "random", rho.mean]) # not normal distribution
t.test(benchmark[Status == "random", rho.mean], mu = benchmark[Status == "true", rho.mean], alternative = "l")$p.value
```
