## Supplementary material for "Predicting substrates for orphan Solute Carrier Proteins using multi-omics datasets": Code for plotting results: Figure S1A-C.R

```
library(data.table)
library(tidyverse)

Sweep_Ax = fread("Desktop/Figure plots/FigS1/S1A-C/Sweep_factorAx.csv")
Sweep_Ay = fread("Desktop/Figure plots/FigS1/S1A-C/Sweep_factorAy.csv")
Sweep_Az = fread("Desktop/Figure plots/FigS1/S1A-C/Sweep_factorAz.csv")

ggplot(Sweep_Ax, aes(x = Th, y = MaxTP)) +
  geom_point() +
  scale_y_continuous(limits = c(0.06, 0.11), breaks = seq(0.06, 0.11, 0.01)) +
  scale_x_continuous(breaks = seq(0,1,0.1)) +
  theme(panel.grid.major = element_blank(), panel.grid.minor = element_blank(),
        panel.background = element_blank(), axis.line = element_line(colour = "black"))

ggplot(Sweep_Ay, aes(x = Th, y = MaxTP)) +
  geom_point() +
  scale_y_continuous(limits = c(0.04, 0.11), breaks = seq(0.04, 0.11, 0.01)) +
  scale_x_continuous(breaks = seq(0,1,0.1)) +
  theme(panel.grid.major = element_blank(), panel.grid.minor = element_blank(),
        panel.background = element_blank(), axis.line = element_line(colour = "black"))

ggplot(Sweep_Az, aes(x = Th, y = MaxTP)) +
  geom_point() +
  scale_y_continuous(limits = c(0.033, 0.039), breaks = seq(0.03, 0.04, 0.001)) +
  scale_x_continuous(breaks = seq(0,1,0.1)) +
  theme(panel.grid.major = element_blank(), panel.grid.minor = element_blank(),
        panel.background = element_blank(), axis.line = element_line(colour = "black"))
```
