## Supplementary material for "Predicting substrates for orphan Solute Carrier Proteins using multi-omics datasets": Code for plotting results: Figure S1D-E.R

```
##### 0 load libraries #####
library(data.table)
library(tidyverse)

Sweep_factorABalpha = fread("Desktop/Figure plots/FigS1/S1D/Sweep_factorABalpha.csv")

ggplot(Sweep_factorABalpha, aes(x = AlphaCutoff, y = Fraction_difference)) +
  geom_point() +
  scale_y_continuous(limits = c(0.02, 0.05)) +
  scale_x_continuous(breaks = seq(0,0.1,0.01)) +
  theme(panel.grid.major = element_blank(), panel.grid.minor = element_blank(),
        panel.background = element_blank(), axis.line = element_line(colour = "black"))

Sweep_factorABscore = fread("Desktop/Figure plots/FigS1/S1D-E/Sweep_factorABscore.csv")

ggplot(Sweep_factorABscore, aes(x = Po, y = MaxQ)) +
  geom_point() +
  #scale_y_continuous(limits = c(0.02, 0.05)) +
  #scale_x_continuous(breaks = seq(0,0.1,0.01)) +
  theme(panel.grid.major = element_blank(), panel.grid.minor = element_blank(),
        panel.background = element_blank(), axis.line = element_line(colour = "black"))
```
