## Supplementary material for "Predicting substrates for orphan Solute Carrier Proteins using multi-omics datasets": Code for plotting results: Figure S1F-H.R

```
##### 0 load libraries #####
library(data.table)
library(tidyverse)

Sweep_ABCz = fread("Desktop/Figure plots/FigS1/S1F-G/Sweep_factorABCz.csv")
ggplot(Sweep_ABCz, aes(x = Th, y = MaxQ)) +
  geom_point() +
  scale_y_continuous(limits = c(0.145, 0.147), breaks = seq(0.145, 0.147, 0.001)) +
  scale_x_continuous(breaks = seq(0,1,0.1)) +
  theme(panel.grid.major = element_blank(), panel.grid.minor = element_blank(),
        panel.background = element_blank(), axis.line = element_line(colour = "black"))


Sweep_ABCp = fread("Desktop/Figure plots/FigS1/S1F-G/Sweep_factorABCp.csv")
ggplot(Sweep_ABCp, aes(x = Po, y = MaxQ)) +
  geom_point() +
  scale_y_continuous(limits = c(0.12, 0.15), breaks = seq(0.12, 0.15, 0.01)) +
  scale_x_continuous(breaks = seq(0,1,0.1)) +
  theme(panel.grid.major = element_blank(), panel.grid.minor = element_blank(),
        panel.background = element_blank(), axis.line = element_line(colour = "black"))
```
