## Supplementary material for "Predicting substrates for orphan Solute Carrier Proteins using multi-omics datasets": Code for plotting results: Figure S1H.R

```
library(data.table)
library(tidyverse)

Sweep_drugAlpha = fread("Desktop/Figure plots/FigS1/S1H/Sweep_drugAlpha.csv")

ggplot(Sweep_drugAlpha, aes(x = alpha, y = discovery)) +
  geom_point() +
  scale_y_continuous(limits = c(0.09, 0.18), breaks = seq(0.09, 0.17,0.02)) +
  scale_x_continuous(breaks = seq(0,0.05,0.01)) +
  theme(panel.grid.major = element_blank(), panel.grid.minor = element_blank(),
        panel.background = element_blank(), axis.line = element_line(colour = "black"))
```
