## Supplementary material for "Predicting substrates for orphan Solute Carrier Proteins using multi-omics datasets": Code for generating results: CCL180 TvsM.R

```
##### 0 load libraries #####
library(reshape2)
library(data.table)
library(dplyr)

##### 1 read files #####
SLC_transcript_counts <- fread("03 Results/01 Correlation Analysis/CCLE2019/CCLE2019 SLC normalised transcript counts.csv")
CCL180_ion_intensity <- fread("01 Raw Data/01 Correlation Analysis/04 CCL180/meta_Intensities.csv")
CCL180_ion_annotations <- fread("01 Raw Data/01 Correlation Analysis/04 CCL180/TopIds.csv")

##### 2 format transcript counts and metabolomics #####
# construct metabolomics with ion intensity and ion annotation
CCL180_ion_intensity[,V3 := CCL180_ion_annotations$`Top annotation name`]
colnames(CCL180_ion_intensity) <- gsub(".*\\/ ", "", colnames(CCL180_ion_intensity))
CCL180_metabolomics <- data.table(matrix(ncol = 180, nrow = 1809))
colnames(CCL180_metabolomics) <- unique(colnames(CCL180_ion_intensity)[4:ncol(CCL180_ion_intensity)])
CCL180_metabolomics <- data.table(Metabolite = CCL180_ion_intensity$V3, CCL180_metabolomics)

for (idx in 2:ncol(CCL180_metabolomics)){
  cell_line_name <- colnames(CCL180_metabolomics)[idx]
  sub_ion_intensity <- CCL180_ion_intensity[, which(colnames(CCL180_ion_intensity) == cell_line_name), with = FALSE]
  CCL180_metabolomics[,(idx) := rowMeans(sub_ion_intensity)]
  print(idx)
}

# match cell line in metabolomics and transcript counts
colnames(SLC_transcript_counts) = gsub("_.*", "", colnames(SLC_transcript_counts))
# trim
CCL180_cell_lines <- colnames(CCL180_metabolomics)[2:ncol(CCL180_metabolomics)]
CCLE2019_cell_lines <- colnames(SLC_transcript_counts)[2:ncol(SLC_transcript_counts)]
overlap <- intersect(CCL180_cell_lines, CCLE2019_cell_lines) # 151 cell name overlap
CCL180_cell_lines [!CCL180_cell_lines %in% overlap] # 29 CCL180 cell lines not sharing with CCLE2019
CCLE2019_cell_lines [!CCLE2019_cell_lines %in% overlap] # 762 CCLE2019 cell lines not sharing with CCL180
# the reason why only 151, not 153 cell lines claimed in original paper, is that the CCLE2019 transcript counts removed cell lines not in CCLE2019 metabolomics
# and the 2 cell lines removed are included in CCL180. For the sake of consistency, they will not be included in this study

# match
CCL180_metabolomics <- CCL180_metabolomics[,c(TRUE, CCL180_cell_lines %in% overlap), with = FALSE]
SLC_transcript_counts <- SLC_transcript_counts[,c(TRUE, CCLE2019_cell_lines %in% overlap), with = FALSE]
CCL180_cell_lines <- colnames(CCL180_metabolomics)[2:ncol(CCL180_metabolomics)] # updated CCL180 cell lines
CCLE2019_cell_lines <- colnames(SLC_transcript_counts)[2:ncol(SLC_transcript_counts)] # updated CCLE2019 cell lines
reorder_idx = match(CCLE2019_cell_lines, CCL180_cell_lines)
CCL180_metabolomics = CCL180_metabolomics[,c(1, reorder_idx+1), with = FALSE]
all(CCLE2019_cell_lines == colnames(CCL180_metabolomics)[2:ncol(CCL180_metabolomics)]) # the cell line order is aligned between transcript counts and metabolomics

# adjust duplicated terms
index_of_change <- c(650, 890, 1137, 1334, 1402, 1468, 1554, 1670)
metabolite_of_change <- c("5a-Androst-3-en-17-one.2", "Acetaminophen glucuronide.2", "Dihydrofukinolide.2", "Acuminoside.2",
                          "alpha-Tocopherol acetate.2", "Theasapogenol A.2", "Musabalbisiane C.2", "Opiorphin.2")
CCL180_metabolomics[index_of_change, Metabolite := metabolite_of_change]

# change ADP-ribose 1""-2"" cyclic phosphate
CCL180_metabolomics[1609, Metabolite := "ADP-ribose 1-2 cyclic phosphate"]

fwrite(SLC_transcript_counts, "03 Results/01 Correlation Analysis/CCL180/CCL180 SLC normalised transcript counts.csv")
fwrite(CCL180_metabolomics, "03 Results/01 Correlation Analysis/CCL180/CCL180 metabolomics.csv")

# cleaning after step 2
rm(CCL180_cell_lines)
rm(CCLE2019_cell_lines)
rm(cell_line_name)
rm(idx)
rm(overlap)
rm(reorder_idx)
rm(index_of_change)
rm(metabolite_of_change)

##### 3 CCL180 SLC TvsM #####
SLC_transcript_counts <- fread("03 Results/01 Correlation Analysis/CCL180/CCL180 SLC normalised transcript counts.csv")
CCL180_metabolomics <- fread("03 Results/01 Correlation Analysis/CCL180/CCL180 metabolomics.csv")

# create empty matrix to store statistical output
SLC_OutputCor <- matrix(0, nrow=nrow(SLC_transcript_counts),ncol=nrow(CCL180_metabolomics))
SLC_OutputP <- matrix(1, nrow=nrow(SLC_transcript_counts),ncol=nrow(CCL180_metabolomics))

for(i in 1:nrow(SLC_transcript_counts)){
  for(j in 1:nrow(CCL180_metabolomics)){
    CorIn <- cor.test(as.numeric(SLC_transcript_counts[i,2:ncol(SLC_transcript_counts)]),
                      as.numeric(CCL180_metabolomics[j,2:ncol(CCL180_metabolomics)]), method="spearman", exact = FALSE)
    SLC_OutputCor[i,j] <- CorIn$estimate
    SLC_OutputP[i,j] <- CorIn$p.value
  }
  print(i)
}

# FDR for multiple correlation analysis
SLC_OutputPadj <- matrix(1, nrow=nrow(SLC_transcript_counts),ncol=nrow(CCL180_metabolomics))
for (k in 1:nrow(SLC_OutputP)) {
  SLC_OutputPadj[k,] <- p.adjust(SLC_OutputP[k,], method = "BH", n = length(SLC_OutputP[k,]))
}

SLC_OutputPadj <- as.data.frame(SLC_OutputPadj)
SLC_OutputCor <- as.data.frame(SLC_OutputCor)
colnames(SLC_OutputPadj) <- CCL180_metabolomics$Metabolite #set up rowname and colname of adjusted p value
colnames(SLC_OutputCor) <- CCL180_metabolomics$Metabolite #set up rowname and colname of correlation value
SLC_OutputPadj$SYMBOL <- SLC_transcript_counts$SYMBOL
SLC_OutputCor$SYMBOL <- SLC_transcript_counts$SYMBOL
SLC_OutputPadj <- SLC_OutputPadj %>% dplyr::select(SYMBOL, everything())
SLC_OutputCor <- SLC_OutputCor %>% dplyr::select(SYMBOL, everything())

# melt correlation matrices
mSLC_OutputPadj <- melt(SLC_OutputPadj, id.vars = "SYMBOL")
mSLC_OutputCor <- melt(SLC_OutputCor, id.vars = "SYMBOL")

# check if melted data frame has the same arrangement
all(mSLC_OutputPadj$SYMBOL == mSLC_OutputCor$SYMBOL) # TRUE
all(mSLC_OutputPadj$variable == mSLC_OutputCor$variable) # TRUE

SLC_interaction = mSLC_OutputPadj
colnames(SLC_interaction) = c("SYMBOL", "METABOLITE", "padj")
SLC_interaction$rho = mSLC_OutputCor$value

fwrite(SLC_interaction, "03 Results/01 Correlation Analysis/CCL180/CCL180 TvsM.csv")
```
