## Supplementary material for "Predicting substrates for orphan Solute Carrier Proteins using multi-omics datasets": Code for generating results: CCLE2019 TvsM.R

```
##### 0 load libraries #####
library(ggplot2)
library(CePa)
library(DESeq2)
library(data.table)
library(biomaRt)
library(dplyr)
library(reshape2)

##### 1 read files #####
transcript_counts <- read.gct(file="01 Raw Data/01 Correlation Analysis/01 CCLE2019/CCLE_RNAseq_genes_counts_20180929.gct")
CCLE2019_metabolomics <- fread("01 Raw Data/01 Correlation Analysis/01 CCLE2019/CCLE_metabolomics_20190502.csv")
SLC_annotation_2024 <- fread("01 Raw Data/00 SLC annotation 2024/SLC_transcript_20240321.csv", select = c(1:6))

##### 2 normalise transcript counts with MRN #####
# create dds object in DESeq2
colData <- data.frame(condition= colnames(transcript_counts), row.names = colnames(transcript_counts))
dds <- DESeqDataSetFromMatrix(countData = transcript_counts,
                             colData = colData,
                             design = ~ condition)
# quality control: filtering rows with less than 10 reads
keep <- rowSums(counts(dds)) >= 10
dds <- dds[keep,]
# normalisation with MRN
dds <- estimateSizeFactors(dds)
MRN_transcript_counts <- counts(dds, normalized = TRUE)

# cleaning after step 2
rm(colData)
rm(dds)
rm(keep)

##### 3 format normalised transcript counts and metabolomics #####
# edit cell line name with wrong format
MRN_transcript_counts <- as.data.frame(MRN_transcript_counts)
colnames(MRN_transcript_counts) [1:15] <- c("22RV1_PROSTATE", "2313287_STOMACH", "253JBV_URINARY_TRACT", "253J_URINARY_TRACT", 
                                            "42MGBA_CENTRAL_NERVOUS_SYSTEM", "5637_URINARY_TRACT", "59M_OVARY", "639V_URINARY_TRACT",
                                            "647V_URINARY_TRACT", "697_HAEMATOPOIETIC_AND_LYMPHOID_TISSUE", "769P_KIDNEY", "786O_KIDNEY",
                                            "8305C_THYROID", "8505C_THYROID", "8MGBA_CENTRAL_NERVOUS_SYSTEM")
# check cell lines specific to data frames (transcript counts and metabolomics)
colnames(MRN_transcript_counts) [!(colnames(MRN_transcript_counts) %in% CCLE2019_metabolomics$CCLE_ID)] #106 cell lines specific to transcript counts
CCLE2019_metabolomics$CCLE_ID [!(CCLE2019_metabolomics$CCLE_ID %in% colnames(MRN_transcript_counts))] #15 cell lines specific to metabolomics
# remove cell lines specific to data frames, 913 cell lines overlapping between transcript counts and metabolomics
MRN_transcript_counts <- MRN_transcript_counts[,!colnames(MRN_transcript_counts) %in%
                                                colnames(MRN_transcript_counts) [!(colnames(MRN_transcript_counts) %in% CCLE2019_metabolomics$CCLE_ID)]]
CCLE2019_metabolomics <- CCLE2019_metabolomics[!CCLE2019_metabolomics$CCLE_ID %in% 
                                                CCLE2019_metabolomics$CCLE_ID [!(CCLE2019_metabolomics$CCLE_ID %in% colnames(MRN_transcript_counts))],]
# match the order of cell lines in transcript counts and metabolomics
CCLE2019_metabolomics <- t(CCLE2019_metabolomics)
colnames(CCLE2019_metabolomics) <- CCLE2019_metabolomics[1,] # cell line name into column name
CCLE2019_metabolomics = CCLE2019_metabolomics[-c(1,2),]
reorder_idx <- match(colnames(MRN_transcript_counts), colnames(CCLE2019_metabolomics))
CCLE2019_metabolomics <- CCLE2019_metabolomics[,reorder_idx]
all(colnames(MRN_transcript_counts) == colnames(CCLE2019_metabolomics)) # the cell line order is aligned between transcript counts and metabolomics
CCLE2019_metabolomics <- data.table(Metabolite = rownames(CCLE2019_metabolomics), CCLE2019_metabolomics)
fwrite(CCLE2019_metabolomics, "03 Results/01 Correlation Analysis/CCLE2019/CCLE2019 metabolomics.csv")

# cleaning after step 3
rm(reorder_idx)

##### 4 translate Ensembl name to gene symbol #####
# translation using biomaRt
transcript_name <- sub('\\.[0-9]*$', '', rownames(MRN_transcript_counts)) # remove the version info from Ensembl name
rownames(MRN_transcript_counts) = transcript_name
mart <- useMart(biomart = "ensembl", dataset = "hsapiens_gene_ensembl")
gene_name <- getBM(attributes = c("ensembl_gene_id","hgnc_symbol"),
                   filters = "ensembl_gene_id", values = transcript_name,
                   mart = mart)
# remove transcripts failed to translate
MRN_transcript_counts <- MRN_transcript_counts[!rownames(MRN_transcript_counts) %in% transcript_name [!(transcript_name %in% gene_name$ensembl_gene_id)],]

# returned 48431 rows in transcript counts, but gene name has 48432 rows
# the naming of one Ensembl name has changed across version and not updated: ENSG00000230417 can be translated to "LINC00595"(.12) and "LINC00856"(.6).
# Here "LINC00595" will be preserved.
gene_name <- gene_name[-which(gene_name$hgnc_symbol == "LINC00856"),]

# attach translated gene symbols to the transcript counts
SYMBOL <- c()
for (i in 1:nrow(MRN_transcript_counts)){
  ensembl_name <- rownames(MRN_transcript_counts)[i]
  symbol_name <- gene_name$hgnc_symbol[which(gene_name$ensembl_gene_id == ensembl_name)]
  SYMBOL <- c(SYMBOL, symbol_name)
  print(i)
}
MRN_transcript_counts$SYMBOL <- SYMBOL
MRN_transcript_counts <- MRN_transcript_counts %>% dplyr::select(SYMBOL, everything()) # move symbol column to the first column
MRN_transcript_counts <- MRN_transcript_counts[-which(MRN_transcript_counts$SYMBOL==""),] # remove blank symbol
fwrite(MRN_transcript_counts, "03 Results/01 Correlation Analysis/CCLE2019/CCLE2019 MRN normalised transcript counts.csv")

# cleaning after step 4
rm(mart)
rm(gene_name)
rm(ensembl_name)
rm(SYMBOL)
rm(i)
rm(symbol_name)
rm(transcript_name)

##### 5 separate SLC genes from translated gene symbol #####
MRN_transcript_counts <- fread("03 Results/01 Correlation Analysis/CCLE2019/CCLE2019 MRN normalised transcript counts.csv")
SLC_transcript_counts <- MRN_transcript_counts[SYMBOL %in% SLC_annotation_2024$HGNC_symbol] # 516 SLC recovered automatically
SLC_annotation_2024[!HGNC_symbol %in% SLC_transcript_counts$SYMBOL] # 3 SLC is left behind: SLC25A53, SLC5A3, SLC6A14
fwrite(SLC_transcript_counts, "03 Results/01 Correlation Analysis/CCLE2019/CCLE2019 SLC normalised transcript counts.csv")

##### 6 CCLE2019 SLC TvsM #####
# create empty matrix to store statistical output
SLC_transcript_counts <- fread("03 Results/01 Correlation Analysis/CCLE2019/CCLE2019 SLC normalised transcript counts.csv")
CCLE2019_metabolomics <- fread("03 Results/01 Correlation Analysis/CCLE2019/CCLE2019 metabolomics.csv")

SLC_OutputCor <- matrix(0, nrow=nrow(SLC_transcript_counts),ncol=nrow(CCLE2019_metabolomics))
SLC_OutputP <- matrix(1, nrow=nrow(SLC_transcript_counts),ncol=nrow(CCLE2019_metabolomics))

for(i in 1:nrow(SLC_transcript_counts)){
  for(j in 1:nrow(CCLE2019_metabolomics)){
    CorIn <- cor.test(as.numeric(SLC_transcript_counts[i,2:ncol(SLC_transcript_counts)]),
                      as.numeric(CCLE2019_metabolomics[j,2:ncol(CCLE2019_metabolomics)]), method="spearman", exact = FALSE)
    SLC_OutputCor[i,j] <- CorIn$estimate
    SLC_OutputP[i,j] <- CorIn$p.value
  }
  print(i)
}

# FDR for multiple correlation analysis
SLC_OutputPadj <- matrix(1, nrow=nrow(SLC_transcript_counts),ncol=nrow(CCLE2019_metabolomics))
for (k in 1:nrow(SLC_OutputP)) {
  SLC_OutputPadj[k,] <- p.adjust(SLC_OutputP[k,], method = "BH", n = length(SLC_OutputP[k,]))
}

SLC_OutputPadj <- as.data.frame(SLC_OutputPadj)
SLC_OutputCor <- as.data.frame(SLC_OutputCor)
colnames(SLC_OutputPadj) <- CCLE2019_metabolomics$Metabolite #set up rowname and colname of adjusted p value
colnames(SLC_OutputCor) <- CCLE2019_metabolomics$Metabolite #set up rowname and colname of correlation value
SLC_OutputPadj$SYMBOL <- SLC_transcript_counts$SYMBOL
SLC_OutputCor$SYMBOL <- SLC_transcript_counts$SYMBOL
SLC_OutputPadj <- SLC_OutputPadj %>% dplyr::select(SYMBOL, everything())
SLC_OutputCor <- SLC_OutputCor %>% dplyr::select(SYMBOL, everything())

# melt correlation matrices
mSLC_OutputPadj <- melt(SLC_OutputPadj, id.vars = "SYMBOL")
mSLC_OutputCor <- melt(SLC_OutputCor, id.vars = "SYMBOL")

# check if melted data frame has the same arrangement
all(mSLC_OutputPadj$SYMBOL == mSLC_OutputCor$SYMBOL) # TRUE
all(mSLC_OutputPadj$variable == mSLC_OutputCor$variable) # TRUE

SLC_interaction <- mSLC_OutputPadj
colnames(SLC_interaction) <- c("SYMBOL", "METABOLITE", "padj")
SLC_interaction$rho <- mSLC_OutputCor$value

fwrite(SLC_interaction, "03 Results/01 Correlation Analysis/CCLE2019/CCLE2019 TvsM.csv")
```
