## Supplementary material for "Predicting substrates for orphan Solute Carrier Proteins using multi-omics datasets": Code for generating results: NCI60a TvsM.R

```
##### 0 load libraries #####
library(ggplot2)
library(CePa)
library(data.table)
library(dplyr)
library(reshape2)
library(org.Hs.eg.db)

##### 1 read files #####
MRN_transcript_counts <- readRDS("01 Raw Data/01 Correlation Analysis/02 NCI60a/NCI60_RNAseq_normalised_counts.rds")
NCI60a_metabolomics <- fread("01 Raw Data/01 Correlation Analysis/02 NCI60a/WEB_DATA_METABOLON.csv") # 352 metabolite over 58 cell lines (https://wiki.nci.nih.gov/display/ncidtpdata/molecular+target+data)
SLC_annotation_2024 <- fread("01 Raw Data/00 SLC annotation 2024/SLC_transcript_20240321.csv", select = c(1:6))

##### 2 format transcriptomics and metabolomics #####
# dcast to form metabolite x cell line matrix
NCI60a_metabolomics <- NCI60a_metabolomics[,c(2,7,8)]
NCI60a_metabolomics <- dcast(data = NCI60a_metabolomics,formula = TITLE~cellname,fun.aggregate = sum,value.var = "VALUE")
# correct name format in metabolomics according to transcript data
overlap <- intersect(colnames(MRN_transcript_counts), colnames(NCI60a_metabolomics)[2:ncol(NCI60a_metabolomics)]) # 50 overlapping cellname
colnames(MRN_transcript_counts) [!colnames(MRN_transcript_counts) %in% overlap] # 10 transcript cell name not in overlap
colnames(NCI60a_metabolomics)[2:ncol(NCI60a_metabolomics)] [!colnames(NCI60a_metabolomics)[2:ncol(NCI60a_metabolomics)] %in% overlap] # 8 metabolomics cell name not in overlap
colnames(NCI60a_metabolomics) <- c( "Metabolite",      "786-0",           "A498",            "A549",            "ACHN",            "BT-549",          "CAKI-1",          "CCRF-CEM",       
                                    "COLO_205",        "DU-145",          "EKVX",            "HCC-2998",        "HCT-116",         "HCT-15",          "HL-60",           "HOP-62",         
                                    "HOP-92",          "HS_578T",         "HT29",            "IGROV1",          "K-562",           "KM12",            "LOX_IMVI",        "M14",            
                                    "MALME-3M",        "MCF7",            "MDA-MB-231",      "MDA-MB-435",      "MOLT-4",          "NCI-H226",        "NCI-H23",         "NCI-H322M",      
                                    "NCI-H460",        "NCI-H522",        "NCI_ADR-RES",     "OVCAR-3",         "OVCAR-4",         "OVCAR-5",         "OVCAR-8",         "PC-3",           
                                    "RPMI-8226",       "RXF_393",         "SF-268",          "SF-295",          "SF-539",          "SK-MEL-28",       "SK-MEL-5",        "SK-OV-3",        
                                    "SN12C",           "SNB-19",          "SNB-75",          "SR",              "SW-620",          "T-47D",           "TK-10",           "U251",           
                                    "UACC-257",        "UACC-62",         "UO-31"   )

# record of change: A549/ATCC --> A549; COLO 205 --> COLO_205; HL-60(TB) --> HL-60; HS 578T --> HS_578T; LOX IMVI --> LOX_IMVI;
#                   MDA-MB-231/ATCC --> MDA-MB-231; NCI/ADR-RES --> NCI_ADR-RES; RXF-393 --> RXF_393
overlap <- intersect(colnames(MRN_transcript_counts), colnames(NCI60a_metabolomics)[2:ncol(NCI60a_metabolomics)]) # second verify, 58 cell name matched
# remove unfound cell name in MRN transcript counts (SK-MEL-2, MDA-MB-468)
MRN_transcript_counts <- as.data.frame(MRN_transcript_counts[, -c(which(colnames(MRN_transcript_counts) == "SK-MEL-2" |
                                                                        colnames(MRN_transcript_counts) == "MDA-MB-468"))])
# translate entrez id to gene symbol
MRN_transcript_counts$SYMBOL <- mapIds(org.Hs.eg.db, keys = rownames(MRN_transcript_counts), keytype = "ENTREZID", column = "SYMBOL")
MRN_transcript_counts <- MRN_transcript_counts %>% dplyr::select(SYMBOL, everything())
# reorder cell name to match transcript data
reorder_idx <- match(colnames(MRN_transcript_counts)[2:ncol(MRN_transcript_counts)], colnames(NCI60a_metabolomics)[2:ncol(NCI60a_metabolomics)])
reorder_idx <- c(1, reorder_idx+1)
NCI60a_metabolomics <- NCI60a_metabolomics[, reorder_idx]
all(colnames(MRN_transcript_counts)[2:ncol(MRN_transcript_counts)] == colnames(NCI60a_metabolomics)[2:ncol(NCI60a_metabolomics)]) # the cell line order is aligned between transcript counts and metabolomics
fwrite(MRN_transcript_counts, "03 Results/01 Correlation Analysis/NCI60a/NCI60 MRN normalised transcript counts.csv")
fwrite(NCI60a_metabolomics, "03 Results/01 Correlation Analysis/NCI60a/NCI60a metabolomics.csv")

# cleaning after step 2
rm(overlap)
rm(reorder_idx)

##### 3 separate SLC genes from translated gene symbol #####
MRN_transcript_counts <- fread("03 Results/01 Correlation Analysis/NCI60a/NCI60 MRN normalised transcript counts.csv")
SLC_transcript_counts <- MRN_transcript_counts[which(MRN_transcript_counts$SYMBOL %in% SLC_annotation_2024$HGNC_symbol),] # 518 SLC transcripts recovered
SLC_annotation_2024[!HGNC_symbol %in% SLC_transcript_counts$SYMBOL] # 1 SLC is left behind: SLCO1B7
fwrite(SLC_transcript_counts, "03 Results/01 Correlation Analysis/NCI60a/NCI60 SLC normalised transcript counts.csv")

##### 4 NCI60a SLC TvsM #####
SLC_transcript_counts <- fread("03 Results/01 Correlation Analysis/NCI60a/NCI60 SLC normalised transcript counts.csv")
NCI60a_metabolomics <- fread("03 Results/01 Correlation Analysis/NCI60a/NCI60a metabolomics.csv")

# create empty matrix to store statistical output
SLC_OutputCor <- matrix(0, nrow=nrow(SLC_transcript_counts),ncol=nrow(NCI60a_metabolomics))
SLC_OutputP <- matrix(1, nrow=nrow(SLC_transcript_counts),ncol=nrow(NCI60a_metabolomics))

for(i in 1:nrow(SLC_transcript_counts)){
  for(j in 1:nrow(NCI60a_metabolomics)){
    CorIn <- cor.test(as.numeric(SLC_transcript_counts[i,2:ncol(SLC_transcript_counts)]),
                      as.numeric(NCI60a_metabolomics[j,2:ncol(NCI60a_metabolomics)]), method="spearman", exact = FALSE)
    SLC_OutputCor[i,j] <- CorIn$estimate
    SLC_OutputP[i,j] <- CorIn$p.value
  }
  print(i)
}

# FDR for multiple correlation analysis
SLC_OutputPadj <- matrix(1, nrow=nrow(SLC_transcript_counts),ncol=nrow(NCI60a_metabolomics))
for (k in 1:nrow(SLC_OutputP)) {
  SLC_OutputPadj[k,] <- p.adjust(SLC_OutputP[k,], method = "BH", n = length(SLC_OutputP[k,]))
}

SLC_OutputPadj <- as.data.frame(SLC_OutputPadj)
SLC_OutputCor <- as.data.frame(SLC_OutputCor)
colnames(SLC_OutputPadj) <- NCI60a_metabolomics$Metabolite #set up rowname and colname of adjusted p value
colnames(SLC_OutputCor) <- NCI60a_metabolomics$Metabolite #set up rowname and colname of correlation value
SLC_OutputPadj$SYMBOL <- SLC_transcript_counts$SYMBOL
SLC_OutputCor$SYMBOL <- SLC_transcript_counts$SYMBOL
SLC_OutputPadj <- SLC_OutputPadj %>% dplyr::select(SYMBOL, everything())
SLC_OutputCor <- SLC_OutputCor %>% dplyr::select(SYMBOL, everything())

# melt correlation matrices
mSLC_OutputPadj <- melt(SLC_OutputPadj, id.vars = "SYMBOL")
mSLC_OutputCor <- melt(SLC_OutputCor, id.vars = "SYMBOL")

# check if melted data frame has the same arrangement
all(mSLC_OutputPadj$SYMBOL == mSLC_OutputCor$SYMBOL) # TRUE
all(mSLC_OutputPadj$variable == mSLC_OutputCor$variable) # TRUE

SLC_interaction <- mSLC_OutputPadj
colnames(SLC_interaction) <- c("SYMBOL", "METABOLITE", "padj")
SLC_interaction$rho <- mSLC_OutputCor$value

fwrite(SLC_interaction, "03 Results/01 Correlation Analysis/NCI60a/NCI60a TvsM.csv")
```
