## Supplementary material for "Predicting substrates for orphan Solute Carrier Proteins using multi-omics datasets": Code for generating results: NCI60b TvsM (abort).R

```
##### 0 load libraries #####
library(data.table)
library(dplyr)
library(org.Hs.eg.db)
##### 1 read files #####
MRN_transcript_counts = readRDS("1 Raw Data/02 NCI60a TvsM/NCI60_RNAseq_normalised_counts.rds") 
NCI60b_metabolomics = fread("1 Raw Data/03 NCI60b TvsM/MOESM2_ESM.csv") # 2181 metabolite over 54 cell lines (https://doi.org/10.1038/s41467-019-09695-9)
SLC_annotation_2023 = fread("1 Raw Data/00 SLC annotation 2023/SLC annotation 2023.csv")
MRN_transcript_counts = as.data.frame(MRN_transcript_counts)
NCI60b_metabolomics = as.data.frame(NCI60b_metabolomics)
SLC_annotation_2023 = as.data.frame(SLC_annotation_2023)

##### 2 format transcriptomics and metabolomics #####
NCI60b_metabolomics = NCI60b_metabolomics[,-c(1,2,4,5)]
colnames(NCI60b_metabolomics) = gsub(".*_", "", colnames(NCI60b_metabolomics))
overlap = intersect(colnames(MRN_transcript_counts), colnames(NCI60b_metabolomics)[2:ncol(NCI60b_metabolomics)]) # 11 overlapping cellname
colnames(MRN_transcript_counts) [!colnames(MRN_transcript_counts) %in% overlap] # 49 transcript cell name not in overlap
colnames(NCI60b_metabolomics)[2:ncol(NCI60b_metabolomics)] [!colnames(NCI60b_metabolomics)[2:ncol(NCI60b_metabolomics)] %in% overlap] # 44 metabolomics cell name not in overlap
colnames(NCI60b_metabolomics) = c( "metabolites","BT-549",     "HS_578T",    "MCF7",       "MDA-MB-231", "T-47D",      "SF-268",     "SF-295",
                                   "SF-539",     "SNB-19",     "SNB-75",     "U251",       "COLO_205",   "HCC-2998",   "HCT-116",    "HCT-15",
                                   "HT29",       "KM12",       "SW-620",     "A549",       "EKVX",       "HOP-62",     "HOP-92",     "NCI-H226",   
                                   "NCI-H23",    "NCI-H322M",  "NCI-H460",   "NCI-H522",   "MDA-MB-468", "LOX_IMVI",   "M14",        "MALME-3M",
                                   "MDA-MB-435", "SK-MEL-2",   "SK-MEL-28",  "SK-MEL-5",   "UACC-257",   "UACC-62",    "IGROV1",     "NCI_ADR-RES",  
                                   "OVCAR-3",    "OVCAR-4",    "OVCAR-5",    "OVCAR-8",    "SK-OV-3",    "DU-145",     "PC-3",       "786-0",      
                                   "A498",       "ACHN",       "CAKI-1",      "RXF_393",     "SN12C",      "TK-10",       "UO-31"   )
# record of change: BT549 --> BT-549; HS578T --> HS_578T; MDAMB231 --> MDA-MB-231; T47D --> T-47D; SF268 --> SF-268; SF295 --> SF-295; SF539 --> SF-539;
#                   SNB19 --> SNB-19; SNB75 --> SNB-75; COLO205 --> COLO_205; HCC2998 --> HCC-2998; HCT116 --> HCT-116; HCT15 --> HCT-15; SW620 --> SW-620;
#                   HOP62 --> HOP-62; HOP92 --> HOP-92; NCIH226 --> NCI-H226; NCIH23 --> NCI-H23; NCIH322M --> NCI-H322M; NCIH460 --> NCI-H460; NCIH522 --> NCI-H522;
#                   MDAMB468 --> MDA-MB-468; LOXIMVI --> LOX_IMVI; MALME3M --> MALME-3M; MDAMB435 --> MDA-MB-435; SKMEL2 --> SK-MEL-2; SKMEL28 --> SK-MEL-28;
#                   SKMEL5 --> SK-MEL-5; UACC257 --> UACC-257; UACC62 --> UACC-62; NCIADRRES --> NCI_ADR-RES; OVCAR3 --> OVCAR-3; OVCAR4 --> OVCAR-4; OVCAR5 --> OVCAR-5;
#                   OVCAR8 --> OVCAR-8; SKOV3 --> SK-OV-3; DU145 --> DU-145; PC3 --> PC-3; 786O --> 786-0; CAKI1 --> CAKI-1; RXF393 --> RXF_393; TK10 --> TK-10;
#                   UO31 --> UO-31
overlap = intersect(colnames(MRN_transcript_counts), colnames(NCI60b_metabolomics)[2:ncol(NCI60b_metabolomics)]) # second verify, 54 cell name matched
# remove unfound cell name in MRN transcript counts (RPMI-8226, SR, K-562, MOLT-4, HL-60, CCRF-CEM)
MRN_transcript_counts = as.data.frame(MRN_transcript_counts[, -c(which(colnames(MRN_transcript_counts) == "RPMI-8226" |
                                                                       colnames(MRN_transcript_counts) == "SR" |
                                                                       colnames(MRN_transcript_counts) == "K-562" |
                                                                       colnames(MRN_transcript_counts) == "MOLT-4" |
                                                                       colnames(MRN_transcript_counts) == "HL-60" |
                                                                       colnames(MRN_transcript_counts) == "CCRF-CEM"))])
# translate entrez id to gene symbol
MRN_transcript_counts$SYMBOL = mapIds(org.Hs.eg.db, keys = rownames(MRN_transcript_counts), keytype = "ENTREZID", column = "SYMBOL")
MRN_transcript_counts = MRN_transcript_counts %>% dplyr::select(SYMBOL, everything())
# reorder cell name to match transcript data
reorder_idx = match(colnames(MRN_transcript_counts)[2:ncol(MRN_transcript_counts)], colnames(NCI60b_metabolomics)[2:ncol(NCI60b_metabolomics)])
reorder_idx = c(1, reorder_idx+1)
NCI60b_metabolomics = NCI60b_metabolomics[, reorder_idx]
all(colnames(MRN_transcript_counts)[2:ncol(MRN_transcript_counts)] == colnames(NCI60b_metabolomics)[2:ncol(NCI60b_metabolomics)]) # the cell line order is aligned between transcript counts and metabolomics
write.csv(MRN_transcript_counts, "3 Results/03 NCI60b TvsM/transcript counts/NCI60b MRN normalised transcript counts.csv")

##### 3 separate SLC genes from translated gene symbol #####
SLC_transcript_counts = MRN_transcript_counts[which(MRN_transcript_counts$SYMBOL %in% SLC_annotation_2023$HGNC_symbol),] # 519 SLC transcripts recovered
SLC_index = which(rownames(MRN_transcript_counts) %in% rownames(SLC_transcript_counts))
nonSLC_transcript_counts = MRN_transcript_counts[-SLC_index,]
write.csv(SLC_transcript_counts, "3 Results/03 NCI60b TvsM/transcript counts/NCI60b SLC transcript counts.csv")
write.csv(nonSLC_transcript_counts, "3 Results/03 NCI60b TvsM/transcript counts/NCI60b non-SLC transcript counts.csv")

##### 4 NCI60b SLC TvsM #####
# create empty matrix to store statistical output
SLC_OutputCor = matrix(0, nrow=nrow(SLC_transcript_counts),ncol=nrow(NCI60b_metabolomics))
SLC_OutputP = matrix(1, nrow=nrow(SLC_transcript_counts),ncol=nrow(NCI60b_metabolomics))

for(i in 1:nrow(SLC_transcript_counts)){
  for(j in 1:nrow(NCI60b_metabolomics)){
    CorIn = cor.test(as.numeric(SLC_transcript_counts[i,2:ncol(SLC_transcript_counts)]),
                     as.numeric(NCI60b_metabolomics[j,2:ncol(NCI60b_metabolomics)]), method="spearman", exact = FALSE)
    SLC_OutputCor[i,j] = CorIn$estimate
    SLC_OutputP[i,j] = CorIn$p.value
  }
  print(i)
}

# standard deviation will be zero for more than 50 interactions, this is because some SLC are not well expressed across cell lines (i.e. most are zeros)
# and the metabolomic profile for these cell lines happen to be NaN, leading to the omission of these cell line, and rest cell lines indeed have sd = 0
# NA will be introduced to such interaction so it will not affect the predictions
# only SLC transcript row 319 (SLC35G4) show ~80 warnings starting from metabolomics row 82, this SLC only has detected transcripts in 2 cell lines


# FDR for multiple correlation analysis
SLC_OutputPadj = matrix(1, nrow=nrow(SLC_transcript_counts),ncol=nrow(NCI60b_metabolomics))
for (k in 1:nrow(SLC_OutputP)) {
  SLC_OutputPadj[k,] = p.adjust(SLC_OutputP[k,], method = "BH", n = length(SLC_OutputP[k,]))
}

SLC_OutputPadj = as.data.frame(SLC_OutputPadj)
SLC_OutputCor = as.data.frame(SLC_OutputCor)
colnames(SLC_OutputPadj) = NCI60b_metabolomics$metabolites #set up rowname and colname of adjusted p value
colnames(SLC_OutputCor) = NCI60b_metabolomics$metabolites #set up rowname and colname of correlation value
SLC_OutputPadj$SYMBOL = SLC_transcript_counts$SYMBOL
SLC_OutputCor$SYMBOL = SLC_transcript_counts$SYMBOL
SLC_OutputPadj = SLC_OutputPadj %>% dplyr::select(SYMBOL, everything())
SLC_OutputCor = SLC_OutputCor %>% dplyr::select(SYMBOL, everything())

write.csv(SLC_OutputCor, "3 Results/03 NCI60b TvsM/correlation matrix/NCI60b SLC TvsM correlation matrix rho.csv")
write.csv(SLC_OutputPadj, "3 Results/03 NCI60b TvsM/correlation matrix/NCI60b SLC TvsM correlation matrix padj.csv")

# melt correlation matrices
mSLC_OutputPadj = melt(SLC_OutputPadj, id.vars = "SYMBOL")
mSLC_OutputCor = melt(SLC_OutputCor, id.vars = "SYMBOL")

# check if melted data frame has the same arrangement
all(mSLC_OutputPadj$SYMBOL == mSLC_OutputCor$SYMBOL) # TRUE
all(mSLC_OutputPadj$variable == mSLC_OutputCor$variable) # TRUE

SLC_interaction = mSLC_OutputPadj
colnames(SLC_interaction) = c("SYMBOL", "METABOLITE", "padj")
SLC_interaction$rho = mSLC_OutputCor$value

write.csv(SLC_interaction, "3 Results/03 NCI60b TvsM/interactions/NCI60b SLC TvsM interaction.csv")

##### 4 NCI60b SLC TvsM #####
# create empty matrix to store statistical output
nonSLC_OutputCor = matrix(0, nrow=nrow(nonSLC_transcript_counts),ncol=nrow(NCI60b_metabolomics))
nonSLC_OutputP = matrix(1, nrow=nrow(nonSLC_transcript_counts),ncol=nrow(NCI60b_metabolomics))

for(i in 10001:nrow(nonSLC_transcript_counts)){
  for(j in 1:nrow(NCI60b_metabolomics)){
    CorIn = cor.test(as.numeric(nonSLC_transcript_counts[i,2:ncol(nonSLC_transcript_counts)]),
                     as.numeric(NCI60b_metabolomics[j,2:ncol(NCI60b_metabolomics)]), method="spearman", exact = FALSE)
    nonSLC_OutputCor[i,j] = CorIn$estimate
    nonSLC_OutputP[i,j] = CorIn$p.value
  }
  print(i)
}

# FDR for multiple correlation analysis
nonSLC_OutputPadj = matrix(1, nrow=nrow(nonSLC_transcript_counts),ncol=nrow(NCI60b_metabolomics))
for (k in 1:nrow(nonSLC_OutputP)) {
  nonSLC_OutputPadj[k,] = p.adjust(nonSLC_OutputP[k,], method = "BH", n = length(nonSLC_OutputP[k,]))
}

nonSLC_OutputPadj = as.data.frame(nonSLC_OutputPadj)
nonSLC_OutputCor = as.data.frame(nonSLC_OutputCor)
colnames(nonSLC_OutputPadj) = NCI60b_metabolomics$metabolites #set up rowname and colname of adjusted p value
colnames(nonSLC_OutputCor) = NCI60b_metabolomics$metabolites #set up rowname and colname of correlation value
nonSLC_OutputPadj$SYMBOL = nonSLC_transcript_counts$SYMBOL
nonSLC_OutputCor$SYMBOL = nonSLC_transcript_counts$SYMBOL
nonSLC_OutputPadj = nonSLC_OutputPadj %>% dplyr::select(SYMBOL, everything())
nonSLC_OutputCor = nonSLC_OutputCor %>% dplyr::select(SYMBOL, everything())

fwrite(nonSLC_OutputCor, "3 Results/03 NCI60b TvsM/correlation matrix/NCI60b non-SLC TvsM correlation matrix rho.csv")
fwrite(nonSLC_OutputPadj, "3 Results/03 NCI60b TvsM/correlation matrix/NCI60b non-SLC TvsM correlation matrix padj.csv")

# melt correlation matrices
mnonSLC_OutputPadj = melt(nonSLC_OutputPadj, id.vars = "SYMBOL")
mnonSLC_OutputCor = melt(nonSLC_OutputCor, id.vars = "SYMBOL")

# check if melted data frame has the same arrangement
all(mnonSLC_OutputPadj$SYMBOL == mnonSLC_OutputCor$SYMBOL) # TRUE
all(mnonSLC_OutputPadj$variable == mnonSLC_OutputCor$variable) # TRUE

nonSLC_interaction = mnonSLC_OutputPadj
colnames(nonSLC_interaction) = c("SYMBOL", "METABOLITE", "padj")
nonSLC_interaction$rho = mnonSLC_OutputCor$value

fwrite(nonSLC_interaction, "3 Results/03 NCI60b TvsM/interactions/NCI60b non-SLC TvsM interaction.csv")
```
