## Supplementary material for "Predicting substrates for orphan Solute Carrier Proteins using multi-omics datasets": Code for generating results: NCI60extra.R

```
library(data.table)
library(org.Hs.eg.db)
library(dplyr)
library(reshape2)
library(ggplot2)

##### 1 read files #####
MRN_transcript_counts = readRDS("01 Raw Data/01 Predictive Scores/02 NCI60a/NCI60_RNAseq_normalised_counts.rds")
NCI60extra_metabolomics = fread("01 Raw Data/01 Predictive Scores/05 NCI60extra/NCI60CORE.csv")
NCI60extra_sample1 = NCI60extra_metabolomics[,c(1,2,3, seq(4, ncol(NCI60extra_metabolomics), by = 2)), with = FALSE]
NCI60extra_sample2 = NCI60extra_metabolomics[,c(1,2,3, seq(5, ncol(NCI60extra_metabolomics), by = 2)), with = FALSE]
SLC_annotation_2023 = fread("01 Raw Data/00 SLC annotation 2023/SLC ontology annotation 2023.csv")

##### 2 prepare extracellular metabolomics for correlation #####
# adjust format #
overlap = intersect(colnames(MRN_transcript_counts), colnames(NCI60extra_sample1)[4:ncol(NCI60extra_sample1)])
colnames(MRN_transcript_counts) [!colnames(MRN_transcript_counts) %in% overlap]
colnames(NCI60extra_sample1)[4:ncol(NCI60extra_sample1)] [!colnames(NCI60extra_sample1)[4:ncol(NCI60extra_sample1)] %in% overlap]
metaolomics_cell_name_corrected = c("Method (a)",      "Metabolite (b)",  "Calibrated (e)",  "UACC-257",        "OVCAR-8",         "OVCAR-5",         "SF-295",         
                                    "A549",            "RXF_393",         "CAKI-1",          "MDA-MB-435",      "EKVX",            "NCI-H23",         "HCC-2998",       
                                    "NCI_ADR-RES",     "HT29",            "M14",             "HCT-116",         "HOP-62",          "786-0",           "NCI-H322M",      
                                    "BT-549",          "MALME-3M",        "TK-10",           "UO-31",           "T-47D",           "MCF7",            "NCI-H460",       
                                    "SK-OV-3",         "SK-MEL-5",        "SNB-19",          "SF-539",          "COLO_205",        "KM12",            "OVCAR-3",        
                                    "SF-268",          "DU-145",          "HS_578T",         "SW-620",          "SK-MEL-28",       "A498",            "IGROV1",         
                                    "MOLT-4",          "K-562",           "HOP-92",          "PC-3",            "HL-60",           "MDA-MB-468",      "SK-MEL-2",       
                                    "NCI-H522",        "HCT-15",          "ACHN",            "U251",            "LOX_IMVI",        "SNB-75",          "OVCAR-4",        
                                    "NCI-H226",        "UACC-62",         "SN12C",           "MDA-MB-231",      "CCRF-CEM",        "RPMI-8226",       "SR"  )
# record of change: "A549/ATCC" --> "A549"; "RXF 393" --> "RXF_393"; "NCI/ADR-RES" --> "NCI_ADR-RES"; "786-O" --> "786-0"; "COLO 205" --> "COLO_205";
#                   "HS 578T" --> "HS_578T"; "SW620" --> "SW-620"; "K562" --> "K-562"; "HL-60(TB)" --> "HL-60"; "LOX IMVI" --> "LOX_IMVI"; 
#                   "MDA-MB-231/ATCC" --> "MDA-MB-231"; "RPMI 8226" --> "RPMI-8226"
colnames(NCI60extra_sample1) = metaolomics_cell_name_corrected
colnames(NCI60extra_sample2) = metaolomics_cell_name_corrected

##### 3 prepare MRN_transcript counts for correlation #####
# translate entrez id to gene symbol
MRN_transcript_counts = as.data.frame(MRN_transcript_counts)
MRN_transcript_counts$SYMBOL = mapIds(org.Hs.eg.db, keys = rownames(MRN_transcript_counts), keytype = "ENTREZID", column = "SYMBOL")
MRN_transcript_counts = MRN_transcript_counts %>% dplyr::select(SYMBOL, everything())
# reorder cell name to match transcript data
reorder_idx = match(colnames(MRN_transcript_counts)[2:ncol(MRN_transcript_counts)], colnames(NCI60extra_sample1)[4:ncol(NCI60extra_sample1)])
reorder_idx = c(1,2,3, reorder_idx+3)
NCI60extra_sample1 = NCI60extra_sample1[, reorder_idx, with = FALSE]
NCI60extra_sample2 = NCI60extra_sample2[, reorder_idx, with = FALSE]
# separate SLC genes from translated gene symbol #
SLC_transcript_counts = MRN_transcript_counts[which(MRN_transcript_counts$SYMBOL %in% SLC_annotation_2023$HGNC_symbol),] # 518 SLC transcripts recovered

##### 4 NCI60extra SLC TvsM #####
# create empty matrix to store statistical output
sample1_Cor = matrix(0, nrow=nrow(SLC_transcript_counts),ncol=nrow(NCI60extra_sample1))
sample1_P = matrix(1, nrow=nrow(SLC_transcript_counts),ncol=nrow(NCI60extra_sample1))
sample2_Cor = matrix(0, nrow=nrow(SLC_transcript_counts),ncol=nrow(NCI60extra_sample2))
sample2_P = matrix(1, nrow=nrow(SLC_transcript_counts),ncol=nrow(NCI60extra_sample2))

for(i in 1:nrow(SLC_transcript_counts)){
  for(j in 1:nrow(NCI60extra_sample1)){
    # Uptake metabolomics #
    CorIn1 = cor.test(as.numeric(SLC_transcript_counts[i,2:ncol(SLC_transcript_counts)]),
                      as.numeric(NCI60extra_sample1[j,4:ncol(NCI60extra_sample1)]), method="spearman", exact = FALSE)
    sample1_Cor[i,j] = CorIn1$estimate
    sample1_P[i,j] = CorIn1$p.value
    # Secretion metabolomics #
    CorIn2 = cor.test(as.numeric(SLC_transcript_counts[i,2:ncol(SLC_transcript_counts)]),
                      as.numeric(NCI60extra_sample2[j,4:ncol(NCI60extra_sample2)]), method="spearman", exact = FALSE)
    sample2_Cor[i,j] = CorIn2$estimate
    sample2_P[i,j] = CorIn2$p.value
  }
  if (i %% 10 == 0){
    print(i)
  }
}

# FDR for multiple correlation analysis
sample1_Padj = matrix(1, nrow=nrow(SLC_transcript_counts),ncol=nrow(NCI60extra_sample1))
for (k in 1:nrow(sample1_P)) {
  sample1_Padj[k,] = p.adjust(sample1_P[k,], method = "BH", n = length(sample1_P[k,]))
}

sample2_Padj = matrix(1, nrow=nrow(SLC_transcript_counts),ncol=nrow(NCI60extra_sample2))
for (k in 1:nrow(sample2_P)) {
  sample2_Padj[k,] = p.adjust(sample2_P[k,], method = "BH", n = length(sample2_P[k,]))
}

sample1_Padj = as.data.table(sample1_Padj)
sample2_Padj = as.data.table(sample2_Padj)
sample1_Cor = as.data.table(sample1_Cor)
sample2_Cor = as.data.table(sample2_Cor)

colnames(sample1_Padj) = NCI60extra_sample1$`Metabolite (b)`
colnames(sample1_Cor) = NCI60extra_sample1$`Metabolite (b)` 
colnames(sample2_Padj) = NCI60extra_sample2$`Metabolite (b)` 
colnames(sample2_Cor) = NCI60extra_sample2$`Metabolite (b)` 

sample1_Padj$SYMBOL = SLC_transcript_counts$SYMBOL
sample1_Cor$SYMBOL = SLC_transcript_counts$SYMBOL
sample2_Padj$SYMBOL = SLC_transcript_counts$SYMBOL
sample2_Cor$SYMBOL = SLC_transcript_counts$SYMBOL

sample1_Padj = sample1_Padj %>% dplyr::select(SYMBOL, everything())
sample1_Cor = sample1_Cor %>% dplyr::select(SYMBOL, everything())
sample2_Padj = sample2_Padj %>% dplyr::select(SYMBOL, everything())
sample2_Cor = sample2_Cor %>% dplyr::select(SYMBOL, everything())

mSample1 = melt(sample1_Cor, id.vars = "SYMBOL")
mSample1$padj = melt(sample1_Padj, id.vars = "SYMBOL")$value
colnames(mSample1) = c("SYMBOL", "METABOLITE", "rho", "padj")
mSample1 = as.data.table(mSample1)

mSample2 = melt(sample2_Cor, id.vars = "SYMBOL")
mSample2$padj = melt(sample2_Padj, id.vars = "SYMBOL")$value
colnames(mSample2) = c("SYMBOL", "METABOLITE", "rho", "padj")
mSample2 = as.data.table(mSample2)

##### 5 Predictive Scores #####
# CalculatePredictive #
CalculatePredictive = function(interactions){
  new_interactions = data.table(SYMBOL = rep(NA, nrow(interactions)), 
                                METABOLITE = rep(NA, nrow(interactions)), 
                                Predictive_Scores = rep(NA, nrow(interactions))
  )
  compound = unique(interactions$METABOLITE)
  for (i in 1:length(compound)){
    matrix_atomic_compound = interactions[which(interactions$METABOLITE == compound[i]),]
    sd_atomic_compound = sd(abs(matrix_atomic_compound$rho))
    mean_atomic_compound = mean(abs(matrix_atomic_compound$rho))
    matrix_atomic_compound$ps = (abs(matrix_atomic_compound$rho) - mean_atomic_compound)/sd_atomic_compound
    start = 1 + nrow(matrix_atomic_compound) * (i - 1)
    end = nrow(matrix_atomic_compound) * i
    new_interactions$SYMBOL[start:end] = matrix_atomic_compound$SYMBOL
    new_interactions$METABOLITE[start:end] = as.character(matrix_atomic_compound$METABOLITE)
    new_interactions$Predictive_Scores[start:end] = matrix_atomic_compound$ps
  }
  return(new_interactions)
}

NCI60extra_sample1_SLC_Predictive = CalculatePredictive(mSample1)
NCI60extra_sample2_SLC_Predictive = CalculatePredictive(mSample2)

fwrite(NCI60extra_sample1_SLC_Predictive, "03 Results/05 NCI60extra/NCI60extra sample1 predictive.csv")
fwrite(NCI60extra_sample2_SLC_Predictive, "03 Results/05 NCI60extra/NCI60extra sample2 predictive.csv")

##### 6 Benchmark #####
tb = fread("01 Raw Data/00 SLC annotation 2023/book construction/step 4 4 matrices book full.csv")
ki = fread("01 Raw data/05 Known interaction/Known interaction formula update with extra.csv")

# ConvertNames #
ConvertNames = function(name, source){
  # Process source input #
  if (is.null(source)){
    stop("Source of metabolite name required!")
  }
  else if (source %in% colnames(tb)){
    # Process name input #
    if (is.null(name)){
      stop("Name of metabolite required!")
    }else if (name %in% tb[,get(source)]){
      ans = tb[get(source) == name]
    }else{
      stop("Name entered not recognised!")
    }
  }else{
    stop("Source entered not recognised!")
  }
  
  # return ans as result #
  return(ans)
}

# BuildEvidenceTable #
BuildEvidenceTable = function(gene, formula){
  # build table #
  ans = data.table(SYMBOL = gene, Formula = formula, NCI60extra_sample1 = NA, NCI60extra_sample2 = NA)
  # process formula #
  line = try(ConvertNames(formula, "Formula"), silent = TRUE)
  if (grepl("Error", line[[1]]) == TRUE){
    stop("Error in converting formula!")
  }else{
    compound_info = ConvertNames(formula, "Formula")
  }
  info_source = c("NCI60extra_sample1", "NCI60extra_sample2")
  for (i in 1:length(info_source)){
    if (is.na(compound_info[,NCI60extra]) == TRUE){
      ans[, info_source[i]:=NA]
    }else{
      ans[, info_source[i]:=get(paste(info_source[i], "_SLC_Predictive", sep = ""))
          [SYMBOL == gene & METABOLITE == compound_info[, NCI60extra]][,Predictive_Scores]]
    }
  }
  return(ans)
}

# constructing a NCI60-extra specific evidence table #
et = data.table(SYMBOL = character(), Formula = character(), NCI60extra_sample1 = numeric(), NCI60extra_sample2 = numeric())
for (i in 1:length(SLC_transcript_counts$SYMBOL)){
  for (j in 1:length(tb$Formula)){
    temp = BuildEvidenceTable(SLC_transcript_counts$SYMBOL[i], tb$Formula[j])
    et = rbind(et, temp)
    print(paste(i, ":", j/length(tb$Formula)))
  }
}
et = et[rowSums(is.na(et[,3:4]))<2] # remove rows with NA in both sample

# constructing a NCI60-extra specific etki #
ki = ki[rowSums(is.na(ki[,4:5]))<2]
etki = ki
etki$NCI60extra_sample1 = vector("numeric", nrow(etki))
etki$NCI60extra_sample2 = vector("numeric", nrow(etki))

for (i in 1:nrow(etki)){
  if (etki$HGNC_symbol[i] %in% unique(et$SYMBOL) & etki$Formula[i] %in% unique(et$Formula)){
    etki[i, c("NCI60extra_sample1", "NCI60extra_sample2") :=
           as.list(et[SYMBOL == etki$HGNC_symbol[i] & Formula == etki$Formula[i]][,NCI60extra_sample1:NCI60extra_sample2])]
  }else{
    etki[i, c("NCI60extra_sample1", "NCI60extra_sample2") := list(NA, NA)]
  }
}
etki = etki[rowSums(is.na(etki[,6:7]))<2] # remove rows with NA in both sample

# investigate concordance #
ggplot(etki, aes(x=NCI60extra_sample1, y=NCI60extra_sample2)) + 
  geom_point()
ggplot(etki, aes(x=NCI60extra_sample1, y=NCI60extra_sample2, color = Localisation)) + 
  geom_point()

# generating fake etki for NCI60-extra #

# initialise fake etki collection
fake_etki_collection = vector("list", 100)
etki_gene_pool = unique(etki$HGNC_symbol)
etki_metabolite_pool = unique(etki$Formula)

for (sample_time in 1:100){
  # initialise fake etki data table #
  fake_etki = setNames(data.table(matrix(ncol = ncol(etki), nrow = nrow(etki))), colnames(etki))
  fake_etki[, Substrate := NULL]
  char.col = colnames(fake_etki)[1:4]
  num.col = colnames(fake_etki)[5:6]
  fake_etki[, (char.col) := lapply(.SD, as.character), .SDcols = char.col]
  fake_etki[, (num.col) := lapply(.SD, as.character), .SDcols = num.col]
  fake_etki[, c("HGNC_symbol", "Localisation") := as.list(etki[, c("HGNC_symbol", "Localisation")])]
  fake_etki[, Status := rep("Fake", nrow(fake_etki))]
  # for each SLC, shuffle fake substrate formula #
  for (j in 1:nrow(fake_etki)){
    # produce fake substrate pools with considerations of adjacency #
    true_substrate = etki[HGNC_symbol == fake_etki$HGNC_symbol[j]][, Formula]
    fake_substrate_pool = etki_metabolite_pool[!etki_metabolite_pool %in% true_substrate]
    fake_substrate = sample(fake_substrate_pool, 1)
    while (nrow(et[SYMBOL == fake_etki$HGNC_symbol[j] & Formula == fake_substrate][,NCI60extra_sample1:NCI60extra_sample2]) == 0){
      fake_substrate = sample(fake_substrate_pool, 1)
    }
    fake_etki[j, Formula := fake_substrate]
    fake_etki[j, c("NCI60extra_sample1", "NCI60extra_sample2") := 
                as.list(et[SYMBOL == fake_etki$HGNC_symbol[j] & Formula == fake_etki$Formula[j]][,NCI60extra_sample1:NCI60extra_sample2])]
    print(paste(sample_time, ":", j/nrow(fake_etki)))
  }
  fake_etki_collection[[sample_time]] = fake_etki
  #fwrite(fake_etki, paste("03 Results/08 Matrix Assembly/fake etki/Fake No", sample_time, ".csv", sep = ""))
  print(paste(sample_time, "done"))
}

# NCI60-extra specific confidence score calculation (only based on predictive scores) #
ps_dist = na.omit(as.numeric(unlist(as.list(etki[,6:7]))))
ps_dist = ps_dist[ps_dist > 0]
q = quantile(ps_dist, seq(0, 1, by = 0.1))

ConfidenceScore = function(ki.data, threshold = NULL){
  ki_score = setNames(data.table(matrix(ncol = 6, nrow = nrow(ki.data))), c("HGNC_symbol","Localisation","Substrate","Formula","Status","Score"))
  char.col = colnames(ki_score)[1:5]
  num.col = colnames(ki_score)[6]
  ki_score[, (char.col) := lapply(.SD, as.character), .SDcols = char.col]
  ki_score[, (num.col) := lapply(.SD, as.character), .SDcols = num.col]
  ki_score[, 1:5 := as.list(ki.data[, 1:5])]
  
  if (is.null(threshold)){
    threshold = 0
  }
  Factor_A_name = c("NCI60extra_sample1", "NCI60extra_sample2")
  
  for (i in 1:nrow(ki.data)){
    # score prediction for 3 matrices #
    Factor_A = vector("integer", length(Factor_A_name))
    for (j in 1:length(Factor_A_name)){
      if (is.na(ki.data[i, get(Factor_A_name[j])]) | ki.data[i, get(Factor_A_name[j])] < threshold){
        Factor_A[j] = 0
      } else if (ki.data[i, get(Factor_A_name[j])] > threshold){
        Factor_A[j] = findInterval(ki.data[i, get(Factor_A_name[j])], q)
      }
    }
    Factor_A = sum(Factor_A)
    ki_score[i, Score := Factor_A]
  }
  
  return(ki_score)
}

# etki confidence score (ecs) across threshold #
ecs_across_threshold = data.table(Threshold = seq(0, 1, by = 0.1), 
                                  Status = rep("ECS", length(seq(0, 1, by = 0.1))),
                                  Mean = vector("numeric", length(seq(0, 1, by = 0.1))))
for (i in 1:nrow(ecs_across_threshold)){
  ecs = ConfidenceScore(etki, ecs_across_threshold[i, Threshold])
  ecs_across_threshold[i, Mean := mean(as.numeric(ecs[,Score]))]
  print(i)
}

# fake etki confidence score (fecs) across threshold #
fecs_across_threshold = data.table(Threshold = rep(seq(0, 1, by = 0.1), each = 100), 
                                   Status = rep("ECS", length(rep(seq(0, 1, by = 0.1), each = 100))),
                                   Mean = vector("numeric", length(rep(seq(0, 1, by = 0.1), each = 100))))
for (t in seq(0, 1, by = 0.1)){
  fecs_confidence_score = lapply(fake_etki_collection, function(x){ConfidenceScore(x, t)})
  fecs_confidence_score_mean = unlist(lapply(fecs_confidence_score, function(x){mean(as.numeric(x[,Score]))}))
  fecs_across_threshold[Threshold == t, Mean := fecs_confidence_score_mean]
  print(t)
}

# plot ecs with fecs across threshold #
ggplot() +
  # box plot of fecs
  geom_boxplot(data = fecs_across_threshold, 
               aes(x = Threshold, y= Mean, group = Threshold)) +
  # points of ecs
  geom_point(data = ecs_across_threshold,
             aes(x= Threshold, y= Mean),
             color = 'red') +
  ylim(1, 8)

# plot ecs with feces across threshold considering adjacency and localisation #
ecs_across_threshold = data.table(Threshold = rep(seq(0, 1, by = 0.1), each = 3), 
                                  Status = rep(c("PS", "PS-IC", "IC"), 11),
                                  Mean = vector("numeric", length(rep(seq(0, 1, by = 0.1), each = 3))))
for (i in unique(ecs_across_threshold$Threshold)){
  ecs = ConfidenceScore(etki, i)
  ecs_across_threshold[Threshold == i & Status == "PS", Mean := mean(as.numeric(ecs[Localisation == "PS", Score]))]
  ecs_across_threshold[Threshold == i & Status == "PS-IC", Mean := mean(as.numeric(ecs[Localisation == "PS-IC", Score]))]
  ecs_across_threshold[Threshold == i & Status == "IC", Mean := mean(as.numeric(ecs[Localisation == "IC", Score]))]
  print(i)
}

ggplot() +
  # box plot of fecs
  geom_boxplot(data = fecs_across_threshold, 
               aes(x = Threshold, y= Mean, group = Threshold)) +
  # points of ecs
  geom_point(data = ecs_across_threshold,
             aes(x= Threshold, y= Mean, colour = Status)) +
  ylim(1, 8)
```
