## Supplementary material for "Predicting substrates for orphan Solute Carrier Proteins using multi-omics datasets": Code for generating results: NCI60extra_con_cytosis.R

```
library(limma)

tab = getGeneKEGGLinks(species="hsa")
tab$Symbol = mapIds(org.Hs.eg.db, tab$GeneID,
                       column="SYMBOL", keytype="ENTREZID")
tab = as.data.table(tab)
tab_cytosis = tab[PathwayID == "hsa04142" | PathwayID == "hsa04721"]
tab_cytosis = tab_cytosis[Symbol %in% SLC_annotation_2023$HGNC_symbol]
etki_cytosis = etki[HGNC_symbol %in% tab_cytosis$Symbol]

# initialise fake etki_cytosis collection
fake_etki_collection = vector("list", 100)
etki_gene_pool = unique(etki_cytosis$HGNC_symbol)
etki_metabolite_pool = unique(etki_cytosis$Formula)

for (sample_time in 1:100){
  # initialise fake etki_cytosis data table #
  fake_etki = setNames(data.table(matrix(ncol = ncol(etki_cytosis), nrow = nrow(etki_cytosis))), colnames(etki_cytosis))
  fake_etki[, Substrate := NULL]
  char.col = colnames(fake_etki)[1:4]
  num.col = colnames(fake_etki)[5:6]
  fake_etki[, (char.col) := lapply(.SD, as.character), .SDcols = char.col]
  fake_etki[, (num.col) := lapply(.SD, as.character), .SDcols = num.col]
  fake_etki[, c("HGNC_symbol", "Localisation") := as.list(etki_cytosis[, c("HGNC_symbol", "Localisation")])]
  fake_etki[, Status := rep("Fake", nrow(fake_etki))]
  # for each SLC, shuffle fake substrate formula #
  for (j in 1:nrow(fake_etki)){
    # produce fake substrate pools with considerations of adjacency #
    true_substrate = etki_cytosis[HGNC_symbol == fake_etki$HGNC_symbol[j]][, Formula]
    fake_substrate_pool = etki_metabolite_pool[!etki_metabolite_pool %in% true_substrate]
    fake_substrate = sample(fake_substrate_pool, 1)
    while (nrow(et[SYMBOL == fake_etki$HGNC_symbol[j] & Formula == fake_substrate][,NCI60extra_sample1:NCI60extra_sample2]) == 0){
      fake_substrate = sample(fake_substrate_pool, 1)
    }
    fake_etki[j, Formula := fake_substrate]
    fake_etki[j, c("NCI60extra_sample1", "NCI60extra_sample2") := 
                as.list(et[SYMBOL == fake_etki$HGNC_symbol[j] & Formula == fake_etki$Formula[j]][,NCI60extra_sample1:NCI60extra_sample2])]
    print(paste(sample_time, ":", j/nrow(fake_etki)))
  }
  fake_etki_collection[[sample_time]] = fake_etki
  #fwrite(fake_etki, paste("03 Results/08 Matrix Assembly/fake etki/Fake No", sample_time, ".csv", sep = ""))
  print(paste(sample_time, "done"))
}

ps_dist = na.omit(as.numeric(unlist(as.list(etki_cytosis[,6:7]))))
ps_dist = ps_dist[ps_dist > 0]
q = quantile(ps_dist, seq(0, 1, by = 0.1))

# etki confidence score (ecs) across threshold #
ecs_across_threshold = data.table(Threshold = seq(0, 1, by = 0.1), 
                                  Status = rep("ECS", length(seq(0, 1, by = 0.1))),
                                  Mean = vector("numeric", length(seq(0, 1, by = 0.1))))
for (i in 1:nrow(ecs_across_threshold)){
  ecs = ConfidenceScore(etki_cytosis, ecs_across_threshold[i, Threshold])
  ecs_across_threshold[i, Mean := mean(as.numeric(ecs[,Score]))]
  print(i)
}

# fake etki confidence score (fecs) across threshold #
fecs_across_threshold = data.table(Threshold = rep(seq(0, 1, by = 0.1), each = 100), 
                                   Status = rep("ECS", length(rep(seq(0, 1, by = 0.1), each = 100))),
                                   Mean = vector("numeric", length(rep(seq(0, 1, by = 0.1), each = 100))))
for (t in seq(0, 1, by = 0.1)){
  fecs_confidence_score = lapply(fake_etki_collection, function(x){ConfidenceScore(x, t)})
  fecs_confidence_score_mean = unlist(lapply(fecs_confidence_score, function(x){mean(as.numeric(x[,Score]))}))
  fecs_across_threshold[Threshold == t, Mean := fecs_confidence_score_mean]
  print(t)
}

# plot ecs with fecs across threshold #
ggplot() +
  # box plot of fecs
  geom_boxplot(data = fecs_across_threshold, 
               aes(x = Threshold, y= Mean, group = Threshold)) +
  # points of ecs
  geom_point(data = ecs_across_threshold,
             aes(x= Threshold, y= Mean),
             color = 'red') +
  ylim(0, 8)

# plot ecs with feces across threshold considering adjacency and localisation #
ecs_across_threshold = data.table(Threshold = rep(seq(0, 1, by = 0.1), each = 3), 
                                  Status = rep(c("PS", "PS-IC", "IC"), 11),
                                  Mean = vector("numeric", length(rep(seq(0, 1, by = 0.1), each = 3))))
for (i in unique(ecs_across_threshold$Threshold)){
  ecs = ConfidenceScore(etki_cytosis, i)
  ecs_across_threshold[Threshold == i & Status == "PS", Mean := mean(as.numeric(ecs[Localisation == "PS", Score]))]
  ecs_across_threshold[Threshold == i & Status == "PS-IC", Mean := mean(as.numeric(ecs[Localisation == "PS-IC", Score]))]
  ecs_across_threshold[Threshold == i & Status == "IC", Mean := mean(as.numeric(ecs[Localisation == "IC", Score]))]
  print(i)
}

ggplot() +
  # box plot of fecs
  geom_boxplot(data = fecs_across_threshold, 
               aes(x = Threshold, y= Mean, group = Threshold)) +
  # points of ecs
  geom_point(data = ecs_across_threshold,
             aes(x= Threshold, y= Mean, colour = Status)) +
  ylim(0, 8)
```
