## Supplementary material for "Predicting substrates for orphan Solute Carrier Proteins using multi-omics datasets": Code for generating results: Rho normalisation.R

```
##### 0 load libraries #####
library(data.table)

##### 1 read files #####
CCLE2019i <- fread("03 Results/01 Correlation Analysis/CCLE2019/CCLE2019 TvsM.csv")
NCI60i <- fread("03 Results/01 Correlation Analysis/NCI60a/NCI60a TvsM.csv")
CCL180i <- fread("03 Results/01 Correlation Analysis/CCL180/CCL180 TvsM.csv")

##### 2 normalise rho with pseudo Z-score #####
Rho_transform_metabolite <- function(dataset){
  dataset_metabolite = dataset[, unique(METABOLITE)]
  reference <- data.table(Metabolite = dataset_metabolite,
                          Mean = unlist(lapply(dataset_metabolite, function(x){dataset[METABOLITE == x, mean(abs(rho))]})),
                          SD = unlist(lapply(dataset_metabolite, function(x){dataset[METABOLITE == x, sd(abs(rho))]})))
  new_dataset <- dataset
  colnames(new_dataset)[4] <- "rho_normalised"
  for (i in 1:nrow(reference)){
    rho_normalise <- (abs(dataset[METABOLITE == reference[i, Metabolite], rho]) - reference[i, Mean])/reference[i, SD]
    new_dataset[METABOLITE == reference[i, Metabolite], rho_normalised := rho_normalise]
  }
  return(new_dataset)
}

CCLE2019i_normalised <- Rho_transform_metabolite(CCLE2019i)
NCI60i_normalised <- Rho_transform_metabolite(NCI60i)
CCL180i_normalised <- Rho_transform_metabolite(CCL180i)

fwrite(CCLE2019i_normalised, "03 Results/01 Correlation Analysis/Rho normalisation/CCLE2019_TvsM_normalised.csv")
fwrite(NCI60i_normalised, "03 Results/01 Correlation Analysis/Rho normalisation/NCI60_TvsM_normalised.csv")
fwrite(CCL180i_normalised, "03 Results/01 Correlation Analysis/Rho normalisation/CCL180_TvsM_normalised.csv")

##### 3 normalise based on each datasets #####
library(data.table)
CCLE2019i <- fread("03 Results/01 Correlation Analysis/CCLE2019/CCLE2019 TvsM.csv")
NCI60i <- fread("03 Results/01 Correlation Analysis/NCI60a/NCI60a TvsM.csv")
CCL180i <- fread("03 Results/01 Correlation Analysis/CCL180/CCL180 TvsM.csv")
Rho_transform <- function(dataset){
  set_mean = mean(dataset[,abs(rho)])
  set_deviation = sd(dataset[,abs(rho)])
  dataset[,rho_normalised := (abs(rho)-set_mean)/set_deviation]
  return(dataset)
}

CCLE2019i_normalised <- Rho_transform(CCLE2019i)
NCI60i_normalised <- Rho_transform(NCI60i)
CCL180i_normalised <- Rho_transform(CCL180i)

fwrite(CCLE2019i_normalised, "03 Results/01 Correlation Analysis/Rho normalisation/CCLE2019_TvsM_normalised (new).csv")
fwrite(NCI60i_normalised, "03 Results/01 Correlation Analysis/Rho normalisation/NCI60_TvsM_normalised (new).csv")
fwrite(CCL180i_normalised, "03 Results/01 Correlation Analysis/Rho normalisation/CCL180_TvsM_normalised (new).csv")
```
