## Supplementary material for "Predicting substrates for orphan Solute Carrier Proteins using multi-omics datasets": Code for generating results: CRISPR dependency with z-score.R

```
##### 0 load libraries #####
library(data.table)
library(tidyverse)

##### 1 Read files #####
CRISPR_GeneEffect <- fread("01 Raw Data/02 CRISPR dependency/CRISPRGeneEffect.csv") # DepMap Public 23Q2 Omics
CCLE2019_metabolomics <- fread("01 Raw Data/01 Correlation Analysis/01 CCLE2019/CCLE_metabolomics_20190502.csv")
Cell_line_metadata <- fread("01 Raw Data/02 CRISPR dependency/Model.csv")
SLC_annotation_2024 <- fread("01 Raw Data/00 SLC annotation 2024/SLC_transcript_20240321.csv", select = c(1:6))

##### 2 Pull SLC_GeneEffect out from CRISPR_GeneEffect
# re-format gene column and extract SLC
colnames(CRISPR_GeneEffect) <- gsub("\\s*\\([^\\)]+\\)","",colnames(CRISPR_GeneEffect))
SLC_GeneEffect <- CRISPR_GeneEffect[, c(TRUE, colnames(CRISPR_GeneEffect)[2:ncol(CRISPR_GeneEffect)] %in% SLC_annotation_2024$HGNC_symbol), with = FALSE] # 500 SLC found
SLC_annotation_2024$HGNC_symbol[!SLC_annotation_2024$HGNC_symbol %in% colnames(SLC_GeneEffect)]

# re-format cell line rows in CERES file and create SLC_GeneEffect and nonSLC_GeneEffect
ach_match <- c()
for (Ach in SLC_GeneEffect$ModelID){
  if (Ach %in% CCLE2019_metabolomics$DepMap_ID){
    ach_match <- c(ach_match, which(SLC_GeneEffect$ModelID == Ach))
  }
}
SLC_GeneEffect <- SLC_GeneEffect[ach_match, ] # 625 cell lines overlapped with CCLE 2019 metabolomics
SLC_GeneEffect <- dcast(melt(SLC_GeneEffect, id.vars = "ModelID"), variable ~ ModelID)
colnames(SLC_GeneEffect)[1] <- "SYMBOL"
fwrite(SLC_GeneEffect, "03 Results/02 CRISPR dependency/SLC_GeneEffect.csv")

# cleaning after step 2
rm(Ach)
rm(ach_match)

##### 3 transform CCLE 2019 metabolomics by z-score #####
# trim unfound cell lines in metadata
CCLE2019_metabolomics <- CCLE2019_metabolomics[!is.na(DepMap_ID)]
all(CCLE2019_metabolomics$DepMap_ID %in% Cell_line_metadata$ModelID) # TRUE
Cell_line_metadata <- Cell_line_metadata[Cell_line_metadata$ModelID %in% CCLE2019_metabolomics$DepMap_ID,]
# check tissue with only 1 cell lines in metadata
tissue_table <- Cell_line_metadata[, .N, by = SampleCollectionSite]
bad_tissue <- tissue_table[N == 1, SampleCollectionSite]
# 4 tissue, cervix, prostate, salivary_gland, unknown, only have 1 cell line linked with it
# remove these 4 tissue from both metadata and metabolomics
bad_cell_line <- Cell_line_metadata$ModelID[Cell_line_metadata$SampleCollectionSite %in% bad_tissue]
CCLE2019_metabolomics <- CCLE2019_metabolomics[-which(CCLE2019_metabolomics$DepMap_ID %in% bad_cell_line), ]
Cell_line_metadata <- Cell_line_metadata[-which(Cell_line_metadata$ModelID %in% bad_cell_line),]
tissue_table <- Cell_line_metadata[, .N, by = SampleCollectionSite]
# construct tissue information
tissue <- tissue_table[, SampleCollectionSite]
tissue_metabolomics_mean <- as.data.frame(matrix(nrow = length(tissue), ncol = ncol(CCLE2019_metabolomics)-1))
tissue_metabolomics_sd <- as.data.frame(matrix(nrow = length(tissue), ncol = ncol(CCLE2019_metabolomics)-1))

colnames(tissue_metabolomics_mean) <- c("Tissue", colnames(CCLE2019_metabolomics)[3:ncol(CCLE2019_metabolomics)])
colnames(tissue_metabolomics_sd) <- c("Tissue", colnames(CCLE2019_metabolomics)[3:ncol(CCLE2019_metabolomics)])
tissue_metabolomics_mean$Tissue <- tissue
tissue_metabolomics_sd$Tissue <- tissue

for (i in 1:length(tissue)){
  cell_line_same_tissue <- Cell_line_metadata$ModelID[Cell_line_metadata$SampleCollectionSite == tissue[i]]
  metabolomics_subset <- CCLE2019_metabolomics[CCLE2019_metabolomics$DepMap_ID %in% cell_line_same_tissue, ]
  tissue_metabolomics_mean[i,2:ncol(tissue_metabolomics_mean)] <- colMeans(metabolomics_subset[,3:ncol(metabolomics_subset)])
  tissue_metabolomics_sd[i,2:ncol(tissue_metabolomics_sd)] <- apply(metabolomics_subset[,3:ncol(metabolomics_subset)],2,sd)
}

CCLE2019_metabolomics_z_score <- as.data.frame(matrix(nrow = nrow(CCLE2019_metabolomics), ncol = ncol(CCLE2019_metabolomics)-1))
colnames(CCLE2019_metabolomics_z_score) <- colnames(CCLE2019_metabolomics)[2:ncol(CCLE2019_metabolomics)]
CCLE2019_metabolomics_z_score$DepMap_ID <- CCLE2019_metabolomics$DepMap_ID

CCLE2019_metabolomics <- as.data.frame(CCLE2019_metabolomics)
for (i in 1:nrow(CCLE2019_metabolomics)){
  tissue_category <- Cell_line_metadata$SampleCollectionSite[Cell_line_metadata$ModelID == CCLE2019_metabolomics$DepMap_ID[i]]
  this_tissue_mean <- tissue_metabolomics_mean[which(tissue_metabolomics_mean$Tissue == tissue_category), ]
  this_tissue_sd <- tissue_metabolomics_sd[which(tissue_metabolomics_sd$Tissue == tissue_category), ]
  z_score <- c()
  for (j in 3:ncol(CCLE2019_metabolomics)){
    z_score <- c(z_score, (CCLE2019_metabolomics[i, j] - this_tissue_mean[, j-1])/this_tissue_sd[, j-1])
  }
  CCLE2019_metabolomics_z_score[i, 2:ncol(CCLE2019_metabolomics_z_score)] <- z_score
  print(i)
}

##### 3 re-format CCLE2019 metabolomics to match with SLC_GeneEffect#####

# re-format cell line rows in CCLE2019 metabolomics
ach_match <- c()
for (Ach in CCLE2019_metabolomics_z_score$DepMap_ID){
  if (Ach %in% colnames(SLC_GeneEffect)){
    ach_match <- c(ach_match, which(CCLE2019_metabolomics_z_score$DepMap_ID == Ach))
  }
}

CCLE2019_metabolomics_z_score <- CCLE2019_metabolomics_z_score[ach_match, ]
CCLE2019_metabolomics_z_score <- as.data.table(CCLE2019_metabolomics_z_score)
CCLE2019_metabolomics_z_score <- dcast(melt(CCLE2019_metabolomics_z_score, id.vars = "DepMap_ID"), variable ~ DepMap_ID)
colnames(CCLE2019_metabolomics_z_score)[1] <- "Metabolite"

ach_match <- c()
for (Ach in colnames(CCLE2019_metabolomics_z_score)){
  if (Ach %in% colnames(SLC_GeneEffect)){
    ach_match <- c(ach_match, which(colnames(SLC_GeneEffect) == Ach))
  }
}

SLC_GeneEffect <- SLC_GeneEffect[, c(1, ach_match), with = FALSE]
all(colnames(SLC_GeneEffect)[2:ncol(SLC_GeneEffect)] == colnames(CCLE2019_metabolomics_z_score)[2:ncol(CCLE2019_metabolomics_z_score)]) #TRUE

fwrite(SLC_GeneEffect, "03 Results/02 CRISPR dependency/SLC_GeneEffect.csv")
fwrite(CCLE2019_metabolomics_z_score, "03 Results/02 CRISPR dependency/CCLE2019_metabolomics_z_score.csv")

##### 3 Wilcoxon test between top 20% (more negative) and bottom 20% (less negative) #####
# remove any SLC that have less 20 cell lines in top 20% and negative 20% after removing positive cell lines (i.e. less than 100 negative cell lines)
cell_line_checker = data.frame(SLC_name = SLC_GeneEffect$SYMBOL,
                               Negative_cell_line = unlist(lapply(1:nrow(SLC_GeneEffect), function(x){length(which(as.numeric(SLC_GeneEffect[x, 2:ncol(SLC_GeneEffect)])<0))})))
SLC_GeneEffect = SLC_GeneEffect[!(SLC_GeneEffect$SYMBOL %in% cell_line_checker$SLC_name[which(cell_line_checker$Negative_cell_line < 100)]),]

# Wilcoxon test
SLC_metab_wcx_table = as.data.frame(matrix(nrow = nrow(SLC_GeneEffect), ncol = nrow(CCLE2019_metabolomics_z_score)))
rownames(SLC_metab_wcx_table) = SLC_GeneEffect$SYMBOL
colnames(SLC_metab_wcx_table) = CCLE2019_metabolomics_z_score$Metabolite
SLC_metab_fc_table = SLC_metab_wcx_table

for (i in 1:nrow(SLC_GeneEffect)){
  # select out negative cell line
  SLC_GeneEffect_negative = SLC_GeneEffect[i, (which(as.numeric(SLC_GeneEffect[i,2:ncol(SLC_GeneEffect)])<0)+1), with = FALSE]
  # select out top 20% (more negative) and bottom 20% (less negative) cell lines
  Top20Per = colnames(SLC_GeneEffect_negative[, which(as.numeric(SLC_GeneEffect_negative) < quantile(as.numeric(SLC_GeneEffect_negative), probs = c(0.2))), with = FALSE])
  Bot20Per = colnames(SLC_GeneEffect_negative[, which(as.numeric(SLC_GeneEffect_negative) > quantile(as.numeric(SLC_GeneEffect_negative), probs = c(0.8))), with = FALSE])
  # match to metabolomics
  Top20Per_meta = CCLE2019_metabolomics_z_score[,colnames(CCLE2019_metabolomics_z_score) %in% Top20Per, with = FALSE]
  Bot20Per_meta = CCLE2019_metabolomics_z_score[,colnames(CCLE2019_metabolomics_z_score) %in% Bot20Per, with = FALSE]
  for (j in 1:nrow(CCLE2019_metabolomics_z_score)){
    SLC_metab_wcx_table[i,j] = wilcox.test(as.numeric(Top20Per_meta[j,]), as.numeric(Bot20Per_meta[j,]), exact = FALSE)$p.value
    SLC_metab_fc_table[i,j] = mean(as.numeric(Top20Per_meta[j,])) - mean(as.numeric(Bot20Per_meta[j,]))
  }
  print(i)
}

# adjust p-value
SLC_metab_wcx_table_padj = SLC_metab_wcx_table
for (i in 1:nrow(SLC_metab_wcx_table)){
  SLC_metab_wcx_table_padj[i,] = p.adjust(SLC_metab_wcx_table[i,], method = "BH", n = length(SLC_metab_wcx_table[i,]))
}

# melt data table
SLC_metab_wcx_table_padj_for_melt = SLC_metab_wcx_table_padj
SLC_metab_wcx_table_padj_for_melt$SYMBOL = rownames(SLC_metab_wcx_table_padj_for_melt)
SLC_metab_wcx_table_padj_melted = melt(SLC_metab_wcx_table_padj_for_melt, id.vars = "SYMBOL")

SLC_metab_fc_table_for_melt = SLC_metab_fc_table
SLC_metab_fc_table_for_melt$SYMBOL = rownames(SLC_metab_fc_table_for_melt)
SLC_metab_fc_table_melted = melt(SLC_metab_fc_table_for_melt, id.vars = "SYMBOL")

all(SLC_metab_fc_table_melted$SYMBOL == SLC_metab_wcx_table_padj_melted$SYMBOL) # TRUE
all(SLC_metab_fc_table_melted$variable == SLC_metab_wcx_table_padj_melted$variable) # TRUE

colnames(SLC_metab_wcx_table_padj_melted) = c("SYMBOL", "METABOLITE", "padj")
colnames(SLC_metab_fc_table_melted) = c("SYMBOL", "METABOLITE", "FC")
SLC_dependency_wcx_table = data.frame(SYMBOL = SLC_metab_wcx_table_padj_melted$SYMBOL, METABOLITE = SLC_metab_wcx_table_padj_melted$METABOLITE, 
                                      padj = SLC_metab_wcx_table_padj_melted$padj, FC = SLC_metab_fc_table_melted$FC)
fwrite(SLC_dependency_wcx_table, "03 Results/02 CRISPR dependency/z-score metabolite dependency wilcoxon test.csv")


##### 4 Benchmarking with known interaction #####
# known interaction
CCLE_ki = na.omit(ki[,c(1,2,7)])
CCLE_ki = CCLE_ki[CCLE_ki$HGNC_symbol %in% rownames(SLC_metab_wcx_table_padj),] # 41 genes from CCLE_ki not in trimmed dependency genes
all(CCLE_ki$CCLE2019 %in% colnames(SLC_metab_wcx_table_padj)) # TRUE
CCLE_ki_wcx_padj = c()
CCLE_ki_fc = c()
for (i in 1:nrow(CCLE_ki)){
  CCLE_ki_wcx_padj = c(CCLE_ki_wcx_padj, SLC_metab_wcx_table_padj[which(rownames(SLC_metab_wcx_table_padj) == CCLE_ki$HGNC_symbol[i]),
                                                                  which(colnames(SLC_metab_wcx_table_padj) == CCLE_ki$CCLE2019[i])])
  CCLE_ki_fc = c(CCLE_ki_fc, SLC_metab_fc_table[which(rownames(SLC_metab_fc_table) == CCLE_ki$HGNC_symbol[i]),
                                                which(colnames(SLC_metab_fc_table) == CCLE_ki$CCLE2019[i])])
}
CCLE_ki$wcx_padj = CCLE_ki_wcx_padj
CCLE_ki$fc = CCLE_ki_fc
#fwrite(CCLE_ki, "3 Results/07 CRISPR dependency/z-score metabolite CCLE ki.csv")

# melt data table
SLC_metab_wcx_table_padj_for_melt = SLC_metab_wcx_table_padj
SLC_metab_wcx_table_padj_for_melt$SYMBOL = rownames(SLC_metab_wcx_table_padj_for_melt)
SLC_metab_wcx_table_padj_melted = melt(SLC_metab_wcx_table_padj_for_melt, id.vars = "SYMBOL")
SLC_metab_fc_table_for_melt = SLC_metab_fc_table
SLC_metab_fc_table_for_melt$SYMBOL = rownames(SLC_metab_fc_table_for_melt)
SLC_metab_fc_table_melted = melt(SLC_metab_fc_table_for_melt, id.vars = "SYMBOL")
all(SLC_metab_fc_table_melted$SYMBOL == SLC_metab_wcx_table_padj_melted$SYMBOL) # TRUE
all(SLC_metab_fc_table_melted$variable == SLC_metab_wcx_table_padj_melted$variable) # TRUE
colnames(SLC_metab_wcx_table_padj_melted) = c("SYMBOL", "METABOLITE", "padj")
colnames(SLC_metab_fc_table_melted) = c("SYMBOL", "METABOLITE", "FC")
SLC_dependency_wcx_table = data.frame(SYMBOL = SLC_metab_wcx_table_padj_melted$SYMBOL, METABOLITE = SLC_metab_wcx_table_padj_melted$METABOLITE, 
                                      padj = SLC_metab_wcx_table_padj_melted$padj, FC = SLC_metab_fc_table_melted$FC)
#fwrite(SLC_dependency_wcx_table, "3 Results/07 CRISPR dependency/z-score metabolite dependency wilcoxon test.csv")

# fake interaction
fake_ki = function(ki, interaction){
  colnames(ki) = c("HGNC_symbol", "Localisation", "metabolite", "padj", "fc")
  ki_metabolite_pool = as.character(as.data.frame(table(ki$metabolite))$Var1)
  fake_ki = ki[,c(1,2)]
  fake_ki_metabolite = c()
  for (i in 1:nrow(fake_ki)){
    specific_interaction = ki[which(ki$HGNC_symbol == ki$HGNC_symbol[i]), ] # find specific interacting metabolites for chosen SLC
    random_metabolite = sample(ki_metabolite_pool[!(ki_metabolite_pool %in% specific_interaction$metabolite)], 1) # randomly select one from nonspecific pool
    fake_ki_metabolite = c(fake_ki_metabolite, random_metabolite)
  }
  fake_ki$metabolite = fake_ki_metabolite
  padj = unlist(lapply(1:nrow(fake_ki), function(j){interaction$padj[which(interaction$METABOLITE == fake_ki$metabolite[j] & interaction$SYMBOL == fake_ki$HGNC_symbol[j])]}))
  FC = unlist(lapply(1:nrow(fake_ki), function(j){interaction$FC[which(interaction$METABOLITE == fake_ki$metabolite[j] & interaction$SYMBOL == fake_ki$HGNC_symbol[j])]}))
  fake_ki$padj = padj
  fake_ki$fc = FC
  return(fake_ki)
}

fake_ki_collection = vector(mode = "list", length = 100)
for (sample_time in 1:100){
  fake_ki_collection[[sample_time]] = fake_ki(CCLE_ki, SLC_dependency_wcx_table)
  print(sample_time)
}

# volcano plot comparing known interaction and fake interaction
# plot all fake interactions
ki_volcano = fake_ki_collection[[1]]
for(i in 2:length(fake_ki_collection)){
  ki_volcano = rbind(ki_volcano, fake_ki_collection[[i]])
  print(i)
}
ki_volcano$type = rep("fake interaction", nrow(ki_volcano))

CCLE_ki_volcano = CCLE_ki
CCLE_ki_volcano$type = rep("known interaction", nrow(CCLE_ki_volcano))
colnames(CCLE_ki_volcano) = colnames(ki_volcano)
ki_volcano = rbind(ki_volcano, CCLE_ki_volcano)

ggplot(data=ki_volcano, aes(x=fc, y=-log10(padj), col = type)) + 
  geom_point() + 
  theme_minimal() + 
  scale_color_manual(values=c("grey", "black"))


# fractions of interaction recovery
fake_significant = unlist(lapply(1:100, function(x){length(which(fake_ki_collection[[x]][,4]<0.05))/length(fake_ki_collection[[x]][,4])}))
boxplot(fake_significant, ylim = c(0, 0.1))
abline(h = length(which(CCLE_ki_wcx_padj<0.05))/length(CCLE_ki_wcx_padj))

benchmark = data.table(fraction = c(fake_significant, length(which(CCLE_ki_wcx_padj<0.05))/length(CCLE_ki_wcx_padj)),
                       class = c(rep("random", 100), "true"))
fwrite(benchmark, "benchmark.csv")

# best p-value cutoff to maximise the recovery
threshold = seq(0,0.4,0.01)
best_check = data.table(X = threshold,
                        Y = rep(0, 41),
                        Z = rep(0, 41),
                        Q = rep(0, 41))
for (i in 1:length(threshold)){
  y = length(which(CCLE_ki_wcx_padj<threshold[i]))/length(CCLE_ki_wcx_padj)
  z = mean(unlist(lapply(1:100, function(x){length(which(fake_ki_collection[[x]][,4]<threshold[i]))/length(fake_ki_collection[[x]][,4])})))
  best_check[i, Y := y]
  best_check[i, Z := z]
  best_check[i, Q := y-z]
}

plot(best_check$X,best_check$Q, pch=18, col = "#69b3a2", xaxt="n")
axis(1, at = seq(0, 1, 0.1))
```
