## Supplementary material for "Predicting substrates for orphan Solute Carrier Proteins using multi-omics datasets": Code for generating results: Benchmark.R

```
##### 0 load libraries #####
library(data.table)
library(tidyverse)

##### 1 read files #####
# translation book (tb)
et <- fread("03 Results/03 Table alignment/Evidence table (normalised rho new).csv")
ki <- fread("01 Raw Data/00 SLC annotation 2024/known interaction/Known interaction formula 20240321.csv")[1:841]

##### 2 assemble etki #####
# remove substrates not covered by evidence table formula and derivatives
ki <- ki[!is.na(Formula)][Status != "Derivative"]

etki = ki
colnames(etki)[1] = "SYMBOL"
etki$NCI60 = vector("numeric", nrow(etki))
etki$CCLE2019 = vector("numeric", nrow(etki))
etki$CCL180 = vector("numeric", nrow(etki))
etki$CRISPRDEP = vector("numeric", nrow(etki))

for (i in 1:nrow(etki)){
  if (etki$SYMBOL[i] %in% unique(et$SYMBOL)){
    etki[i, c("NCI60", "CCLE2019", "CCL180", "CRISPRDEP") :=
           as.list(et[SYMBOL == etki$SYMBOL[i] & Formula == etki$Formula[i]][,NCI60:CRISPRDEP])]
  }else{
    etki[i, c("NCI60", "CCLE2019", "CCL180", "CRISPRDEP") := list(NA, NA, NA, NA)]
  }
}
etki = etki[rowSums(is.na(etki[,8:11]))<4]

##### 3 benckmark individual etki #####
benchmark = function(dataset, source){
  fake_etki_collection = vector("list", 100)
  etki_gene_pool = unique(dataset$SYMBOL)
  etki_metabolite_pool = unique(dataset$Formula)
  
  for (sample_time in 1:100){
    # initialise fake dataset #
    fake_etki = setNames(data.table(matrix(ncol = ncol(dataset), nrow = nrow(dataset))), colnames(dataset))
    fake_etki[, Substrate := NULL]
    char.col = colnames(fake_etki)[1:3]
    num.col = colnames(fake_etki)[4]
    fake_etki[, (char.col) := lapply(.SD, as.character), .SDcols = char.col]
    fake_etki[, (num.col) := lapply(.SD, as.character), .SDcols = num.col]
    fake_etki[, c("SYMBOL", "Gene_type") := as.list(dataset[, c("SYMBOL", "Gene_type")])]
    # for each SLC, shuffle fake substrate formula #
    for (j in 1:nrow(fake_etki)){
      true_substrate = dataset[SYMBOL == fake_etki$SYMBOL[j], Formula]
      fake_substrate_pool = etki_metabolite_pool[!etki_metabolite_pool %in% true_substrate]
      fake_substrate = sample(fake_substrate_pool, 1)
      while (nrow(et[SYMBOL == fake_etki$SYMBOL[j] & Formula == fake_substrate][,NCI60:CRISPRDEP]) == 0){
        fake_substrate = sample(fake_substrate_pool, 1)
      }
      fake_etki[j, Formula := fake_substrate]
      fake_etki[j, (source) := 
                  et[SYMBOL == fake_etki$SYMBOL[j] & Formula == fake_etki$Formula[j], source, with = FALSE]]
      print(paste(sample_time, ":", round(j/nrow(fake_etki),2)))
    }
    fake_etki_collection[[sample_time]] = fake_etki
    print(paste(sample_time, "done"))
  }
  return(fake_etki_collection)
}

# individual etki
NCI60_etki = etki[!is.na(NCI60)][, c(1,2,4,5,8)]
CCLE2019_etki = etki[!is.na(CCLE2019)][, c(1,2,4,5,9)]
CCL180_etki = etki[!is.na(CCL180)][, c(1,2,4,5,10)]
CRISPR_etki = etki[!is.na(CRISPRDEP)][, c(1,2,4,5,11)]

# generate individual simulation
NCI60_simulated_random = benchmark(NCI60_etki, "NCI60")
CCLE2019_simulated_random = benchmark(CCLE2019_etki, "CCLE2019")
CCL180_simulated_random = benchmark(CCL180_etki, "CCL180")
CRISPR_simulated_random = benchmark(CRISPR_etki, "CRISPRDEP")

##### 4 report on benchmark #####
# Correlation analysis
NCI60_report = data.table(
  Data = rep("NCI60", 101),
  Type = c("Known", rep("Simulated", 100)),
  Mean = c(NCI60_etki[, mean(NCI60)], 
           unlist(lapply(NCI60_simulated_random, function(x){x[, mean(as.numeric(NCI60))]})))
)

CCLE2019_report = data.table(
  Data = rep("CCLE2019", 101),
  Type = c("Known", rep("Simulated", 100)),
  Mean = c(CCLE2019_etki[, mean(CCLE2019)], 
           unlist(lapply(CCLE2019_simulated_random, function(x){x[, mean(as.numeric(CCLE2019))]})))
)

CCL180_report = data.table(
  Data = rep("CCL180", 101),
  Type = c("Known", rep("Simulated", 100)),
  Mean = c(CCL180_etki[, mean(CCL180)], 
           unlist(lapply(CCL180_simulated_random, function(x){x[, mean(as.numeric(CCL180))]})))
)

benchmark_report = rbind(NCI60_report, CCLE2019_report, CCL180_report)
fwrite(benchmark_report, "03 Results/03 Table alignment/Correlation analysis benchmark report (new).csv")

# CRISPR Dependency
CRISPR_report = data.table(
  Type = c("Known", rep("Simulated", 100)),
  Mean = c(CRISPR_etki[, mean(CRISPRDEP)], 
           unlist(lapply(CRISPR_simulated_random, function(x){x[, mean(as.numeric(CRISPRDEP))]}))),
  Fraction = c(CRISPR_etki[CRISPRDEP < 0.05, .N]/CRISPR_etki[,.N], 
               unlist(lapply(CRISPR_simulated_random, function(x){x[CRISPRDEP < 0.05, .N]/x[,.N]})))
)
fwrite(CRISPR_report, "03 Results/03 Table alignment/CRISPRDEP benchmark report.csv")

# CRISPR Dependency recovery across alpha
known_recovery = data.table(Type = rep("Known", 100),
                            AlphaCutoff = seq(0.001, 0.1, 0.001),
                            Fraction_recovery = vector("numeric", 100)
)
for (i in 1:nrow(known_recovery)){
  nr = CRISPR_etki[as.numeric(CRISPRDEP) < known_recovery[i, AlphaCutoff], .N]/CRISPR_etki[,.N]
  known_recovery[i, Fraction_recovery := nr]
}

simulated_recovery = data.table(Type = rep("Simulated", 10000),
                                AlphaCutoff = rep(seq(0.001, 0.1, 0.001), each = 100),
                                Fraction_recovery = vector("numeric", 10000)
)
for (i in seq(0.001, 0.1, 0.001)){
  nr = unlist(lapply(CRISPR_simulated_random, function(x){x[CRISPRDEP < i, .N]/x[,.N]}))
  simulated_recovery[AlphaCutoff == i, Fraction_recovery := nr]
}
recovery = rbind(known_recovery, simulated_recovery)
fwrite(recovery, "03 Results/03 Table alignment/CRISPRDEP_FractionRecoveryAcrossAlpha.csv")
```
