## Supplementary material for "Predicting substrates for orphan Solute Carrier Proteins using multi-omics datasets": Code for generating results: Cell line alignment info.R

```
##### 0 load libraries #####
library(data.table)
library(tidyverse)

##### 1 Concentration concordance #####
#### 1.1 read files ####
CCLE2019_metabolomics = fread("01 Raw Data/01 Predictive Scores/01 CCLE2019/CCLE_metabolomics_20190502.csv")
NCI60_metabolomics = fread("01 Raw Data/01 Predictive Scores/02 NCI60a/NCI60 metabolomics.csv")
CCL180_metabolomics = fread("01 Raw Data/01 Predictive Scores/04 CCL180/CCL180 metabolomics.csv")
model = fread("01 Raw Data/01 Predictive Scores/DepMap Model.csv")
tb = fread("01 Raw Data/00 SLC annotation 2023/book construction/step 3 3 matrices book full.csv")

#### 1.2 unify the cell line names across metabolomics (use ACH code) ####
# updating CCLE2019 information
# CCLE2019: OC315_OVARY - NA
CCLE2019_metabolomics[CCLE_ID == "OC315_OVARY", DepMap_ID := "ACH-001144"]
# CCLE2019: removing HCC1588_LUNG - ACH-001078 as the cell line was from ACH-000007
# evidence post [https://forum.depmap.org/t/important-update-fingerprinting-data-corrections-21q3-release/810]
CCLE2019_metabolomics = CCLE2019_metabolomics[DepMap_ID != "ACH-001078"]
CCLE2019_metabolomics = CCLE2019_metabolomics[, -1, with = FALSE]
CCLE2019_metabolomics = dcast(melt(CCLE2019_metabolomics, id.vars = "DepMap_ID"), variable ~ DepMap_ID)
colnames(CCLE2019_metabolomics)[1] = "metabolite"

# updating NCI60 information
colnames(NCI60_metabolomics) = c(
  "metabolite",      "ACH-000649",      "ACH-000555",      "ACH-000681",      "ACH-000046",      "ACH-000288",         
  "ACH-000433",      "ACH-001738",      "ACH-001039",      "ACH-000979",      "ACH-000706",      "ACH-001081",       
  "ACH-000971",      "ACH-000997",      "ACH-000002",      "ACH-000861",      "ACH-000825",      "ACH-000148",        
  "ACH-000552",      "ACH-000966",      "ACH-000551",      "ACH-000969",      "ACH-000750",      "ACH-002159",            
  "ACH-000477",      "ACH-000019",      "ACH-000768",      "MDA-MB-435",      "ACH-001127",      "ACH-000367",       
  "ACH-000900",      "ACH-002172",      "ACH-000463",      "ACH-000343",      "NCI/ADR-RES",     "ACH-000001",       
  "ACH-000617",      "ACH-001151",      "ACH-000696",      "ACH-000090",      "ACH-000817",      "ACH-002195",        
  "ACH-000655",      "ACH-000376",      "ACH-000273",      "ACH-000615",      "ACH-000730",      "ACH-000811",        
  "ACH-002199",      "ACH-001198",      "ACH-000504",      "SR",              "ACH-000651",      "ACH-000147",          
  "ACH-001208",      "ACH-000232",      "ACH-000579",      "ACH-000425",      "ACH-000428"  
)

# updating CCL180 information
colnames(CCL180_metabolomics) = c(
  "metabolite",   "ACH-002319", "ACH-000905",  "ACH-000896",  "ACH-000411",  "ACH-000649",  "977",         
  "ACH-000757",   "ACH-000555", "ACH-000681",  "ACH-000429",  "ACH-000046",  "ACH-000222",  "ACH-000536",        
  "ACH-000288",   "ACH-000802", "ACH-000535",  "ACH-000433",  "ACH-000856",  "ACH-000511",  "ACH-000392",       
  "ACH-000264",   "ACH-001738", "ACH-000867",  "ACH-001039",  "ACH-000662",  "ACH-000257",  "CORL88A",     
  "ACH-000530",   "ACH-000698", "ACH-000979",  "ACH-000258",  "ACH-000563",  "ACH-000706",  "ACH-000585",    
  "ACH-000843",   "ACH-000374", "ACH-000111",  "ACH-000699",  "ACH-000878",  "ACH-000624",  "ACH-000223",     
  "ACH-000276",   "ACH-000668", "ACH-000971",  "ACH-000997",  "ACH-000643",  "ACH-000002",  "ACH-000861",       
  "ACH-000825",   "ACH-000270", "ACH-000148",  "ACH-000547",  "ACH-000724",  "ACH-000552",  "HX147",       
  "ACH-000966",   "ACH-000396", "ACH-000551",  "ACH-000969",  "ACH-000858",  "ACH-000265",  "ACH-000486",      
  "ACH-000769",   "ACH-000750", "ACH-000390",  "ACH-002159",  "ACH-000477",  "ACH-000019",  "ACH-000621",    
  "ACH-000768",   "ACH-000884", "ACH-000573",  "ACH-000849",  "MGHU3",       "ACH-000601",  "ACH-001127",       
  "ACH-000866",   "ACH-000980", "ACH-000510",  "ACH-000327",  "ACH-000589",  "ACH-000506",  "ACH-000892",    
  "ACH-000035",   "ACH-000893", "ACH-000448",  "ACH-000021",  "ACH-000747",  "ACH-000675",  "ACH-000282",    
  "ACH-000496",   "ACH-000888", "ACH-000733",  "ACH-000894",  "ACH-000587",  "ACH-001137",  "ACH-000886",    
  "ACH-000521",   "ACH-000590", "ACH-000451",  "ACH-000290",  "ACH-000311",  "ACH-000785",  "ACH-001363",    
  "ACH-000481",   "ACH-000525", "ACH-000924",  "ACH-000447",  "ACH-000367",  "ACH-000912",  "ACH-000718",    
  "ACH-000900",   "ACH-000951", "ACH-000875",  "ACH-000186",  "ACH-001075",  "ACH-000837",  "ACH-000109",    
  "ACH-001364",   "ACH-000860", "ACH-000638",  "ACH-000800",  "ACH-000463",  "NCIH460DNP53","ACH-000871",     
  "ACH-000395",   "ACH-000343", "ACH-000767",  "ACH-000628",  "ACH-000378",  "ACH-000853",  "ACH-000775",     
  "ACH-000789",   "ACH-000355", "ACH-000416",  "ACH-000001",  "ACH-000617",  "ACH-001151",  "ACH-000696",      
  "ACH-000139",   "ACH-000235", "ACH-000417",  "ACH-000164",  "PC14PE6",     "ACH-000090",  "ACH-000779",         
  "ACH-000261",   "ACH-000442", "ACH-000817",  "ACH-000242",  "ACH-002195",  "ACH-000670",  "ACH-000839",      
  "ACH-000655",   "ACH-000376", "ACH-000273",  "ACH-000790",  "ACH-000309",  "ACH-001190",  "ACH-000730",      
  "ACH-000665",   "ACH-000811", "ACH-002199",  "ACH-000504",  "SR8714",      "ACH-000114",  "ACH-001390",    
  "ACH-001398",   "ACH-000677", "ACH-000651",  "ACH-000384",  "ACH-000669",  "ACH-000018",  "ACH-000147",        
  "ACH-000579",   "ACH-000425", "ACH-001417",  "ACH-000522",  "ACH-000428",  "ACH-000545"  
)
```
