## Supplementary material for "Predicting substrates for orphan Solute Carrier Proteins using multi-omics datasets": Code for generating results: Concordance.R

```
##### 0 load libraries #####
library(data.table)
library(tidyverse)
#### 3.1 read files ####
et = fread("03 Results/03 Table alignment/Evidence table (rho).csv")

#### 3.2 mutual Spearman concordance ####
### CCLE2019 and NCI60 ###
et_1 <- et[rowSums(is.na(et[,3:4])) == 0][, !5]
length(unique(et_1$Formula))
length(unique(et_1$SYMBOL))
cor.test(et_1$CCLE2019, et_1$NCI60, method = "spearman", exact = F)$p.value
cor.test(et_1$CCLE2019, et_1$NCI60, method = "spearman", exact = F)$estimate

### CCLE2019 and CCL180 ###
et_2 = et[rowSums(is.na(et[,4:5])) == 0][, !3]
length(unique(et_2$Formula))
length(unique(et_2$SYMBOL))
cor.test(et_2$CCLE2019, et_2$CCL180, method = "spearman", exact = F)$p.value
cor.test(et_2$CCLE2019, et_2$CCL180, method = "spearman", exact = F)$estimate

### NCI60 and CCL180 ###
et_3 = et[rowSums(is.na(et[,c(3,5)])) == 0][, !4]
length(unique(et_3$Formula))
length(unique(et_3$SYMBOL))
cor.test(et_3$NCI60, et_3$CCL180, method = "spearman", exact = F)$p.value
cor.test(et_3$NCI60, et_3$CCL180, method = "spearman", exact = F)$estimate

#### 3.3 all Spearman concordance ####
et_oa = et[rowSums(is.na(et[,3:5])) == 0]
length(unique(et_oa$Formula))
length(unique(et_oa$SYMBOL))
cor.test(et_oa$CCLE2019, et_oa$NCI60, method = "spearman", exact = F)
cor.test(et_oa$CCLE2019, et_oa$CCL180, method = "spearman", exact = F)
cor.test(et_oa$NCI60, et_oa$CCL180, method = "spearman", exact = F)
```
