## Supplementary material for "Predicting substrates for orphan Solute Carrier Proteins using multi-omics datasets": Code for generating results: Name conversion and evidence table (normalised rho) generation.R

```
##### 0 load libraries #####
library(data.table)

##### 1 read files #####
# translation book (tb)
tb = fread("01 Raw Data/00 SLC annotation 2024/book construction/Translation book (3 Matrix).csv")
CCLE2019_SLC_interaction = fread("03 Results/01 Correlation Analysis/Rho normalisation/CCLE2019_TvsM_normalised (new).csv")
NCI60_SLC_interaction = fread("03 Results/01 Correlation Analysis/Rho normalisation/NCI60_TvsM_normalised (new).csv")
CCL180_SLC_interaction = fread("03 Results/01 Correlation Analysis/Rho normalisation/CCL180_TvsM_normalised (new).csv")
CRISPR_dependency = fread("03 Results/02 CRISPR dependency/z-score metabolite dependency wilcoxon test.csv")

##### 2 define functions #####

# ConvertNames #
ConvertNames = function(name, source){
  # Process source input #
  if (is.null(source)){
    stop("Source of metabolite name required!")
  }
   else if (source %in% colnames(tb)){
    # Process name input #
     if (is.null(name)){
       stop("Name of metabolite required!")
     }else if (name %in% tb[,get(source)]){
       ans = tb[get(source) == name]
     }else{
       stop("Name entered not recognised!")
     }
  }else{
    stop("Source entered not recognised!")
  }
  
  # return ans as result #
  return(ans)
}
# BuildEvidenceTable #
BuildEvidenceTable = function(gene, formula){
  # build table #
  ans = data.table(SYMBOL = gene, Formula = formula, NCI60 = NA, CCLE2019 = NA, CCL180 = NA)
  # process formula #
  line = try(ConvertNames(formula, "Formula"), silent = TRUE)
  if (grepl("Error", line[[1]]) == TRUE){
    stop("Error in converting formula!")
  }else{
    compound_info = ConvertNames(formula, "Formula")
  }
  info_source = c("NCI60", "CCLE2019", "CCL180")
  for (i in 1:length(info_source)){
    if (is.na(compound_info[,get(info_source[i])]) == TRUE){
      ans[, info_source[i]:=NA]
    }else{
      ans[, info_source[i]:=get(paste(info_source[i], "_SLC_interaction", sep = ""))
          [SYMBOL == gene & METABOLITE == compound_info[,get(info_source[i])]][,rho_normalised]]
    }
  }
  return(ans)
}
# Outersect #
outersect = function(x, y) {
  sort(c(x[!x%in%y],
         y[!y%in%x]))
}

##### 3 assemble predictive evidence table #####

# assemble evidence table (et) #
specific_SLC_pool = outersect(unique(CCLE2019_SLC_interaction$SYMBOL), unique(NCI60_SLC_interaction$SYMBOL))
overlapped_SLC_pool = intersect(unique(CCLE2019_SLC_interaction$SYMBOL), unique(NCI60_SLC_interaction$SYMBOL))

# overlapping SLC #
et = data.table(SYMBOL = character(), Formula = character(), NCI60 = numeric(), CCLE2019 = numeric(), CCL180 = numeric())
for (i in 1:length(overlapped_SLC_pool)){
  for (j in 1:length(tb$Formula)){
    temp = BuildEvidenceTable(overlapped_SLC_pool[i], tb$Formula[j])
    et = rbind(et, temp)
  }
  print(i)
}

# specific SLC #
"SLC25A53" %in% unique(NCI60_SLC_interaction$SYMBOL)
"SLC5A3" %in% unique(NCI60_SLC_interaction$SYMBOL)
"SLC6A14" %in% unique(NCI60_SLC_interaction$SYMBOL)
"SLCO1B7" %in% unique(CCLE2019_SLC_interaction$SYMBOL)

et_specific = data.table(SYMBOL = character(), Formula = character(), NCI60 = numeric(), CCLE2019 = numeric(), CCL180 = numeric())

# specific SLC - specific to NCI60 #
for (i in 1:(length(specific_SLC_pool)-1)){
  for (j in 1:length(tb$Formula)){
    temp = data.table(SYMBOL = specific_SLC_pool[i], Formula = tb$Formula[j], NCI60 = NA, CCLE2019 = NA, CCL180 = NA)
    compound_info = ConvertNames(tb$Formula[j], "Formula")
    if (is.na(compound_info[,NCI60] == TRUE)){
      temp[, NCI60:=NA]
    }else{
      temp[, NCI60:=NCI60_SLC_interaction[SYMBOL == specific_SLC_pool[i] & METABOLITE == compound_info[,NCI60]][,rho_normalised]]
    }
    et_specific = rbind(et_specific, temp)
    print(paste(i, ":", j/length(tb$Formula)))
  }
}

# merge et_specific to et #
et_specific = et_specific[rowSums(is.na(et_specific[,3:5]))<3] # remove all interaction with NA in all three matrices
et = rbind(et, et_specific)

# merge with CRISPR dependency wilcoxon test padj
et$CRISPRDEP = vector('numeric',nrow(et))
for (i in 1:nrow(et)){
  compound_info = ConvertNames(et$Formula[i], "Formula")
  if (is.na(compound_info[,CCLE2019])){
    et[i, CRISPRDEP:= NA]
    print(i/nrow(et))
  }else if (!et$SYMBOL[i] %in% unique(CRISPR_dependency$SYMBOL)){
    et[i, CRISPRDEP:= NA]
    print(i/nrow(et))
  }else{
    et[i, CRISPRDEP:= CRISPR_dependency[SYMBOL == et$SYMBOL[i] & METABOLITE == compound_info[,CCLE2019]][,padj]]
  }
  print(i/nrow(et))
}

fwrite(et, "03 Results/03 Table alignment/Evidence table (normalised rho new).csv")
```
