## Supplementary material for "Predicting substrates for orphan Solute Carrier Proteins using multi-omics datasets": Code for generating results: Adjacency matrix via MetaboSignal.R

```
# load libraries
library(MetaboSignal)
library(data.table)
library(progress)

# get human pathways
human_pathways = MS_getPathIds(organism_code = "hsa")
human_metabolic_pathways = human_pathways[human_pathways[, "Path_type"] == "metabolic",]
human_signaling_pathways = human_pathways[human_pathways[, "Path_type"] == "signaling",]

# build reaction network
hmn = MS_reactionNetwork(human_metabolic_pathways[, "Path_id"]) # human metabolic network

# calculate node betweenness (centrality)
# NodeBW = MS_nodeBW(hmn, "out", normalized = TRUE)
# NodeBW_compound = NodeBW[grep("cpd", names(NodeBW))]
# NodeBW_data = data.frame(Compound = substr(names(NodeBW_compound), 5, nchar(names(NodeBW_compound))), NodeBW = as.numeric(NodeBW_compound))

# build adjacency matrix
compound = unique(hmn[,"source"][grep("cpd", hmn[,"source"])])
AM = matrix(nrow = length(compound), ncol = length(compound))
colnames(AM) = compound
rownames(AM) = compound
pb = progress_bar$new(format = "(:spin) [:bar] :percent [Elapsed time: :elapsedfull || Estimated time remaining: :eta]",
                      total = length(compound),
                      complete = "=",   # Completion bar character
                      incomplete = "-", # Incomplete bar character
                      current = ">",    # Current bar character
                      clear = FALSE,    # If TRUE, clears the bar when finish
                      width = 100)      # Width of the progress bar
for (i in 1:length(compound)){
  pb$tick()
  for (j in 1:length(compound)){
    if (i == j){
      AM[i,j] = 0
    }else{
      distance = MS_shortestPaths(hmn, compound[i], compound[j], mode = "out", type = "first")
      if (is.null(distance)){
        AM[i,j] = Inf
      }else{
        AM[i,j] = (length(distance)-1)/2
      }
    }
  }
}

MS_exportCytoscape(hmn, "hsa")

AM_for_output = as.data.table(AM)
AM_for_output[, COMPOUND := compound]
setcolorder(AM_for_output, "COMPOUND")

fwrite(AM_for_output, "03 Results/04 Adjacency/Metabolie Adjacency Matrix.csv")
```
