## Supplementary material for "Predicting substrates for orphan Solute Carrier Proteins using multi-omics datasets": Code for generating results: Adjacency matrix via Recon3D.R

```
# Load required packages
library(R.matlab)
library(igraph)

##### Step 1: represent recon 3D as a graph #####
# Load Recon3D data from .mat file
recon3d_data <- readMat("01 Raw Data/04 Adjacency/Recon3D_301.mat")

# Extract metabolites and reactions
mets <- unlist(recon3d_data[["Recon3D"]][,,1]["mets"])
rxns <- unlist(recon3d_data[["Recon3D"]][,,1]["rxns"])
S <- recon3d_data[["Recon3D"]][,,1][["S"]]

# Create empty graph object
metabolic_graph <- make_empty_graph()

# Add metabolite nodes to the graph
for (metabolite_id in mets) {
  metabolic_graph <- add_vertices(metabolic_graph, nv = 1, name = metabolite_id)
}

# Add reaction edges to the graph
for (i in 1:ncol(S)) {
  reactants <- which(S[, i] < 0)
  products <- which(S[, i] > 0)
  for (reactant_index in reactants) {
    for (product_index in products) {
      metabolic_graph <- add_edges(metabolic_graph, c(mets[reactant_index], mets[product_index]))
    }
  }
}

##### Step 2: remove vertex with super-high connectivities #####
#reconMetabolite = fread("01 Raw Data/04 Adjacency/reconMetabolite.tsv")
# degree centrality #
#metabolic_degree = degree(metabolic_graph)
#metabolic_degree[order(metabolic_degree, decreasing = T)][1:400]
# betweenness centrality #
#metabolic_betweenness = betweenness(metabolic_graph)
#metabolic_betweenness[order(metabolic_betweenness, decreasing = T)][1:400]

#high_centrality = intersect(names(metabolic_degree[order(metabolic_degree, decreasing = T)][1:400]),
#                            names(metabolic_betweenness[order(metabolic_betweenness, decreasing = T)][1:400]))
high_centrality_to_remove = c("h2o","h2o2","h","o2","oh1","hco3","atp","adp","amp","pi","ppi",
                              "nad","nadh","nadp","nadph","fadh2","fad",
                              "co2","nh4","cl","so4","so3","o2s",
                              "na1","zn2","mg2","cu2","ca2","fe2","fe3","k")
all_vertex = V(metabolic_graph)$name
all_vertex_no_compartment = gsub("\\[", "", gsub("[\\[a-z]]", "", all_vertex))
vertex_to_remove = all_vertex[all_vertex_no_compartment %in% high_centrality_to_remove]
metabolic_graph <- delete_vertices(metabolic_graph, vertex_to_remove)

##### Step 3: deploy pathfinding algorithm to find the shortest path #####
dist_matrix = matrix(nrow = length(metabolic_graph), ncol = length(metabolic_graph), 
                     dimnames = list(row_names = V(metabolic_graph)$name, col_names = V(metabolic_graph)$name))

for (i in 1:ncol(dist_matrix)){
  road_length = lapply(1:length(metabolic_graph), function(x){
    length(shortest_paths(metabolic_graph, colnames(dist_matrix)[i], rownames(dist_matrix)[x])$vpath[[1]])-1
  })
  dist_matrix[,i] = unlist(road_length)
  print(i)
}

# 0 means the distance between the same vertice, -1 means not reachable between designated vertices,
# other value denotes the length of shortest path

fwrite(dist_matrix, "03 Results/04 Adjacency/Metabolie Adjacency Matrix Recon3D.csv")
```
