## Supplementary material for "Predicting substrates for orphan Solute Carrier Proteins using multi-omics datasets": Code for generating results: proximal and distant (rho).R

```
##### 0 load libraries #####
library(data.table)
library(tidyverse)

##### 1 read files #####
AM = fread("03 Results/04 Adjacency/Metabolie Adjacency Matrix Recon3D.csv")
tb = fread("01 Raw Data/00 SLC annotation 2024/book construction/Translation book (3 Matrix).csv")
et = fread("03 Results/03 Table alignment/Evidence table (rho).csv")
ki = fread("01 Raw Data/00 SLC annotation 2024/known interaction/Known interaction formula 20240321.csv")[1:837]

##### 2.1 Initialise Known interaction for Evidence table (etki) #####
##### 2 assemble etki #####
# remove substrates not covered by evidence table formula and derivatives
ki = ki[!is.na(Formula)][Status != "Derivative"]

etki = ki
colnames(etki)[1] = "SYMBOL"
etki$NCI60 = vector("numeric", nrow(etki))
etki$CCLE2019 = vector("numeric", nrow(etki))
etki$CCL180 = vector("numeric", nrow(etki))

for (i in 1:nrow(etki)){
  if (is.na(etki$Formula[i])){
    etki[i, c("NCI60", "CCLE2019", "CCL180") := list(NA, NA, NA)]
  }else if (etki$SYMBOL[i] %in% unique(et$SYMBOL)){
    etki[i, c("NCI60", "CCLE2019", "CCL180") :=
           as.list(et[SYMBOL == etki$SYMBOL[i] & Formula == etki$Formula[i]][,NCI60:CCL180])]
  }else{
    etki[i, c("NCI60", "CCLE2019", "CCL180") := list(NA, NA, NA)]
  }
}

##### 2.2 Initialise adjacency matrix #####
extracellular = grep("\\[e\\]", colnames(AM))
AM = AM[-extracellular, -extracellular, with = F]
AM = cbind(Compound = colnames(AM), AM)
AM_compartment = data.table(Compound = gsub("\\[.*?\\]", "", AM$Compound),
                            Compartment = gsub(".*(\\[.*\\]).*", "\\1", AM$Compound))

# ConvertNames #
# this function interchanges information in the translation table #
ConvertNames = function(name, source){
  # Process source input #
  if (is.null(source)){
    stop("Source of metabolite name required!")
  }
  else if (source %in% colnames(tb)){
    # Process name input #
    if (is.null(name)){
      stop("Name of metabolite required!")
    }else if (name %in% tb[,get(source)]){
      ans = tb[get(source) == name]
    }else{
      stop("Name entered not recognised!")
    }
  }else{
    stop("Source entered not recognised!")
  }
  
  # return ans as result #
  return(ans)
}

# getAM #
# this function finds the derivative compound of input formula, return a named list of kegg id, with the name indicating the steps
# required to reach the vertices
getAM = function(formula){
  recon_id = as.character(ConvertNames(formula, "Formula")[,"Recon3D"])
  qc = TRUE # logical term of quality control
  if(recon_id == ""){ # if there is no corresponding recon id annotated
    qc = FALSE
  }else{ # if there is/are corresponding recon id annotated
    recon_id = unlist(str_split(recon_id, "; "))
    recon_id_qc = recon_id[recon_id %in% AM_compartment$Compound]
    if (rlang::is_empty(recon_id_qc)){
      qc = FALSE
    }
  }
  
  # if compound passed qc and there's no requirement of finding infinite metabolites #
  if(qc){ # if compound pass qc #
    recon_id_zoom_in = c()
    # gather adjacent metabolite for the given recon id #
    for (i in 1:length(recon_id_qc)){
      AM_format_id_table = AM_compartment[Compound == recon_id_qc[i]]
      # get metabolites in different compartments, and remove
      AM_format_id = unlist(lapply(1:nrow(AM_format_id_table), function(x){paste(AM_format_id_table$Compound[x],
                                                                           AM_format_id_table$Compartment[x],
                                                                           sep = "")}))
      recon_id_zoom_in = c(recon_id_zoom_in, AM_format_id)
    }
    AM_related = AM[Compound %in% AM_format_id]
    return(AM_related)
  }else{ # if compound did not pass qc #
    return(NA)
  }
}

# extract.reason #
# this function is applied when step is 1, in order to discern transport activity from a step of conversion
extract.reason = function(ans){
  ans[, reason := vector("character", nrow(ans))]
  for (i in 1:nrow(ans)){
    substrate_chem = gsub("\\[.*?\\]", "", ans[i, substrate])
    substrate_com = gsub(".*(\\[.*\\]).*", "\\1", ans[i, substrate])
    derivative_chem = gsub("\\[.*?\\]", "", ans[i, derivative])
    derivative_com = gsub(".*(\\[.*\\]).*", "\\1",  ans[i, derivative])
    if (substrate_chem == derivative_chem & substrate_com != derivative_com){
      ans[i, reason := "Transport"]
    }else{
      ans[i, reason := paste("Conversion in", substrate_com)]
    }
  }
  return(ans)
}

# locate.proximity #
# this function returns a list containing information about the step of conversion from one node (substrate) to another (derivative)
locate.proximity = function(formula, step){
  proximal = getAM(formula)
  if (is.null(nrow(proximal))){
    return(NA)
  }else{
    mat = as.matrix(proximal)
    mat[, 2:ncol(mat)] = apply(mat[, 2:ncol(mat), drop = FALSE], 2, function(x) as.numeric(trimws(x)))
    coords = which(mat == step, arr.ind = TRUE)
    if (nrow(coords) == 0){
      return(NA)
    }else{
      ans = data.table(substrate = unlist(lapply(1:nrow(coords), function(x){proximal[,1][coords[x,1]]})),
                       derivative = unlist(lapply(1:nrow(coords), function(x){colnames(proximal)[coords[x,2]]})),
                       step = step)
      if (step == 1){
        ans = extract.reason(ans)
        ans = ans[reason != "Transport"]
        if (nrow(ans) == 0){
          return(NA)
        }
      }
      return(ans)
    }
  }
}

##### 3 calculating WT.pvalue for etki_qc across steps #####
# translate() capture and translate the recon id from the output of locate.proximity()
# How to use: Recon.id = locate.proximity("C10H16N5O13P3 (ATP)", 1)[, derivative]
translate = function(Recon.id){
  Recon.id = unique(gsub("\\[.*?\\]", "", Recon.id))
  Recon_info = data.table(Recon = as.character(Recon.id),
                          Formula = vector("character", length(Recon.id)))
  for (j in 1:nrow(Recon_info)){
    info = grep(Recon_info$Recon[j], tb$Recon3D)
    if (length(info) == 0){
      Recon_info[j, Formula := NA]
    }else{
      check_info = tb$Recon3D[info]
      true_info = c()
      for (k in 1:length(info)){
        if (Recon_info$Recon[j] %in% unlist(str_split(check_info[k], "; "))){
          true_info = c(true_info, info[k])
        }
      }
      if (length(true_info) != 0){
        Recon_info[j, Formula := paste(tb$Formula[true_info], collapse = "<>")]
      }else{
        Recon_info[j, Formula := NA]
      }
    }
  }
  # Remove the derivative that's not found in the translation book #
  Recon_info = na.omit(Recon_info)
  # if no derivative is found in the translation book, stop translation #
  if (nrow(Recon_info) == 0){
    return(NA)
  }else if (length(grep("<>", Recon_info$Formula)) == 0){
    return(Recon_info)
  }else{
    # Break the <> and insert a new row #
    Recon_info = Recon_info %>% 
      mutate(Formula = strsplit(as.character(Formula), "<>")) %>% 
      unnest(Formula)
    Recon_info = as.data.table(Recon_info)
    return(Recon_info)
  }
}

# getNR() gathers the information for formula after the formula is translated by translate()
getNR = function(gene, translated.formula){
  data = data.table(Formula = translated.formula[, Formula],
                    NCI60 = vector("numeric", nrow(translated.formula)),
                    CCLE2019 = vector("numeric", nrow(translated.formula)),
                    CCL180 = vector("numeric", nrow(translated.formula)))
  for (k in 1:nrow(data)){
    if(nrow(et[SYMBOL == gene & Formula == data[k, Formula]]) != 0){
      data[k, c("NCI60", "CCLE2019", "CCL180") :=
             as.list(et[SYMBOL == gene & Formula == data[k, Formula]][,NCI60:CCL180])]
    }else{
      data[k, c("NCI60", "CCLE2019", "CCL180") := as.list(NA, NA, NA)]
    }
  }
  return(data)
}

# getSource() acquires the metabolomics which contains the concentration info for this formula #
getSource = function(formula){
  logical = as.logical((is.na(tb[Formula == formula, NCI60:CCL180])))
  ans = colnames(tb[,NCI60:CCL180])[!logical]
  return(ans)
}

# channel calculates the PS difference between compounds and derivatives #
channel = function(derivative_table, substrate_table, s_source){
  # collecting rho and p value #
  rho_diff_val = c()
  for (m in 1:nrow(derivative_table)){
    d_source = getSource(derivative_table$Formula[m]) # the source of derivatives
    channeling = length(s_source)*length(d_source) # calculate channeling value
    ### 1 substrate source and 1 derivative source ###
    if (channeling == 1){
      rho_diff_val = c(rho_diff_val, as.numeric(substrate_table[, s_source, with = FALSE]) - 
                         as.numeric(derivative_table[m, d_source, with = FALSE])
      )
    }
    ### 1 comp source and 2 deri source | 2 comp source and 1 deri source ###
    else if(channeling == 2 | channeling == 3){
      same_source = intersect(s_source, d_source)
      ## if there is one omic source overlapped ##
      if (length(same_source) != 0){
        rho_diff_val = c(rho_diff_val, as.numeric(substrate_table[, same_source, with = FALSE]) - 
                           as.numeric(derivative_table[m, same_source, with = FALSE])
        )
        ## if there is no omic source overlapped, calculate the average of correlation statistics ##
      }else{
        diff = c()
        if (length(s_source) > length(d_source)){
          for (n in 1:length(s_source)){
            diff = c(diff, as.numeric(substrate_table[, s_source[n], with = FALSE]) -
                       as.numeric(derivative_table[m, d_source, with = FALSE]))
          }
          rho_diff_val = c(rho_diff_val, mean(diff, na.rm = TRUE))
        }else{
          for (n in 1:length(d_source)){
            diff = c(diff, as.numeric(substrate_table[, s_source, with = FALSE]) -
                       as.numeric(derivative_table[m, d_source[n], with = FALSE]))
          }
          rho_diff_val = c(rho_diff_val, mean(diff, na.rm = TRUE))
        }
      }
    }
    ### 2 comp source and 2 deri source, 3 comp source and 3 deri source ###
    else if (channeling == 4 | channeling == 9){
      same_source = intersect(s_source, d_source)
      ## if there is 1 omics source shared ##
      if (length(same_source) == 1){
        rho_diff_val = c(rho_diff_val, as.numeric(substrate_table[, same_source, with = FALSE]) - 
                           as.numeric(derivative_table[m, same_source, with = FALSE])
        )
        ## if there are 2 or 3 omics source shared, calculate the average of correlation statistics ##
      }else if (length(same_source) > 1){
        diff = c()
        for (n in 1:length(same_source)){
          diff = c(diff, as.numeric(substrate_table[, same_source[n], with = FALSE]) - 
                     as.numeric(derivative_table[m, same_source[n], with = FALSE]))
        }
        rho_diff_val = c(rho_diff_val, mean(diff, na.rm = TRUE))
      }
    }
    ### 2 comp source and 3 deri source, 3 comp source and 2 deri source ###
    else if (channeling == 6){
      ## however change, there will be 2 overlapping omics ##
      same_source = intersect(s_source, d_source)
      diff = c()
      for (n in 1:length(same_source)){
        diff = c(diff, as.numeric(substrate_table[, same_source[n], with = FALSE]) - 
                   as.numeric(derivative_table[m, same_source[n], with = FALSE]))
      }
      rho_diff_val = c(rho_diff_val, mean(diff, na.rm = TRUE))
    }
  }
  return(rho_diff_val)
}

# check_NA checks whether the output is NA
check_NA = function(x){
  if (length(x) == 1){
    if (is.na(x)){
      return(TRUE)
    }else{
      return(FALSE)
    }
  }else{
    return(FALSE)
  }
}

step_table = data.table(Steps = seq(1, 10, 1),
                        WT.pvalue = vector("numeric", length(seq(1, 10, 1))))
Inf_across_step = vector("list", nrow(step_table))
for (i in 1:nrow(step_table)){
  step = step_table[i, Steps]
  # make dNR_step #
  dNR_step = data.table(SYMBOL = etki[,SYMBOL],
                        Formula = etki[,Formula],
                        dNR = vector("numeric", nrow(etki))
  )
  for (j in 1:nrow(etki)){
    proximal = locate.proximity(etki[j, Formula], step)
    if (is.null(nrow(proximal))){               # if there is no proximal derivative at this step
      dNR_step[j, dNR := NaN]               # there won't be a NR difference to compare for this compound.
    }else{                                  # if there is/are proximal derivative(s) at this step
      proximal_temp = translate(proximal[, derivative])   # translate the proximal derivative(s) to formula name(s).
      if (check_NA(proximal_temp)){
        dNR_step[j, dNR := NaN]
      }else{
        proximal_temp = getNR(etki[j, SYMBOL], proximal_temp)
        proximal_dNR = channel(proximal_temp, etki[j], getSource(etki[j, Formula]))
        dNR_step[j, dNR := mean(abs(proximal_dNR), na.rm = TRUE)]
      }
    }
    print(paste("step", i ,":", j))
  }
  dNR_step = na.omit(dNR_step)
  
  # take the dNR_step and make dNR_inf #
  dNR_inf = data.table(SYMBOL = dNR_step[,SYMBOL],
                       Formula = dNR_step[,Formula],
                       dNR = vector("numeric", nrow(dNR_step))
  )
  for (j in 1:nrow(dNR_inf)){
    infinite = locate.proximity(dNR_step[j,Formula], -1)
    if (is.null(nrow(infinite))){             # if there is no infinite derivative at this step
      dNR_inf[j, dNR := NaN]                # there won't be a NR difference to compare for this compound.
    }else{                                  # if there is/are infinite derivative(s) at this step
      infinite_temp = translate(infinite[, derivative])   # translate the infinite derivative(s) to formula name(s).
      if (check_NA(infinite_temp)){      # if the infinite only has one length
        dNR_inf[j, dNR := NaN]
      }else{
        infinite_temp = infinite_temp[sample(1:nrow(infinite_temp), 20)]
        infinite_temp = getNR(dNR_inf[j, SYMBOL], infinite_temp)
        etki_reference = etki[SYMBOL == dNR_inf[j, SYMBOL] & Formula == dNR_inf[j, Formula]]
        if(nrow(etki_reference) > 1){
          etki_reference = etki_reference[1]
        }
        infinite_dNR = channel(infinite_temp, etki_reference, getSource(dNR_inf[j, Formula]))
        dNR_inf[j, dNR := mean(abs(infinite_dNR), na.rm = TRUE)]
      }
    }
    print(paste("step inf:", j))
  }
  dNR_inf = na.omit(dNR_inf)
  WT = wilcox.test(dNR_step$dNR, dNR_inf$dNR, alternative = "less")$p.value
  # store data #
  Inf_across_step[[i]] = dNR_inf
  step_table[i, WT.pvalue := WT]
  print(paste("step", i, "completed"))
}

##### 5 Benchmarking #####
step_table_benchmarking = data.table(Steps = rep(seq(1,10,1), each = 100),
                                     WT.pvalue = vector("numeric", length(rep(seq(1,10,1), each = 100)))
)

for (i in 1:nrow(step_table_benchmarking)){
  step = step_table_benchmarking[i, Steps]
  # make dNR_step #
  dNR_step = data.table(SYMBOL = etki[,SYMBOL],
                        Formula = etki[,Formula],
                        dNR = vector("numeric", nrow(etki))
  )
  for (j in 1:nrow(etki)){
    proximal = locate.proximity(etki[j, Formula], step)
    if (is.null(nrow(proximal))){             # if there is no proximal derivative at this step
      dNR_step[j, dNR := NaN]               # there won't be a PS difference to compare for this compound.
    }else{                                  # if there is/are proximal derivative(s) at this step
      proximal_temp = translate(proximal[, derivative])   # translate the proximal derivative(s) to formula name(s).
      if (check_NA(proximal_temp)){      # if the proximal only has one length,
        dNR_step[j, dNR := NaN]
      }else{
        temp_fake = locate.proximity(etki[j, Formula], -1)
        temp_fake = translate(temp_fake[, derivative])
        temp_fake = temp_fake[sample(1:nrow(temp_fake), length(proximal_temp))]
        temp_fake = getNR(etki[j, SYMBOL], temp_fake)
        fake_dNR = channel(temp_fake, etki[j], getSource(etki[j, Formula]))
        dNR_step[j, dNR := mean(abs(fake_dNR), na.rm = TRUE)]
      }
    }
    print(paste("row", i ,":", j))
  }
  dNR_step = na.omit(dNR_step)
  
  # compare with stored dNR_inf #
  dNR_inf = Inf_across_step[[(step)]]
  WT = wilcox.test(dNR_step$dNR, dNR_inf$dNR, alternative = "less")$p.value
  # store data #
  step_table_benchmarking[i, WT.pvalue := WT]
  print(paste("row", i, "completed"))
}

# adjust p-value for every step due to multiple comparison #
step_table_benchmarking[, class := rep("simulated", nrow(step_table_benchmarking))]
step_table[, class := rep("known", nrow(step_table))]

step_assembly = rbind(step_table_benchmarking, step_table)

for (step in seq(1, 10, 1)){
  p = step_assembly[Steps == step, WT.pvalue]
  step_assembly[Steps == step, WT.pvalue := p.adjust(p, method = "BH", n = length(p))]
  print(step)
}

fwrite(step_assembly, "03 Results/04 Adjacency/proximal and distant (rho).csv")

step_simulated = step_assembly[class == "simulated"]
step_known = step_assembly[class == "known"]

g = ggplot() +
  # box plot of fecs
  geom_boxplot(data = step_simulated, 
               aes(x = Steps, y= -log10(WT.pvalue), group = Steps)) +
  # points of ecs
  geom_line(data = step_known,
            aes(x= Steps, y= -log10(WT.pvalue)),
            color = "#990000",
            linetype="dashed") + 
  theme(
    legend.position="none",
    plot.title = element_text(size=11)
  ) +
  theme(panel.grid.major = element_blank(), panel.grid.minor = element_blank(),
        panel.background = element_blank(), axis.line = element_line(colour = "black")) +
  scale_x_continuous(breaks=seq(1,11,2))
g
```
