## Supplementary material for "Predicting substrates for orphan Solute Carrier Proteins using multi-omics datasets": Code for generating results: Step0_Control Generation.R

```
library(data.table)
library(tidyverse)

##### 1 read file #####
et <- fread("03 Results/03 Table alignment/Evidence table (normalised rho).csv")
ki <- fread("01 Raw Data/00 SLC annotation 2024/known interaction/Known interaction formula 20240321.csv")[1:841]
tb <- fread("01 Raw Data/00 SLC annotation 2024/book construction/Translation book (3 Matrix).csv")

##### 2 assemble etki #####
# remove substrates not covered by evidence table formula and derivatives
ki <- ki[!is.na(Formula)][Status != "Derivative"]

etki = ki
colnames(etki)[1] = "SYMBOL"
etki$NCI60 = vector("numeric", nrow(etki))
etki$CCLE2019 = vector("numeric", nrow(etki))
etki$CCL180 = vector("numeric", nrow(etki))
etki$CRISPRDEP = vector("numeric", nrow(etki))

for (i in 1:nrow(etki)){
  if (etki$SYMBOL[i] %in% unique(et$SYMBOL)){
    etki[i, c("NCI60", "CCLE2019", "CCL180", "CRISPRDEP") :=
           as.list(et[SYMBOL == etki$SYMBOL[i] & Formula == etki$Formula[i]][,NCI60:CRISPRDEP])]
  }else{
    etki[i, c("NCI60", "CCLE2019", "CCL180", "CRISPRDEP") := list(NA, NA, NA, NA)]
  }
}
etki = etki[rowSums(is.na(etki[,8:11]))<4]

##### 3 Generate fake etki #####
fake_etki_collection = vector("list", 100)
etki_gene_pool = unique(etki$SYMBOL)
etki_metabolite_pool = unique(etki$Formula)

for (sample_time in 1:100){
  # initialise fake etki data table #
  fake_etki = setNames(data.table(matrix(ncol = ncol(etki), nrow = nrow(etki))), colnames(etki))
  fake_etki[, Substrate := NULL]
  char.col = colnames(fake_etki)[1:6]
  num.col = colnames(fake_etki)[7:10]
  fake_etki[, (char.col) := lapply(.SD, as.character), .SDcols = char.col]
  fake_etki[, (num.col) := lapply(.SD, as.character), .SDcols = num.col]
  fake_etki[, c("SYMBOL", "Gene_type","Localisation") := as.list(etki[, c("SYMBOL", "Gene_type","Localisation")])]
  fake_etki[, Status := rep("Fake", nrow(fake_etki))]
  # for each SLC, shuffle fake substrate formula #
  for (j in 1:nrow(fake_etki)){
    true_substrate = etki[SYMBOL == fake_etki$SYMBOL[j]][, Formula]
    fake_substrate_pool = etki_metabolite_pool[!etki_metabolite_pool %in% true_substrate]
    fake_substrate = sample(fake_substrate_pool, 1)
    while (nrow(et[SYMBOL == fake_etki$SYMBOL[j] & Formula == fake_substrate][,NCI60:CRISPRDEP]) == 0){
      fake_substrate = sample(fake_substrate_pool, 1)
    }
    fake_etki[j, Formula := fake_substrate]
    fake_etki[j, c("NCI60", "CCLE2019", "CCL180", "CRISPRDEP") := 
                as.list(et[SYMBOL == fake_etki$SYMBOL[j] & Formula == fake_etki$Formula[j]][,NCI60:CRISPRDEP])]
    print(paste(sample_time, ":", round(j/nrow(fake_etki),2)))
  }
  fake_etki_collection[[sample_time]] = fake_etki
  print(paste(sample_time, "done"))
}
```
