## Supplementary material for "Predicting substrates for orphan Solute Carrier Proteins using multi-omics datasets": Code for generating results: Step1_Building Factor A (HPC).R

```
##### 0 load library and random data #####
library(data.table)
load("~/Desktop/SLC ver3/03 Results/05 Building ConfidenceScore/Building ConfidenceScore (clean start with generated random).RData")

##### 1 Respective predictive score distribution for each data #####
numbering_range = seq(-2, 2, 0.1)

ConfidenceScore_A = function(ki.data, a, b, c){
  ki_score = setNames(data.table(matrix(ncol = 6, nrow = nrow(ki.data))), c("HGNC_symbol","Localisation","Substrate","Formula","Status","Score"))
  char.col = colnames(ki_score)[1:5]
  num.col = colnames(ki_score)[6]
  ki_score[, (char.col) := lapply(.SD, as.character), .SDcols = char.col]
  ki_score[, (num.col) := lapply(.SD, as.character), .SDcols = num.col]
  ki_score[, 1:5 := as.list(ki.data[, 1:5])]
  
  # Factor A: Predictive Scores
  ## quantile ##
  ns_dist1 = na.omit(etki$NCI60) # NCI60 #
  ns_dist2 = na.omit(etki$CCLE2019) # CCLE2019 #
  ns_dist3 = na.omit(etki$CCL180) # CCL180 #
  q1 = quantile(ns_dist1[ns_dist1 > as.numeric(a)], seq(0, 1, by = 0.1))
  q2 = quantile(ns_dist2[ns_dist2 > as.numeric(b)], seq(0, 1, by = 0.1))
  q3 = quantile(ns_dist3[ns_dist3 > as.numeric(c)], seq(0, 1, by = 0.1))
  
  Factor_A_info = data.table(Factor_A_name = c("NCI60", "CCLE2019", "CCL180"),
                             Factor_A_dist = c("q1", "q2", "q3"),
                             Factor_A_threshold = c(as.numeric(a), as.numeric(b), as.numeric(c)))
  for (i in 1:nrow(ki.data)){
    # score prediction for 3 matrices #
    Factor_A = vector("integer", nrow(Factor_A_info))
    for (j in 1:nrow(Factor_A_info)){
      n = as.numeric(ki.data[i, get(Factor_A_info[j, Factor_A_name])])
      if (is.na(n) | n < Factor_A_info[j, Factor_A_threshold]){
        Factor_A[j] = 0
      } else if (n > Factor_A_info[j, Factor_A_threshold]){
        Factor_A[j] = findInterval(n, get(Factor_A_info[j, Factor_A_dist]))*3
      }
    }
    Factor_A = sum(Factor_A)
    # calculate final score #
    ki_score[i, Score := Factor_A]
  }
  return(ki_score)
}
Per_collection = vector("list", length = length(numbering_range)^3)
for (a in numbering_range){
  for (b in numbering_range){
    for (c in numbering_range){
      ecs = ConfidenceScore_A(etki,a,b,c)
      fecs = lapply(fake_etki_collection, function(x){ConfidenceScore_A(x,a,b,c)})
      Per = data.table(A = a,
                       B = b,
                       C = c,
                       X = 1:40,
                       Y = rep(0, 40),
                       Z = rep(0, 40),
                       Q = rep(0, 40))
      for (n in 1:40){
        y = ecs[as.numeric(Score) > n, .N]/ecs[, .N]
        z = mean(unlist(lapply(fecs, function(f){f[as.numeric(Score) > n, .N]})))/ecs[, .N]
        Per[n, Y := y]
        Per[n, Z := z]
        Per[n, Q := y-z]
      }
      index = which(numbering_range == a)*which(numbering_range == b)*which(numbering_range == c)
      Per_collection[[index]] = Per
      print(index)
    }
  }
  save_path = paste("~/BuildA/a_",a, "collection.RData")
}
```
