## Supplementary material for "Predicting substrates for orphan Solute Carrier Proteins using multi-omics datasets": Code for generating results: Step1_Building Factor A.R

```
##### 0 load library and random data #####
library(data.table)
library(tidyverse)
load("~/Desktop/SLC ver3/03 Results/05 Building ConfidenceScore/Building ConfidenceScore (clean start with generated random).RData")

##### 1 Respective predictive score distribution for each data #####
numbering_range = seq(0,1,0.1)

ConfidenceScore_A = function(ki.data, a, b, c){
  ki_score = setNames(data.table(matrix(ncol = 6, nrow = nrow(ki.data))), c("HGNC_symbol","Localisation","Substrate","Formula","Status","Score"))
  char.col = colnames(ki_score)[1:5]
  num.col = colnames(ki_score)[6]
  ki_score[, (char.col) := lapply(.SD, as.character), .SDcols = char.col]
  ki_score[, (num.col) := lapply(.SD, as.character), .SDcols = num.col]
  ki_score[, 1:5 := as.list(ki.data[, 1:5])]
  
  # Factor A: Predictive Scores
  ## quantile ##
  ns_dist1 = na.omit(etki$NCI60) # NCI60 #
  ns_dist2 = na.omit(etki$CCLE2019) # CCLE2019 #
  ns_dist3 = na.omit(etki$CCL180) # CCL180 #
  q1 = quantile(ns_dist1[ns_dist1 > as.numeric(a)], seq(0, 1, by = 0.1))
  q2 = quantile(ns_dist2[ns_dist2 > as.numeric(b)], seq(0, 1, by = 0.1))
  q3 = quantile(ns_dist3[ns_dist3 > as.numeric(c)], seq(0, 1, by = 0.1))
  
  Factor_A_info = data.table(Factor_A_name = c("NCI60", "CCLE2019", "CCL180"),
                             Factor_A_dist = c("q1", "q2", "q3"),
                             Factor_A_threshold = c(as.numeric(a), as.numeric(b), as.numeric(c)))
  for (i in 1:nrow(ki.data)){
    # score prediction for 3 matrices #
    Factor_A = vector("integer", nrow(Factor_A_info))
    for (j in 1:nrow(Factor_A_info)){
      n = as.numeric(ki.data[i, get(Factor_A_info[j, Factor_A_name])])
      if (is.na(n) | n < Factor_A_info[j, Factor_A_threshold]){
        Factor_A[j] = 0
      } else if (n > Factor_A_info[j, Factor_A_threshold]){
        Factor_A[j] = findInterval(n, get(Factor_A_info[j, Factor_A_dist]))*3
      }
    }
    Factor_A = sum(Factor_A)
    # calculate final score #
    ki_score[i, Score := Factor_A]
  }
  return(ki_score)
}
##### 2 ConfidenceScore_Ax: changing NCI60 threshold (q1) #####
## Calculate Confidence Score for known and random interactions, changing CCLE2019 threshold ##
ecs_collection_Ax = vector(mode = "list", length = length(numbering_range))
names(ecs_collection_Ax) = numbering_range
for (i in 1:length(ecs_collection_Ax)){
  ecs_collection_Ax[[i]] = ConfidenceScore_A(etki, 
                                             a = names(ecs_collection_Ax)[i], 
                                             b = Inf,
                                             c = Inf)
  print(i)
}
fecs_collection_Ax = vector(mode = "list", length = length(numbering_range))
names(fecs_collection_Ax) = numbering_range
for (i in 1:length(fecs_collection_Ax)){
  fecs_collection_Ax[[i]] = lapply(fake_etki_collection, function(x){ConfidenceScore_A(x, 
                                                                                       a = names(ecs_collection_Ax)[i], 
                                                                                       b = Inf,
                                                                                       c = Inf)})
  print(i)
}

## Collect statistical reporting in known interaction across CCLE2019 threshold ##
ecs_across_threshold = data.table(Threshold = numbering_range, 
                                  Status = rep("true", length(numbering_range)),
                                  Mean = vector("numeric", length(numbering_range)))
for (i in 1:length(ecs_collection_Ax)){
  ecs = ecs_collection_Ax[[i]]
  ecs_across_threshold[i, Mean := mean(as.numeric(ecs[,Score]))]
  print(i)
}

## Collect statistical reporting in random interaction across CCLE2019 threshold ##
fecs_across_threshold = data.table(Threshold = rep(numbering_range, each = 100), 
                                   Status = rep("random", length(rep(numbering_range, each = 100))),
                                   Mean = vector("numeric", length(rep(numbering_range, each = 100))))
for (i in 1:length(fecs_collection_Ax)){
  fecs_confidence_score = fecs_collection_Ax[[i]]
  fecs_confidence_score_mean = unlist(lapply(fecs_confidence_score, function(x){mean(as.numeric(x[,Score]))}))
  fecs_across_threshold[Threshold == names(fecs_collection_Ax)[i], Mean := fecs_confidence_score_mean]
  print(i)
}

## construct table for Boxplot figure ##
across_threshold_x = rbind(ecs_across_threshold, fecs_across_threshold)
fwrite(across_threshold_x, "03 Results/05 Building ConfidenceScore/Factor A/across threshold_factorAx.csv")

# Across all threshold selected for CCLE2019, known interaction behave better than random #
ggplot() +
  # box plot of fecs
  geom_boxplot(data = fecs_across_threshold, 
               aes(x = Threshold, y= Mean, group = Threshold)) +
  # points of ecs
  geom_point(data = ecs_across_threshold,
             aes(x= Threshold, y= Mean),
             color = 'red')

## Generate SWEEP table ##
# Th: Predictive score cutoff for the interaction to be marked as "discovered"
# X: cutoff for real score
# Y: percentage of real interactions > X
# Z: percentage of fake interactions > X

Per0 = data.table(Th = NULL, 
                  X = NULL,
                  Y = NULL,
                  Z = NULL,
                  Q = NULL)

for (i in 1:length(numbering_range)){
  ecs = ecs_collection_Ax[[i]]
  fecs = fecs_collection_Ax[[i]]
  Per = data.table(Th = rep(names(ecs_collection_Ax)[i], 40), 
                   X = 1:40,
                   Y = rep(0, 40),
                   Z = rep(0, 40),
                   Q = rep(0, 40))
  for (x in 1:40){
    y = ecs[as.numeric(Score) > x, .N]/ecs[, .N]
    z = mean(unlist(lapply(fecs, function(a){a[as.numeric(Score) > x, .N]})))/ecs[, .N]
    Per[x, Y := y]
    Per[x, Z := z]
    Per[x, Q := y-z]
  }
  Per0 = rbind(Per0, Per)
  print(i)
}

ConfAx = Per0
fwrite(ConfAx, "03 Results/05 Building ConfidenceScore/Factor A/Sweep_factorAx.csv")


## Sweep for maximum fraction difference ##

Sweep = c()
for(i in unique(ConfAx$Th)){
  temp = ConfAx[Th == i] # temporary data table for given threshold
  #plot(temp$X, temp$Q, ylab="Fraction difference",xlab="Confidence cutoff", pch=16, main=paste("True positive vs cutoff, Th=",i, sep=""))
  MaxQ = mean(which(temp$Q==max(temp$Q)))
  MaxX = mean(temp$X[MaxQ])
  Sweep = rbind(Sweep, c(MaxX,temp$Q[MaxQ]))
}

Sweep = cbind(numbering_range,Sweep)
colnames(Sweep) = c("Th","Xmax","MaxTP")

plot(Sweep[,1],Sweep[,3], pch=18, col = "#69b3a2", xaxt="n")
axis(1, at = numbering_range)

## RESULT: use -1.2 for NCI60 threshold ##

#######################################################################################################################

##### 3 ConfidenceScore_Ay: changing CCLE2019 threshold (q2) #####
## Calculate Confidence Score for known and random interactions, changing CCLE2019 threshold ##
ecs_collection_Ay = vector(mode = "list", length = length(numbering_range))
names(ecs_collection_Ay) = numbering_range
for (i in 1:length(ecs_collection_Ay)){
  ecs_collection_Ay[[i]] = ConfidenceScore_A(etki, 
                                             a = 0, 
                                             b = names(ecs_collection_Ax)[i],
                                             c = Inf)
  print(i)
}
fecs_collection_Ay = vector(mode = "list", length = length(numbering_range))
names(fecs_collection_Ay) = numbering_range
for (i in 1:length(fecs_collection_Ay)){
  fecs_collection_Ay[[i]] = lapply(fake_etki_collection, function(x){ConfidenceScore_A(x, 
                                                                                       a = 0, 
                                                                                       b = names(ecs_collection_Ax)[i],
                                                                                       c = Inf)})
  print(i)
}

## Collect statistical reporting in known interaction across NCI60 threshold ##
ecs_across_threshold = data.table(Threshold = numbering_range, 
                                  Status = rep("true", length(numbering_range)),
                                  Mean = vector("numeric", length(numbering_range)))
for (i in 1:length(ecs_collection_Ay)){
  ecs = ecs_collection_Ay[[i]]
  ecs_across_threshold[i, Mean := mean(as.numeric(ecs[,Score]))]
  print(i)
}

# Collect statistical reporting in random interaction across NCI60 threshold #
fecs_across_threshold = data.table(Threshold = rep(numbering_range, each = 100), 
                                   Status = rep("random", length(rep(numbering_range, each = 100))),
                                   Mean = vector("numeric", length(rep(numbering_range, each = 100))))
for (i in 1:length(fecs_collection_Ay)){
  fecs_confidence_score = fecs_collection_Ay[[i]]
  fecs_confidence_score_mean = unlist(lapply(fecs_confidence_score, function(x){mean(as.numeric(x[,Score]))}))
  fecs_across_threshold[Threshold == names(fecs_collection_Ay)[i], Mean := fecs_confidence_score_mean]
  print(i)
}

## construct table for figure ##
across_threshold_y = rbind(ecs_across_threshold, fecs_across_threshold)
fwrite(across_threshold_y, "03 Results/05 Building ConfidenceScore/Factor A/across threshold_factorAy.csv")

# Across all threshold selected for NCI60, known interaction behave better than random #
ggplot() +
  # box plot of fecs
  geom_boxplot(data = fecs_across_threshold, 
               aes(x = Threshold, y= Mean, group = Threshold)) +
  # points of ecs
  geom_point(data = ecs_across_threshold,
             aes(x= Threshold, y= Mean),
             color = 'red')

## Generate SWEEP table ##
# Th: Predictive score cutoff for the interaction to be marked as "discovered"
# X: cutoff for real score
# Y: percentage of real interactions > X
# Z: percentage of fake interactions > X

Per0 = data.table(Th = NULL, 
                  X = NULL,
                  Y = NULL,
                  Z = NULL,
                  Q = NULL)

for (i in 1:length(numbering_range)){
  ecs = ecs_collection_Ay[[i]]
  fecs = fecs_collection_Ay[[i]]
  Per = data.table(Th = rep(names(ecs_collection_Ay)[i], 40), 
                   X = 1:40,
                   Y = rep(0, 40),
                   Z = rep(0, 40),
                   Q = rep(0, 40))
  for (x in 1:40){
    y = ecs[as.numeric(Score) > x, .N]/ecs[, .N]
    z = mean(unlist(lapply(fecs, function(a){a[as.numeric(Score) > x, .N]})))/ecs[, .N]
    Per[x, Y := y]
    Per[x, Z := z]
    Per[x, Q := y-z]
  }
  Per0 = rbind(Per0, Per)
  print(i)
}

ConfAy = Per0
fwrite(ConfAy, "03 Results/05 Building ConfidenceScore/Factor A/Sweep_factorAy.csv")

## Sweep for maximum fraction difference ##
Sweep = c()

for(i in unique(ConfAy$Th)){
  temp = ConfAy[Th == i] # temporary data table for given threshold
  #plot(temp$X, temp$Q, ylab="Fraction difference",xlab="Confidence cutoff", pch=16, main=paste("True positive vs cutoff, Th=",i, sep=""))
  MaxQ = mean(which(temp$Q==max(temp$Q)))
  MaxX = mean(temp$X[MaxQ])
  Sweep = rbind(Sweep, c(MaxX,temp$Q[MaxQ]))
}

Sweep = cbind(numbering_range,Sweep)
colnames(Sweep) = c("Th","Xmax","MaxTP")

plot(Sweep[,1],Sweep[,3], pch=18, col = "#69b3a2", xaxt="n")
axis(1, at = numbering_range)

## RESULT: -1.4 for CCLE2019 threshold ##


#######################################################################################################################


##### 4 ConfidenceScore_Az: changing CCL180 threshold (q3) #####

## Calculate Confidence Score for known and random interactions, changing CCLE2019 threshold ##
ecs_collection_Az = vector(mode = "list", length = length(numbering_range))
names(ecs_collection_Az) = numbering_range
for (i in 1:length(ecs_collection_Az)){
  ecs_collection_Az[[i]] = ConfidenceScore_A(etki, 
                                             a = 0, 
                                             b = 0,
                                             c = names(ecs_collection_Az)[i])
  print(i)
}
fecs_collection_Az = vector(mode = "list", length = length(numbering_range))
names(fecs_collection_Az) = numbering_range
for (i in 1:length(fecs_collection_Az)){
  fecs_collection_Az[[i]] = lapply(fake_etki_collection, function(x){ConfidenceScore_A(x, 
                                                                                       a = 0, 
                                                                                       b = 0,
                                                                                       c = names(ecs_collection_Az)[i])})
  print(i)
}

## Collect statistical reporting in known interaction across CCL180 threshold ##
ecs_across_threshold = data.table(Threshold = numbering_range, 
                                  Status = rep("true", length(numbering_range)),
                                  Mean = vector("numeric", length(numbering_range)))
for (i in 1:length(ecs_collection_Az)){
  ecs = ecs_collection_Az[[i]]
  ecs_across_threshold[i, Mean := mean(as.numeric(ecs[,Score]))]
  print(i)
}

## Collect statistical reporting in random interaction across CCL180 threshold ##
fecs_across_threshold = data.table(Threshold = rep(numbering_range, each = 100), 
                                   Status = rep("random", length(rep(numbering_range, each = 100))),
                                   Mean = vector("numeric", length(rep(numbering_range, each = 100))))
for (i in 1:length(fecs_collection_Az)){
  fecs_confidence_score = fecs_collection_Az[[i]]
  fecs_confidence_score_mean = unlist(lapply(fecs_confidence_score, function(x){mean(as.numeric(x[,Score]))}))
  fecs_across_threshold[Threshold == names(fecs_collection_Az)[i], Mean := fecs_confidence_score_mean]
  print(i)
}

## construct table for figure ##
across_threshold_z = rbind(ecs_across_threshold, fecs_across_threshold)
fwrite(across_threshold_z, "03 Results/05 Building ConfidenceScore/Factor A/across threshold_factorAz.csv")

# Across all threshold selected for CCL180, known interaction behave better than random #
ggplot() +
  # box plot of fecs
  geom_boxplot(data = fecs_across_threshold, 
               aes(x = Threshold, y= Mean, group = Threshold)) +
  # points of ecs
  geom_point(data = ecs_across_threshold,
             aes(x= Threshold, y= Mean),
             color = 'red')

## Generate SWEEP table ##
# Th: Predictive score cutoff for the interaction to be marked as "discovered"
# X: cutoff for real score
# Y: percentage of real interactions > X
# Z: percentage of fake interactions > X

Per0 = data.table(Th = NULL, 
                  X = NULL,
                  Y = NULL,
                  Z = NULL,
                  Q = NULL)

for (i in 1:length(numbering_range)){
  ecs = ecs_collection_Az[[i]]
  fecs = fecs_collection_Az[[i]]
  Per = data.table(Th = rep(names(ecs_collection_Az)[i], 40), 
                   X = 1:40,
                   Y = rep(0, 40),
                   Z = rep(0, 40),
                   Q = rep(0, 40))
  for (x in 1:40){
    y = ecs[as.numeric(Score) > x, .N]/ecs[, .N]
    z = mean(unlist(lapply(fecs, function(a){a[as.numeric(Score) > x, .N]})))/ecs[, .N]
    Per[x, Y := y]
    Per[x, Z := z]
    Per[x, Q := y-z]
  }
  Per0 = rbind(Per0, Per)
  print(i)
}

ConfAz = Per0
fwrite(ConfAz, "03 Results/05 Building ConfidenceScore/Factor A/Sweep_factorAz.csv")

## Sweep ##
Sweep = c()

for(i in unique(ConfAz$Th)){
  temp = ConfAz[Th == i] # temporary data table for given threshold
  #plot(temp$X, temp$Q, ylab="Fraction difference",xlab="Confidence cutoff", pch=16, main=paste("True positive vs cutoff, Th=",i, sep=""))
  MaxQ = mean(which(temp$Q==max(temp$Q)))
  MaxX = mean(temp$X[MaxQ])
  Sweep = rbind(Sweep, c(MaxX,temp$Q[MaxQ]))
}

Sweep = cbind(numbering_range,Sweep)
colnames(Sweep) = c("Th","Xmax","MaxTP")

plot(Sweep[,1],Sweep[,3], pch=18, col = "#69b3a2", xaxt="n")
axis(1, at = numbering_range)

## RESULT: 1.9 for CCL180 threshold ##

##### 5 Discovery made from ConfidenceScore_A #####
# Calculating confidence score #
ConfidenceScore_A = function(ki.data){
  ki_score = setNames(data.table(matrix(ncol = 6, nrow = nrow(ki.data))), c("HGNC_symbol","Localisation","Substrate","Formula","Status","Score"))
  char.col = colnames(ki_score)[1:5]
  num.col = colnames(ki_score)[6]
  ki_score[, (char.col) := lapply(.SD, as.character), .SDcols = char.col]
  ki_score[, (num.col) := lapply(.SD, as.character), .SDcols = num.col]
  ki_score[, 1:5 := as.list(ki.data[, 1:5])]
  
  # Factor A: Predictive Scores
  ## quantile ##
  ns_dist1 = na.omit(etki$NCI60) # NCI60 #
  ns_dist2 = na.omit(etki$CCLE2019) # CCLE2019 #
  ns_dist3 = na.omit(etki$CCL180) # CCL180 #
  q1 = quantile(ns_dist1[ns_dist1 > 0], seq(0, 1, by = 0.1))
  q2 = quantile(ns_dist2[ns_dist2 > 0], seq(0, 1, by = 0.1))
  q3 = quantile(ns_dist3[ns_dist3 > 0.8], seq(0, 1, by = 0.1))

  Factor_A_info = data.table(Factor_A_name = c("NCI60", "CCLE2019", "CCL180"),
                             Factor_A_dist = c("q1", "q2", "q3"),
                             Factor_A_threshold = c(0, 0, 0.8))
  for (i in 1:nrow(ki.data)){
    # score prediction for 3 matrices #
    Factor_A = vector("integer", nrow(Factor_A_info))
    for (j in 1:nrow(Factor_A_info)){
      n = as.numeric(ki.data[i, get(Factor_A_info[j, Factor_A_name])])
      if (is.na(n) | n < Factor_A_info[j, Factor_A_threshold]){
        Factor_A[j] = 0
      } else if (n > Factor_A_info[j, Factor_A_threshold]){
        Factor_A[j] = findInterval(n, get(Factor_A_info[j, Factor_A_dist]))*3
      }
    }
    Factor_A = sum(Factor_A)
    # calculate final score #
    ki_score[i, Score := Factor_A]
  }
  return(ki_score)
}
ecs1 = ConfidenceScore_A(etki)
fecs1 = lapply(fake_etki_collection, ConfidenceScore_A)
threshold1 = data.table(Class = c("true", rep("random", 100)),
                        Score = c(mean(as.numeric(ecs1[, Score])),
                                  unlist(lapply(fecs1, function(x){mean(as.numeric(x[, Score]))}))))
threshold1_random = threshold1[Class == "random"]

fwrite(threshold1, "03 Results/05 Building ConfidenceScore/Factor A/Factor_A_ScoreResult.csv")

# plot confidence score of known vs simulated #
library(hrbrthemes)
library(viridis)
ggplot(threshold1_random, aes(x = Class, y = Score)) +
  geom_boxplot() +
  scale_fill_viridis(discrete = TRUE, alpha=0.6, option="C") +
  #ylim(c(10,18)) +
  theme(
    legend.position="none",
    plot.title = element_text(size=11)
  ) +
  geom_hline(aes(yintercept=threshold1[Class == "true", Score]), colour="#990000", linetype="dashed") +
  #scale_y_continuous(limits = c(8, 16), breaks = seq(8,16,2)) +
  theme(panel.grid.major = element_blank(), panel.grid.minor = element_blank(),
        panel.background = element_blank(), axis.line = element_line(colour = "black"))

shapiro.test(threshold1_random[, Score]) # normally distributed
t.test(threshold1_random[, Score], mu = threshold1[Class == "true", Score])$p.value

# plot number of recovery across confidence score threshold
known_recovery = data.table(Type = rep("Known", 80),
                            ConfidenceScoreCutoff = 1:80,
                            Fraction_recovery = vector("numeric", 80)
                            )
for (i in 1:nrow(known_recovery)){
  nr = ecs1[as.numeric(Score) > known_recovery[i, ConfidenceScoreCutoff], .N]/ecs1[,.N]
  known_recovery[i, Fraction_recovery := nr]
}

simulated_recovery = data.table(Type = rep("Simulated", 8000),
                                ConfidenceScoreCutoff = rep(1:80, each = 100),
                                Fraction_recovery = vector("numeric", 8000)
                                )
for (i in 1:80){
  nr = unlist(lapply(fecs1, function(x){x[as.numeric(Score) > i, .N]/x[,.N]}))
  simulated_recovery[ConfidenceScoreCutoff == i, Fraction_recovery := nr]
}

wilcox.test(known_recovery[, Fraction_recovery], simulated_recovery[, Fraction_recovery], alternative = "greater")

ggplot() +
  geom_violin(simulated_recovery, 
              mapping = aes(x = ConfidenceScoreCutoff, y = Fraction_recovery, group = ConfidenceScoreCutoff),
              color = "grey") +
  geom_line(known_recovery,
            mapping = aes(x= ConfidenceScoreCutoff, y= Fraction_recovery),
            color = "#990000",
            linetype="dashed") + 
  theme(
    legend.position="none", plot.title = element_text(size=11),
    panel.grid.major = element_blank(), panel.grid.minor = element_blank(),
    panel.background = element_blank(), axis.line = element_line(colour = "black")) +
  scale_x_continuous(breaks=seq(0,80,20)) +
  scale_y_continuous(limits = c(0, 1.0), breaks = seq(0,1.0,0.1))


recovery = rbind(known_recovery, simulated_recovery)

ggplot(recovery, aes(x = Type, y = Fraction_recovery, fill = Type)) +
  geom_violin(width=1.4) +
  geom_boxplot(width=0.05, color="black", fill = "white") +
  scale_fill_manual(values=c("#990000", "grey")) +
  theme(
    legend.position="none", plot.title = element_text(size=11),
    panel.grid.major = element_blank(), panel.grid.minor = element_blank(),
    panel.background = element_blank(), axis.line = element_line(colour = "black"))

fwrite(recovery, "03 Results/05 Building ConfidenceScore/Factor A/Factor_A_FractionRecoveryAcrossConfidence.csv")
```
