## Supplementary material for "Predicting substrates for orphan Solute Carrier Proteins using multi-omics datasets": Code for generating results: Step2_Building Factor AB.R

```
##### 0 load library and random data #####
library(data.table)
library(tidyverse)
load("~/Desktop/SLC ver3/03 Results/05 Building ConfidenceScore/Building ConfidenceScore (clean start with generated random).RData")

numbering_range = seq(0.1, 3, 0.1)
##### 1 select the best score to add #####
ConfidenceScore_AB = function(ki.data, scalar){
  ki_score = setNames(data.table(matrix(ncol = 6, nrow = nrow(ki.data))), c("HGNC_symbol","Localisation","Substrate","Formula","Status","Score"))
  char.col = colnames(ki_score)[1:5]
  num.col = colnames(ki_score)[6]
  ki_score[, (char.col) := lapply(.SD, as.character), .SDcols = char.col]
  ki_score[, (num.col) := lapply(.SD, as.character), .SDcols = num.col]
  ki_score[, 1:5 := as.list(ki.data[, 1:5])]
  #
  ## Factor A quantile
  #ns_dist1 = na.omit(etki$NCI60) # NCI60 #
  #ns_dist2 = na.omit(etki$CCLE2019) # CCLE2019 #
  #ns_dist3 = na.omit(etki$CCL180) # CCL180 #
  #q1 = quantile(ns_dist1[ns_dist1 > -1.2], seq(0, 1, by = 0.1))
  #q2 = quantile(ns_dist2[ns_dist2 > -1.4], seq(0, 1, by = 0.1))
  #q3 = quantile(ns_dist3[ns_dist3 > 1.4], seq(0, 1, by = 0.1))
  
  # Factor B quantile
  ns_dist4 = -log10(na.omit(etki$CRISPRDEP))
  q4 = quantile(ns_dist4[ns_dist4 > -log10(0.077)], seq(0, 1, by = 0.1))
  
  #Factor_A_info = data.table(Factor_A_name = c("NCI60", "CCLE2019", "CCL180"),
  #                           Factor_A_dist = c("q1", "q2", "q3"),
  #                           Factor_A_threshold = c(-1.2, -1.4, 1.4))
  for (i in 1:nrow(ki.data)){
    # score prediction for 3 matrices #
    #Factor_A = vector("integer", nrow(Factor_A_info))
    #for (j in 1:nrow(Factor_A_info)){
    #  n = as.numeric(ki.data[i, get(Factor_A_info[j, Factor_A_name])])
    #  if (is.na(n) | n < Factor_A_info[j, Factor_A_threshold]){
    #    Factor_A[j] = 0
    #  } else if (n > Factor_A_info[j, Factor_A_threshold]){
    #    Factor_A[j] = findInterval(n, get(Factor_A_info[j, Factor_A_dist]))*3
    #  }
    #}
    #Factor_A = sum(Factor_A)
    
    # Factor B: Dependency Scores #
    if (is.na(ki.data[i, CRISPRDEP]) | ki.data[i, -log10(as.numeric(CRISPRDEP))] < -log10(0.077)){
      Factor_B = 0
    } else if (ki.data[i, -log10(as.numeric(CRISPRDEP))] > -log10(0.077)){
      Factor_B = findInterval(ki.data[i, -log10(as.numeric(CRISPRDEP))], q4)*as.numeric(scalar)
    }
    # calculate final score #
    #ki_score[i, Score := Factor_A + Factor_B]
    ki_score[i, Score := Factor_B]
  }
  return(ki_score)
}
ecs_collection_AB = vector(mode = "list", length = length(numbering_range))
names(ecs_collection_AB) = numbering_range
for (i in 1:length(ecs_collection_AB)){
  ecs_collection_AB[[i]] = ConfidenceScore_AB(etki, as.numeric(names(ecs_collection_AB)[i]))
  print(i)
}
fecs_collection_AB = vector(mode = "list", length = length(numbering_range))
names(fecs_collection_AB) = numbering_range
for (i in 1:length(fecs_collection_AB)){
  fecs_collection_AB[[i]] = lapply(fake_etki_collection, function(x){ConfidenceScore_AB(x, as.numeric(names(fecs_collection_AB)[i]))})
  print(i)
}

## Collect statistical reporting in known interaction across CCL180 threshold ##
ecs_across_threshold = data.table(Threshold = numbering_range, 
                                  Status = rep("true", length(numbering_range)),
                                  Mean = vector("numeric", length(numbering_range)))
for (i in 1:length(ecs_collection_AB)){
  ecs = ecs_collection_AB[[i]]
  ecs_across_threshold[i, Mean := mean(as.numeric(ecs[,Score]))]
  print(i)
}

## Collect statistical reporting in random interaction across CCL180 threshold ##
fecs_across_threshold = data.table(Threshold = rep(numbering_range, each = 100), 
                                   Status = rep("random", length(rep(numbering_range, each = 100))),
                                   Mean = vector("numeric", length(rep(numbering_range, each = 100))))
for (i in 1:length(fecs_collection_AB)){
  fecs_confidence_score = fecs_collection_AB[[i]]
  fecs_confidence_score_mean = unlist(lapply(fecs_confidence_score, function(x){mean(as.numeric(x[,Score]))}))
  fecs_across_threshold[Threshold == names(fecs_collection_AB)[i], Mean := fecs_confidence_score_mean]
  print(i)
}

## construct table for figure ##
across_threshold = rbind(ecs_across_threshold, fecs_across_threshold)
ggplot() +
  # box plot of fecs
  geom_boxplot(data = fecs_across_threshold, 
               aes(x = Threshold, y= Mean, group = Threshold)) +
  # points of ecs
  geom_point(data = ecs_across_threshold,
             aes(x= Threshold, y= Mean),
             color = 'red')


## Generate SWEEP table ##
# Po: Points to add for an interaction with adjusted p-value < 0.077
# X: cutoff for real score
# Y: percentage of real interactions > X
# Z: percentage of fake interactions > X

Per0 = data.table(Po = NULL, 
                  X = NULL,
                  Y = NULL,
                  Z = NULL,
                  Q = NULL)

for (i in 1:length(numbering_range)){
  ecs = ecs_collection_AB[[i]]
  fecs = fecs_collection_AB[[i]]
  Per = data.table(Po = rep(names(ecs_collection_AB)[i], 40), 
                   X = 1:10,
                   Y = rep(0, 10),
                   Z = rep(0, 10),
                   Q = rep(0, 10))
  for (x in 1:10){
    y = ecs[as.numeric(Score) > x, .N]/ecs[, .N]
    z = mean(unlist(lapply(fecs, function(a){a[as.numeric(Score) > x, .N]})))/ecs[, .N]
    Per[x, Y := y]
    Per[x, Z := z]
    Per[x, Q := y-z]
  }
  Per0 = rbind(Per0, Per)
  print(i)
}

ConfABs = Per0
fwrite(ConfABs, "03 Results/05 Building ConfidenceScore/Factor AB/Sweep_factorAB_scalar.csv")

## Sweep for maximum fraction difference ##
Sweep = c()

for(i in unique(ConfABs$Po)){
  temp = ConfABs[Po == i] # temporary data table for given threshold
  #plot(temp$X, temp$Q, ylab="Fraction difference",xlab="Confidence cutoff", pch=16, main=paste("True positive vs cutoff, Th=",i, sep=""))
  MaxQ = mean(which(temp$Q==max(temp$Q)))
  MaxX = mean(temp$X[MaxQ])
  Sweep = rbind(Sweep, c(MaxX,temp$Q[MaxQ]))
}

Sweep = cbind(numbering_range,Sweep)
colnames(Sweep) = c("Po","Xmax","MaxTP")

plot(Sweep[,1],Sweep[,3], pch=18, col = "#69b3a2", xaxt="n")
axis(1, at = numbering_range)

## RESULT: a scalar of 2.0 is the points to add for independent (0,1) ##
## RESULT: a scalar of 0.1 is the points to add for interaction with adjusted p-value < 0.077 ##
```
