## Supplementary material for "Predicting substrates for orphan Solute Carrier Proteins using multi-omics datasets": Code for generating results: Step3_Building Factor ABC.R

```
##### 0 load library, random data and tool kit #####
library(data.table)
library(tidyverse)

##### 0 New random generation with derivatives #####
et <- fread("03 Results/03 Table alignment/Evidence table (normalised rho).csv")
ki <- fread("01 Raw Data/00 SLC annotation 2024/known interaction/Known interaction formula 20240321.csv")[1:837]
tb <- fread("01 Raw Data/00 SLC annotation 2024/book construction/Translation book (3 Matrix).csv")

ki <- ki[!is.na(Status)]

etki = ki
colnames(etki)[1] = "SYMBOL"
etki$Formula_info = paste(etki$Formula, etki$KEGG_ID)
etki$NCI60 = vector("numeric", nrow(etki))
etki$CCLE2019 = vector("numeric", nrow(etki))
etki$CCL180 = vector("numeric", nrow(etki))
etki$CRISPRDEP = vector("numeric", nrow(etki))

for (i in 1:nrow(etki)){
  if (is.na(etki$Formula[i])){
    etki[i, c("NCI60", "CCLE2019", "CCL180", "CRISPRDEP") := list(NA, NA, NA, NA)]
  }else if (etki$SYMBOL[i] %in% unique(et$SYMBOL)){
    etki[i, c("NCI60", "CCLE2019", "CCL180", "CRISPRDEP") :=
           as.list(et[SYMBOL == etki$SYMBOL[i] & Formula == etki$Formula[i]][,NCI60:CRISPRDEP])]
  }else{
    etki[i, c("NCI60", "CCLE2019", "CCL180", "CRISPRDEP") := list(NA, NA, NA, NA)]
  }
}

fake_etki_collection = vector("list", 100)
etki_gene_pool = unique(etki$SYMBOL)
etki_metabolite_pool = unique(etki$Formula_info)

for (sample_time in 1:100){
  # initialise fake etki data table #
  fake_etki = setNames(data.table(matrix(ncol = ncol(etki), nrow = nrow(etki))), colnames(etki))
  fake_etki[, Substrate := NULL]
  char.col = colnames(fake_etki)[1:7]
  num.col = colnames(fake_etki)[8:11]
  fake_etki[, (char.col) := lapply(.SD, as.character), .SDcols = char.col]
  fake_etki[, (num.col) := lapply(.SD, as.character), .SDcols = num.col]
  fake_etki[, c("SYMBOL", "Gene_type","Localisation") := as.list(etki[, c("SYMBOL", "Gene_type","Localisation")])]
  fake_etki[, Status := rep("Fake", nrow(fake_etki))]
  # for each SLC, shuffle fake substrate formula info #
  for (j in 1:nrow(fake_etki)){
    true_substrate = etki[SYMBOL == fake_etki$SYMBOL[j], Formula_info]
    fake_substrate_pool = etki_metabolite_pool[!etki_metabolite_pool %in% true_substrate]
    Selected = "NO"
    while(Selected == "NO"){
      fake_substrate = sample(fake_substrate_pool, 1)
      if (str_sub(fake_substrate, 1,2) == "NA"){
        fake_etki[j, Formula_info := fake_substrate]
        fake_etki[j, Formula := NA]
        fake_etki[j, KEGG_ID := str_sub(fake_substrate, 4)]
        fake_etki[j, c("NCI60", "CCLE2019", "CCL180", "CRISPRDEP") := list(NA, NA, NA, NA)]
        Selected = "YES"
      }else{
        if (fake_etki$SYMBOL[j] %in% c("SLC25A53", "SLC5A3", "SLC6A14")){
          fake_substrate_pool_specific = intersect(str_trim(fake_substrate_pool), unique(et[!is.na(NCI60)][, Formula]))
          fake_substrate = sample(fake_substrate_pool_specific, 1)
          fake_etki[j, Formula := fake_substrate]
          fake_etki[j, c("NCI60", "CCLE2019", "CCL180", "CRISPRDEP") := 
                      as.list(et[SYMBOL == fake_etki$SYMBOL[j] & Formula == fake_etki$Formula[j]][,NCI60:CRISPRDEP])]
          Selected = "YES"
        }else{
          fake_substrate = str_trim(fake_substrate)
          if (nrow(et[SYMBOL == fake_etki$SYMBOL[j] & Formula == fake_substrate][,NCI60:CRISPRDEP]) == 0){
            Selected = "NO"
          }else{
            fake_etki[j, Formula := fake_substrate]
            fake_etki[j, c("NCI60", "CCLE2019", "CCL180", "CRISPRDEP") := 
                        as.list(et[SYMBOL == fake_etki$SYMBOL[j] & Formula == fake_etki$Formula[j]][,NCI60:CRISPRDEP])]
            Selected = "YES"
          }
        }
      }
    }
    print(paste(sample_time, ":", round(j/nrow(fake_etki),2)))
  }
  fake_etki_collection[[sample_time]] = fake_etki
  print(paste(sample_time, "done"))
}

## load tool kit ##
AM = fread("03 Results/04 Adjacency/Metabolie Adjacency Matrix.csv")
colnames(AM) = gsub("cpd:", "", colnames(AM))
AM$COMPOUND = gsub("cpd:", "", AM$COMPOUND)
ConvertNames = function(name, source){
  # Process source input #
  if (is.null(source)){
    stop("Source of metabolite name required!")
  }
  else if (source %in% colnames(tb)){
    # Process name input #
    if (is.null(name)){
      stop("Name of metabolite required!")
    }else if (name %in% tb[,get(source)]){
      ans = tb[get(source) == name]
    }else{
      stop("Name entered not recognised!")
    }
  }else{
    stop("Source entered not recognised!")
  }
  
  # return ans as result #
  return(ans)
}
getAM = function(formula, step.inf = NULL, is.kegg = FALSE){
  # if the formula entered is already a kegg id (is.kegg = TRUE)
  if (isTRUE(is.kegg)){
    kegg_id = formula
  }else{
    kegg_id = as.character(ConvertNames(formula, "Formula")[,"KEGG ID"])
  }
  qc = TRUE # logical term of quality control
  generic.lipid = c("C00157", "C00165", "C00195", "C00350", "C00416", "C00422", "C00550", "C02530", "C04230", "C04438") # kegg id belonging to lipidomics
  # if kegg_id is NA in translation table #
  if(is.na(kegg_id)){
    qc = FALSE
    # if kegg_id is not NA, but not found in the adjacency matrix #
  }else if (nchar(kegg_id) == 6 & !kegg_id %in% AM$COMPOUND){
    qc = FALSE
    # if kegg_id is not NA, only has one associated kegg id, but is lipidomic #
  }else if (nchar(kegg_id) == 6 & kegg_id %in% generic.lipid){
    qc = FALSE
    # if kegg_id is not NA, only has one associated kegg id, and is found in the adjacency matrix #
  }else if (nchar(kegg_id) == 6 & kegg_id %in% AM$COMPOUND){
    kegg_id_qc = kegg_id
    # if kegg_id is not NA, but contain multiple kegg id linked with the input formula #
  }else if (nchar(kegg_id) > 6){
    kegg_id = gsub(" ", "", unlist(strsplit(kegg_id, ";"))) # break the kegg id into individual id and remove space
    kegg_id_qc = kegg_id[kegg_id %in% AM$COMPOUND & !kegg_id %in% generic.lipid] # only select the kegg id found in AM and not lipid
    # if all kegg ids are not found in AM
    if (rlang::is_empty(kegg_id_qc)){
      qc = FALSE
    }
  }else{
    qc = FALSE
  }
  
  # if compound passed qc and there's no requirement of finding infinite metabolites #
  if(qc & is.null(step.inf)){
    ans = c()
    # gather adjacent metabolite for the given kegg id #
    for (i in 1:length(kegg_id_qc)){
      AM_related = AM[COMPOUND == kegg_id_qc[i]]
      AM_related = AM_related[,.SD,.SDcols=which(AM_related[,.SD] != Inf & AM_related[,.SD] != 0)]
      # if there is no adjacent metabolite derivative for the given kegg id #
      if (ncol(AM_related) == 1){
        ans_temp = character(0)
        # if there is adjacent metabolite derivative for the given kegg id #
      }else{
        ans_temp = colnames(AM_related)[2:ncol(AM_related)]
        names(ans_temp) = as.numeric(AM_related[,2:ncol(AM_related)])
        ans = c(ans, ans_temp)
      }
    }
    ans = ans[!ans %in% generic.lipid] # remove lipid derivatives in the answer
    # if the accumulated number of adjacent metabolite derivative equals to 0 #
    if (is.null(ans)){
      return(NA)
      # if the accumulated number of adjacent metabolite derivative does not equals to 0 #
    }else{
      return(ans)
    }
    # if compound passed qc and there's requirement of finding infinite metabolites #
  }else if (qc & !is.null(step.inf)){
    ans = vector("list", length(kegg_id_qc))
    # gather infinite metabolite for the given kegg id #
    for (i in 1:length(kegg_id_qc)){
      AM_related = AM[COMPOUND == kegg_id_qc[i]]
      AM_related = AM_related[,.SD,.SDcols=which(AM_related[,.SD] == Inf & AM_related[,.SD] != 0)]
      ans[[i]] = colnames(AM_related)[2:ncol(AM_related)]
    }
    # if there is only 1 kegg id passing qc, return the info of infinite metabolites #
    if (length(ans) == 1){
      return(ans[[1]])
      # if there is multiple kegg ids passing qc, return the intersect of the infinite metabolites #
    }else{
      ans_intersect = intersect(ans[[1]], ans[[2]])
      if (length(ans) == 2){
        return(ans_intersect)
      }else{
        for (j in 3:length(ans)){
          ans_intersect = intersect(ans_intersect, ans[[j]])
        }
        return(ans_intersect)
      }
    }
    # if compound did not pass qc #
  }else{
    return(NA)
  }
}
translate = function(KEGG.id){
  kegg_info = data.table(KEGG = as.character(KEGG.id),
                         Formula = vector("character", length(KEGG.id)))
  for (j in 1:nrow(kegg_info)){
    info = grep(kegg_info$KEGG[j], tb$`KEGG ID`)
    if (length(info) == 0){
      kegg_info[j, Formula := NA]
    }else if (length(info) == 1){
      kegg_info[j, Formula := tb$Formula[info]]
    }else if (length(info) > 1){
      kegg_info[j, Formula := paste(tb$Formula[info], collapse = "<>")]
    }
  }
  # Remove the derivative that's not found in the translation book #
  kegg_info = na.omit(kegg_info)
  # if no derivative is found in the translation book, stop translation #
  if (nrow(kegg_info) == 0){
    return(NA)
    # if there's only one formula matching the given kegg
  }else if (length(grep("<>", kegg_info$Formula)) == 0){
    return(kegg_info)
  }else{
    # Break the <> and insert a new row #
    kegg_info = kegg_info |> 
      mutate(Formula = strsplit(as.character(Formula), "<>")) |> 
      unnest(Formula) |>
      group_by(Formula) |>
      summarise(KEGG = toString(sort(unique(KEGG)), .groups='drop'))
    kegg_info = as.data.table(kegg_info)
    return(kegg_info)
  }
}
getNR = function(gene, translated.formula){
  data = data.table(Formula = translated.formula[, Formula],
                    NCI60 = vector("numeric", nrow(translated.formula)),
                    CCLE2019 = vector("numeric", nrow(translated.formula)),
                    CCL180 = vector("numeric", nrow(translated.formula)))
  for (k in 1:nrow(data)){
    if(nrow(et[SYMBOL == gene & Formula == data[k, Formula]]) != 0){
      data[k, c("NCI60", "CCLE2019", "CCL180") :=
             as.list(et[SYMBOL == gene & Formula == data[k, Formula]][,NCI60:CCL180])]
    }else{
      data[k, c("NCI60", "CCLE2019", "CCL180") := as.list(NA, NA, NA)]
    }
  }
  return(data)
}
check_NA = function(x){
  if (length(x) == 1){
    if (is.na(x)){
      return(TRUE)
    }else{
      return(FALSE)
    }
  }else{
    return(FALSE)
  }
}

## Start package for re-visit ##
load("~/Desktop/SLC ver3/03 Results/05 Building ConfidenceScore/Building ConfidenceScore ABC (clean start with generated random).RData")

##### 3 Necessity of considering CCL180 derivatives in lower range #####
ConfidenceScore_ABCz = function(ki.data, threshold){
  ki_score = setNames(data.table(matrix(ncol = 9, nrow = nrow(ki.data))), 
                      c("SYMBOL","Gene_type","Localisation","Substrate","Formula","KEGG_ID","Status","Formula_info", "Score")
  )
  char.col = colnames(ki_score)[1:8]
  num.col = colnames(ki_score)[9]
  ki_score[, (char.col) := lapply(.SD, as.character), .SDcols = char.col]
  ki_score[, (num.col) := lapply(.SD, as.character), .SDcols = num.col]
  ki_score[, 1:8 := as.list(ki.data[, 1:8])]
  
  data_info = data.table(data_name = c("NCI60", "CCLE2019", "CCL180"),
                         data_dist = c("q1", "q2", "q3"),
                         Factor_A_threshold = c(0.0, 0.0, 0.8),
                         Factor_C_threshold = c(0.0, 0.0, threshold))
  
  # Factor A quantile
  ns_dist1 = na.omit(etki$NCI60) # NCI60 #
  ns_dist2 = na.omit(etki$CCLE2019) # CCLE2019 #
  ns_dist3 = na.omit(etki$CCL180) # CCL180 #
  q1 = quantile(ns_dist1[ns_dist1 > 0.0], seq(0, 1, by = 0.1))
  q2 = quantile(ns_dist2[ns_dist2 > 0.0], seq(0, 1, by = 0.1))
  q3 = quantile(ns_dist3[ns_dist3 > 0.8], seq(0, 1, by = 0.1))
  # Factor B quantile
  ns_dist4 = -log10(na.omit(etki$CRISPRDEP))
  q4 = quantile(ns_dist4[ns_dist4 > -log10(0.077)], seq(0, 1, by = 0.1))
  
  for (i in 1:nrow(ki.data)){
    # score prediction for 3 matrices #
    Factor_A = vector("integer", nrow(data_info))
    for (j in 1:nrow(data_info)){
      n = ki.data[i, get(data_info[j, data_name])]
      if (is.na(n) | n < data_info[j, Factor_A_threshold]){
        Factor_A[j] = 0
      } else if (n > data_info[j, Factor_A_threshold]){
        Factor_A[j] = findInterval(n, get(data_info[j, data_dist]))*3
      }
    }
    Factor_A = sum(Factor_A)
    
    # Factor B: Dependency Scores #
    if (is.na(ki.data[i, CRISPRDEP]) | ki.data[i, -log10(as.numeric(CRISPRDEP))] < -log10(0.077)){
      Factor_B = 0
    } else if (ki.data[i, -log10(as.numeric(CRISPRDEP))] > -log10(0.077)){
      Factor_B = findInterval(ki.data[i, -log10(as.numeric(CRISPRDEP))], q4)*1.1
    }
    
    # Factor C: Adjacency #
    if (is.na(ki.data$Formula[i])){
      temp = getAM(ki.data$KEGG_ID[i], is.kegg = TRUE)
    }else{
      temp = getAM(ki.data$Formula[i])
    }
    if (check_NA(temp)){
      Factor_C = 0
    }else if (!1 %in% names(temp) & !2 %in% names(temp)){
      Factor_C = 0
    }else{
      proximal = translate(temp[which(names(temp) %in% c(1,2))])
      if (check_NA(proximal)){
        Factor_C = 0
      }else{
        proximal_ns = getNR(ki.data[i, SYMBOL], proximal)
        if (nrow(proximal_ns) != 0) {
          Factor_C = 0
          for (l in 1:nrow(data_info)){
            N = unique(proximal_ns[, get(data_info[l, data_name])])
            for (n in N){
              if (is.na(n) | n < data_info[l, Factor_C_threshold]){
                Factor_C = Factor_C + 0
              }else if (n > data_info[l, Factor_C_threshold] & n < data_info[l, Factor_A_threshold]){
                Factor_C = Factor_C + 0.9/2
              }else if (n > data_info[l, Factor_A_threshold]){
                Factor_C = Factor_C + findInterval(n, get(data_info[l, data_dist]))*0.9
              }
            }
          }
        }else{
          Factor_C = 0
        }
      }
    }
    # calculate final score #
    ki_score[i, Score := Factor_A + Factor_B + Factor_C]
  }
  return(ki_score)
}

## Calculate Confidence Score for known and random interactions, changing CCLE2019 threshold ##
ecs_collection_ABCz = vector(mode = "list", length = 9)
names(ecs_collection_ABCz) = seq(0, 0.8, 0.1)
for (i in 1:length(ecs_collection_ABCz)){
  ecs_collection_ABCz[[i]] = ConfidenceScore_ABCz(etki, names(ecs_collection_ABCz)[i])
  print(i)
}
fecs_collection_ABCz = vector(mode = "list", length = 9)
names(fecs_collection_ABCz) = seq(0, 0.8, 0.1)
for (i in 1:length(fecs_collection_ABCz)){
  fecs_collection_ABCz[[i]] = lapply(fake_etki_collection, function(x){ConfidenceScore_ABCz(x, names(fecs_collection_ABCz)[i])})
  print(i)
}

## Collect statistical reporting in known interaction across CCLE2019 threshold ##
ecs_across_threshold = data.table(Threshold = seq(0, 0.8, 0.1), 
                                  Status = rep("true", length(seq(0, 0.8, 0.1))),
                                  Mean = vector("numeric", length(seq(0, 0.8, 0.1))))
for (i in 1:length(ecs_collection_ABCz)){
  ecs = ecs_collection_ABCz[[i]]
  ecs_across_threshold[i, Mean := mean(as.numeric(ecs[,Score]))]
  print(i)
}

## Collect statistical reporting in random interaction across CCLE2019 threshold ##
fecs_across_threshold = data.table(Threshold = rep(seq(0, 0.8, 0.1), each = 100), 
                                   Status = rep("random", length(rep(seq(0, 0.8, 0.1), each = 100))),
                                   Mean = vector("numeric", length(rep(seq(0, 0.8, 0.1), each = 100))))
for (i in 1:length(fecs_collection_ABCz)){
  fecs_confidence_score = fecs_collection_ABCz[[i]]
  fecs_confidence_score_mean = unlist(lapply(fecs_confidence_score, function(x){mean(as.numeric(x[,Score]))}))
  fecs_across_threshold[Threshold == names(fecs_collection_ABCz)[i], Mean := fecs_confidence_score_mean]
  print(i)
}

## construct table for Boxplot figure ##
across_threshold_ABCz = rbind(ecs_across_threshold, fecs_across_threshold)
fwrite(across_threshold_ABCz, "03 Results/05 Building ConfidenceScore/Factor ABC/across threshold_factorABCz.csv")

# Across all derivative threshold selected for CCLE2019, known interaction behave better than random #
ggplot() +
  # box plot of fecs
  geom_boxplot(data = fecs_across_threshold, 
               aes(x = Threshold, y= Mean, group = Threshold)) +
  # points of ecs
  geom_point(data = ecs_across_threshold,
             aes(x= Threshold, y= Mean),
             color = 'red')

## Generate SWEEP table ##
# Th: Predictive score cutoff for the interaction to be marked as "discovered"
# X: cutoff for real score
# Y: percentage of real interactions > X
# Z: percentage of fake interactions > X

Per0 = data.table(Th = NULL, 
                  X = NULL,
                  Y = NULL,
                  Z = NULL,
                  Q = NULL)

for (i in 1:9){
  ecs = ecs_collection_ABCz[[i]]
  fecs = fecs_collection_ABCz[[i]]
  Per = data.table(Th = rep(names(ecs_collection_ABCz)[i], 40), 
                   X = 1:40,
                   Y = rep(0, 40),
                   Z = rep(0, 40),
                   Q = rep(0, 40))
  for (x in 1:40){
    y = ecs[as.numeric(Score) > x, .N]/ecs[, .N]
    z = mean(unlist(lapply(fecs, function(a){a[as.numeric(Score) > x, .N]})))/ecs[, .N]
    Per[x, Y := y]
    Per[x, Z := z]
    Per[x, Q := y-z]
  }
  Per0 = rbind(Per0, Per)
  print(i)
}

ConfABCz = Per0
fwrite(ConfABCz, "03 Results/05 Building ConfidenceScore/Factor ABC/Sweep_factorABCz.csv")

## Sweep for maximum fraction difference ##
Sweep = c()
for(i in unique(ConfABCz$Th)){
  temp = ConfABCz[Th == i] # temporary data table for given threshold
  #plot(temp$X, temp$Q, ylab="Fraction difference",xlab="Confidence cutoff", pch=16, main=paste("True positive vs cutoff, Th=",i, sep=""))
  MaxQ = mean(which(temp$Q==max(temp$Q)))
  MaxX = mean(temp$X[MaxQ])
  Sweep = rbind(Sweep, c(MaxX,temp$Q[MaxQ]))
}

Sweep = cbind(seq(0, 0.8, 0.1),Sweep)
colnames(Sweep) = c("Th","Xmax","MaxTP")

plot(Sweep[,1],Sweep[,3], pch=18, col = "#69b3a2", xaxt="n")
axis(1, at = seq(0, 0.8, 0.1))

## RESULT: use 0.1 for CCL180 threshold ##

#######################################################################################################################

##### 4 What score to add for predicted derivatives? #####
ConfidenceScore_ABCp = function(ki.data, scalar){
  ki_score = setNames(data.table(matrix(ncol = 9, nrow = nrow(ki.data))), 
                      c("SYMBOL","Gene_type","Localisation","Substrate","Formula","KEGG_ID","Status","Formula_info", "Score")
                      )
  char.col = colnames(ki_score)[1:8]
  num.col = colnames(ki_score)[9]
  ki_score[, (char.col) := lapply(.SD, as.character), .SDcols = char.col]
  ki_score[, (num.col) := lapply(.SD, as.character), .SDcols = num.col]
  ki_score[, 1:8 := as.list(ki.data[, 1:8])]
  
  data_info = data.table(data_name = c("NCI60", "CCLE2019", "CCL180"),
                         data_dist = c("q1", "q2", "q3"),
                         Factor_A_threshold = c(0.0, 0.0, 0.0),
                         Factor_C_threshold = c(0.0, 0.0, 0.0))
  
  # Factor A quantile
  ns_dist1 = na.omit(etki$NCI60) # NCI60 #
  ns_dist2 = na.omit(etki$CCLE2019) # CCLE2019 #
  ns_dist3 = na.omit(etki$CCL180) # CCL180 #
  q1 = quantile(ns_dist1[ns_dist1 > 0.0], seq(0, 1, by = 0.1))
  q2 = quantile(ns_dist2[ns_dist2 > 0.0], seq(0, 1, by = 0.1))
  q3 = quantile(ns_dist3[ns_dist3 > 0.0], seq(0, 1, by = 0.1))
  # Factor B quantile
  ns_dist4 = -log10(na.omit(etki$CRISPRDEP))
  q4 = quantile(ns_dist4[ns_dist4 > -log10(0.077)], seq(0, 1, by = 0.1))
  
  for (i in 1:nrow(ki.data)){
    # score prediction for 3 matrices #
    Factor_A = vector("integer", nrow(data_info))
    for (j in 1:nrow(data_info)){
      n = ki.data[i, get(data_info[j, data_name])]
      if (is.na(n) | n < data_info[j, Factor_A_threshold]){
        Factor_A[j] = 0
      } else if (n > data_info[j, Factor_A_threshold]){
        Factor_A[j] = findInterval(n, get(data_info[j, data_dist]))*3
      }
    }
    Factor_A = sum(Factor_A)
    
    # Factor B: Dependency Scores #
    if (is.na(ki.data[i, CRISPRDEP]) | ki.data[i, -log10(as.numeric(CRISPRDEP))] < -log10(0.077)){
      Factor_B = 0
    } else if (ki.data[i, -log10(as.numeric(CRISPRDEP))] > -log10(0.077)){
      Factor_B = findInterval(ki.data[i, -log10(as.numeric(CRISPRDEP))], q4)*1.6
    }
    
    # Factor C: Adjacency #
    if (is.na(ki.data$Formula[i])){
      temp = getAM(ki.data$KEGG_ID[i], is.kegg = TRUE)
    }else{
      temp = getAM(ki.data$Formula[i])
    }
    if (check_NA(temp)){
      Factor_C = 0
    }else if (!1 %in% names(temp) & !2 %in% names(temp)){
      Factor_C = 0
    }else{
      proximal = translate(temp[which(names(temp) %in% c(1,2))])
      if (check_NA(proximal)){
        Factor_C = 0
      }else{
        proximal_ns = getNR(ki.data[i, SYMBOL], proximal)
        if (nrow(proximal_ns) != 0) {
          Factor_C = 0
          for (l in 1:nrow(data_info)){
            N = unique(proximal_ns[, get(data_info[l, data_name])])
            for (n in N){
              if (is.na(n) | n < data_info[l, Factor_C_threshold]){
                Factor_C = Factor_C + 0
              }else if (n > data_info[l, Factor_C_threshold] & n < data_info[l, Factor_A_threshold]){
                Factor_C = Factor_C + scalar/2
              }else if (n > data_info[l, Factor_A_threshold]){
                Factor_C = Factor_C + findInterval(n, get(data_info[l, data_dist]))*scalar
              }
            }
          }
        }else{
          Factor_C = 0
        }
      }
    }
    # calculate final score #
    ki_score[i, Score := Factor_A + Factor_B + Factor_C]
  }
  return(ki_score)
}

## Calculate Confidence Score for known and random interactions, changing CCLE2019 threshold ##
ecs_collection_ABCp = vector(mode = "list", length = 10)
names(ecs_collection_ABCp) = seq(0.1, 1, 0.1)
for (i in 1:length(ecs_collection_ABCp)){
  ecs_collection_ABCp[[i]] = ConfidenceScore_ABCp(etki, as.numeric(names(ecs_collection_ABCp)[i]))
  print(i)
}
fecs_collection_ABCp = vector(mode = "list", length = 10)
names(fecs_collection_ABCp) = seq(0.1, 1, 0.1)
for (i in 1:length(fecs_collection_ABCp)){
  fecs_collection_ABCp[[i]] = lapply(fake_etki_collection, function(x){ConfidenceScore_ABCp(x, as.numeric(names(fecs_collection_ABCp)[i]))})
  print(i)
}

## Generate SWEEP table ##
# Po: Points to add for each interaction marked as predicted
# X: cutoff for real score
# Y: percentage of real interactions > X
# Z: percentage of fake interactions > X

Per0 = data.table(Po = NULL, 
                  X = NULL,
                  Y = NULL,
                  Z = NULL,
                  Q = NULL)

for (i in 1:10){
  ecs = ecs_collection_ABCp[[i]]
  fecs = fecs_collection_ABCp[[i]]
  Per = data.table(Po = rep(names(ecs_collection_ABCp)[i], 40), 
                   X = 1:40,
                   Y = rep(0, 40),
                   Z = rep(0, 40),
                   Q = rep(0, 40))
  for (x in 1:40){
    y = ecs[as.numeric(Score) > x, .N]/ecs[, .N]
    z = mean(unlist(lapply(fecs, function(a){a[as.numeric(Score) > x, .N]})))/ecs[, .N]
    Per[x, Y := y]
    Per[x, Z := z]
    Per[x, Q := y-z]
  }
  Per0 = rbind(Per0, Per)
  print(i)
}

ConfABCp = Per0

fwrite(ConfABCp, "03 Results/05 Building ConfidenceScore/Factor ABC/Sweep_factorABCp.csv")

## Sweep for maximum fraction difference ##
Sweep = c()
for(i in unique(ConfABCp$Po)){
  temp = ConfABCp[Po == i] # temporary data table for given threshold
  #plot(temp$X, temp$Q, ylab="Fraction difference",xlab="Confidence cutoff", pch=16, main=paste("True positive vs cutoff, Th=",i, sep=""))
  MaxQ = mean(which(temp$Q==max(temp$Q)))
  MaxX = mean(temp$X[MaxQ])
  Sweep = rbind(Sweep, c(MaxX,temp$Q[MaxQ]))
}

Sweep = cbind(seq(0.1,1,0.1),Sweep)
colnames(Sweep) = c("Po","Xmax","MaxTP")

plot(Sweep[,1],Sweep[,3], pch=18, col = "#69b3a2", xaxt="n")
axis(1, at = seq(0.1,1,0.1))

# use 0.9 as the score to add
```
