## Supplementary material for "Predicting substrates for orphan Solute Carrier Proteins using multi-omics datasets": Code for generating results: Step5_Survey Confidence Score.R

```
##### 0 load library #####
library(data.table)
library(tidyverse)
library(hrbrthemes)
library(viridis)

##### 1 read files #####
etcs = fread("03 Results/05 Building ConfidenceScore/Evidence table (Confidence Score).csv")
known_list = fread("01 Raw Data/00 SLC annotation 2024/known interaction/Known interaction formula 20240321.csv")[1:841]
orphan_list = fread("01 Raw Data/00 SLC annotation 2024/SLC_transcript_20240321.csv", select = c(1:6))[Knowledge_type == "Orphan"]

##### 2 is rank in known set prone to appear closer to start? #####
etcs_ki = known_list[!is.na(Formula)]
colnames(etcs_ki)[1] = "SYMBOL"
etcs_ki = data.table(etcs_ki,
                     Rank = vector("numeric", nrow(etcs_ki)),
                     A = vector("numeric", nrow(etcs_ki)),
                     B = vector("numeric", nrow(etcs_ki)),
                     C = vector("numeric", nrow(etcs_ki)),
                     Score = vector("numeric", nrow(etcs_ki)))
for (i in 1:nrow(etcs_ki)){
  if (etcs_ki[i, SYMBOL] %in% unique(etcs$SYMBOL)){
    #etcs_atomic = etcs[SYMBOL == etcs_ki$SYMBOL[i]][order(A, Score, decreasing = TRUE)]
    etcs_atomic = etcs[SYMBOL == etcs_ki$SYMBOL[i]][order(Score, A, decreasing = TRUE)]
    #etcs_atomic = etcs[SYMBOL == etcs_ki$SYMBOL[i]][order(Score, decreasing = TRUE)]
    etcs_ki[i, c("A", "B", "C", "Score") := etcs_atomic[Formula == etcs_ki$Formula[i], c("A", "B", "C", "Score")]]
    etcs_ki[i, Rank := which(etcs_atomic$Formula == etcs_ki$Formula[i])]
  }else{
    etcs_ki[i, Rank := NaN]
  }
}

## Generate fake etcs_ki ##
fake_collection = vector("list", 100)
etcs_ki_gene_pool = unique(etcs_ki$SYMBOL)
etcs_ki_metabolite_pool = unique(etcs_ki$Formula)

for (sample_time in 1:100){
  # initialise fake etcs_ki data table #
  fake_etcs_ki = setNames(data.table(matrix(ncol = ncol(etcs_ki), nrow = nrow(etcs_ki))), colnames(etcs_ki))
  fake_etcs_ki[, Substrate := NULL]
  char.col = colnames(fake_etcs_ki)[1:6]
  num.col = colnames(fake_etcs_ki)[7]
  fake_etcs_ki[, (char.col) := lapply(.SD, as.character), .SDcols = char.col]
  fake_etcs_ki[, (num.col) := lapply(.SD, as.character), .SDcols = num.col]
  fake_etcs_ki[, c("SYMBOL", "Gene_type","Localisation") := as.list(etcs_ki[, c("SYMBOL", "Gene_type","Localisation")])]
  fake_etcs_ki[, Status := rep("Fake", nrow(fake_etcs_ki))]
  # for each SLC, shuffle fake substrate formula #
  for (j in 1:nrow(fake_etcs_ki)){
    if (!fake_etcs_ki[j, SYMBOL] %in% unique(etcs[, SYMBOL])){
      fake_etcs_ki[j, Rank := NaN]
      print(paste(sample_time, ":", round(j/nrow(fake_etcs_ki),2)))
    }else{
      true_substrate = etcs_ki[SYMBOL == fake_etcs_ki$SYMBOL[j]][, Formula]
      fake_substrate_pool = etcs_ki_metabolite_pool[!etcs_ki_metabolite_pool %in% true_substrate]
      fake_substrate = sample(fake_substrate_pool, 1)
      while (nrow(etcs[SYMBOL == fake_etcs_ki$SYMBOL[j] & Formula == fake_substrate]) == 0){
        fake_substrate = sample(fake_substrate_pool, 1)
      }
      fake_etcs_ki[j, Formula := fake_substrate]
      etcs_atomic = etcs[SYMBOL == etcs_ki$SYMBOL[i]][order(Score, A, decreasing = TRUE)]
      #etcs_atomic = etcs[SYMBOL == fake_etcs_ki$SYMBOL[j]][order(A, Score, decreasing = TRUE)]
      #etcs_atomic = etcs[SYMBOL == fake_etcs_ki$SYMBOL[j]][order(Score, decreasing = TRUE)]
      fake_etcs_ki[j, Rank := which(etcs_atomic$Formula == fake_etcs_ki$Formula[j])]
      print(paste(sample_time, ":", round(j/nrow(fake_etcs_ki),2)))
    }
  }
  fake_collection[[sample_time]] = fake_etcs_ki
  print(paste(sample_time, "done"))
}

fake_median = unlist(lapply(1:100, function(x){fake_collection[[x]][, median(as.numeric(Rank), na.rm = T)]}))
known_median = etcs_ki[, median(Rank, na.rm = T)]
median_benchmark = data.table(Class = c("true", rep("simulated", 100)),
                              Median = c(known_median, fake_median))
ggplot(median_benchmark[Class == "simulated"], aes(x = Class, y = Median)) +
  geom_boxplot(fill = "grey") +
  scale_fill_viridis(discrete = TRUE, alpha=0.6, option="C") +
  #ylim(c()) +
  theme(
    legend.position="none",
    plot.title = element_text(size=11)
  ) +
  geom_hline(aes(yintercept=median_benchmark[Class == "true", Median]), colour="#990000", linetype="dashed") +
  theme(panel.grid.major = element_blank(), panel.grid.minor = element_blank(),
        panel.background = element_blank(), axis.line = element_line(colour = "black"))
  #scale_y_continuous(limits = c(150, 400), breaks=seq(150, 400,50)) 
  #expand_limits()

shapiro.test(median_benchmark[Class == "simulated", Median]) # not normally distributed
wilcox.test(median_benchmark[Class == "simulated", Median], mu = median_benchmark[Class == "true", Median], alternative = "g")$p.value

##### 3 the best rank cutoff to use #####
Sweep = data.table(X = rep(1:500, 100),
                   Q = vector("numeric", 50000))
for (x in 1:500){
  y = etcs_ki[Rank <= x, .N]/etcs_ki[,.N]
  z = unlist(lapply(1:100, function(a){fake_collection[[a]][as.numeric(Rank) <= x, .N]/fake_collection[[a]][,.N]}))
  q = y-z
  Sweep[X == x, Q := q]
}

ggplot() +
  geom_violin(Sweep, mapping = aes(x = X, y = Q, group = X, alpha = 0.2), color = "grey", fill = "grey") +
  geom_line(data.table(X = 1:500, Q = Sweep[, median(Q), by = X][, V1]), mapping = aes(x = X, y = Q)) +
  theme(
    legend.position="none", plot.title = element_text(size=11),
    panel.grid.major = element_blank(), panel.grid.minor = element_blank(),
    panel.background = element_blank(), axis.line = element_line(colour = "black"))


for (x in 178){
  y = etcs_ki[Rank <= x, .N]/etcs_ki[,.N]
  z = unlist(lapply(1:100, function(a){fake_collection[[a]][as.numeric(Rank) <= x, .N]/fake_collection[[a]][,.N]}))
  q = y-z
  #Sweep[X == x, Q := median(q)]
}


mean(z)
2*sd(z)

## use rank <= 182 ##
```
