## Supplementary material for "Predicting substrates for orphan Solute Carrier Proteins using multi-omics datasets": Code for generating results: Step6_ORA.R

```
library(data.table)
library(tidyverse)
library(KEGGREST)

##### 1 generate list of human metabolic pathways and linked compound to each pathways #####
# get human pathways (metabolic pathway definition from CCL180 paper)
human_metabolic_pathways = fread("01 Raw Data/05 Building ConfidenceScore/Pathways.csv")

# in data table format for re-visit
human_metabolic_pathways = data.table(human_metabolic_pathways,
                                      linked_compound = vector("character", length = nrow(human_metabolic_pathways)),
                                      compound_no = vector("numeric", length = nrow(human_metabolic_pathways))
                                      )
for (i in 1:nrow(human_metabolic_pathways)){
  q = keggGet(human_metabolic_pathways[i, "Pathway id"])[[1]][["COMPOUND"]]
  id = paste(names(q), collapse = ";")
  no = length(q)
  human_metabolic_pathways[i, linked_compound := id]
  human_metabolic_pathways[i, compound_no := no]
  print(i)
}

# in list format for ORA (function category list for ORA)
human_metabolic_pathways_list = vector("list", nrow(human_metabolic_pathways))
names(human_metabolic_pathways_list) = unlist(human_metabolic_pathways[, "Pathway id"])
for (i in 1:length(human_metabolic_pathways_list)){
  human_metabolic_pathways_list[[i]] = names(keggGet(names(human_metabolic_pathways_list)[i])[[1]][["COMPOUND"]])
  print(i)
}

##### 2 gather feature parameter for ORA #####
etcs = fread("03 Results/05 Building ConfidenceScore/Evidence table (Confidence Score).csv")
tb = fread("01 Raw Data/00 SLC annotation 2024/book construction/Translation book (3 Matrix).csv")
known_list = fread("01 Raw Data/00 SLC annotation 2024/known interaction/Known interaction formula 20240321.csv")[1:841][!is.na(Status)]
orphan_list = fread("01 Raw Data/00 SLC annotation 2024/SLC_transcript_20240321.csv", select = c(1:6))[Knowledge_type == "Orphan"]

##### 3 ORA #####
# Lazy fisher's exact test
# test against the hypothesis that the feature does not positively linked to category
fisherExactTest <- function(features, category, universe){
  contingencyTable <- matrix(nrow = 2, ncol = 2)
  rownames(contingencyTable) <- c("inCategory", "outCategory")
  colnames(contingencyTable) <- c("inFeature", "outFeature")
  # prepare values
  x <- sum(features %in% category)
  k <- length(category)
  m <- length(features)
  N <- length(universe)
  n <- N - m
  # fill contingency table
  contingencyTable["inCategory", "inFeature"] <- x
  contingencyTable["inCategory", "outFeature"] <- k - x
  contingencyTable["outCategory", "inFeature"] <- m - x
  contingencyTable["outCategory", "outFeature"] <- n - (k - x)
  # apply fisher exact test
  p <- stats::fisher.test(x = contingencyTable, alternative = "greater")$p.value 
  # there are MORE number in the overlap of feature and category (top left cells)
  return(p)
}
# Over-representation analysis
overRepresentationAnalysis <- function(features, funCatList, universe,
                                       minSize = 1, maxSize = Inf,
                                       removeNoOverlap = TRUE, pAdjustMethod = "BH") {
  
  # if missing universe, use all features in the functional category list
  if(missing(universe)) {
    universe <- unique(unlist(funCatList))
  }
  # remove categories out of size
  size <- lapply(funCatList, length) %>%
    unlist()
  funCatList <- funCatList[size >= minSize & size <= maxSize]
  # calculate overlap and p value for each funCat, returning a one-row data frame with results
  dfList <- lapply(funCatList, function(funCat) {
    
    overlappingFeatures <- features[features %in% funCat]
    nOverlap <- length(overlappingFeatures)
    if(nOverlap == 0) {
      overlappingFeatures <- "No overlap"
    } else {
      overlappingFeatures <- paste(unique(overlappingFeatures), collapse = ", ")
    }
    # get enrichment p values
    fisherP <- fisherExactTest(features = features, category = funCat, universe = universe)
    # return out df
    outDf <- data.frame(overlap = overlappingFeatures, nOverlap = nOverlap, pValue = fisherP)
    return(outDf)
    
  })
  # bind rows
  resDf <- dplyr::bind_rows(dfList, .id = "functionalCategory")
  # adjust p values
  if(removeNoOverlap) {
    resDf <- subset(resDf, resDf$nOverlap != 0)
  }
  resDf[ , "pAdj"] <- stats::p.adjust(resDf$pValue, method = pAdjustMethod)
  # return data frame
  return(resDf)
}
# toolkit
ConvertNames = function(name, source){
  # Process source input #
  if (is.null(source)){
    stop("Source of metabolite name required!")
  }
  else if (source %in% colnames(tb)){
    # Process name input #
    if (is.null(name)){
      stop("Name of metabolite required!")
    }else if (name %in% tb[,get(source)]){
      ans = tb[get(source) == name]
    }else{
      stop("Name entered not recognised!")
    }
  }else{
    stop("Source entered not recognised!")
  }
  
  # return ans as result #
  return(ans)
}
check_NA = function(x){
  if (length(x) == 1){
    if (is.na(x)){
      return(TRUE)
    }else{
      return(FALSE)
    }
  }else{
    return(FALSE)
  }
}

# Universe
linked_to_metabolomics = unique(str_trim(unlist(str_split(unlist(tb[, "KEGG ID"]), ";"))))
linked_to_pathway = unique(str_trim(unlist(str_split(unlist(human_metabolic_pathways[, linked_compound]), ";"))))
univ = intersect(linked_to_metabolomics, linked_to_pathway)

## build known list for ORA ##
known_list_ORA = data.table(SYMBOL = known_list[, unique(HGNC_symbol)],
                            Substrate = vector("character", length(known_list[, unique(HGNC_symbol)])),
                            KEGG_ID = vector("character", length(known_list[, unique(HGNC_symbol)])))
for (i in 1:nrow(known_list_ORA)){
  s = paste(known_list[HGNC_symbol == known_list_ORA[i, SYMBOL], Substrate], collapse = "; ")
  f = known_list[HGNC_symbol == known_list_ORA[i, SYMBOL], Formula]
  if (length(which(is.na(f))) == 0){
    k = paste(na.omit(unlist(lapply(f, function(x){as.character(ConvertNames(x, "Formula")[,"KEGG ID"])}))), collapse = "; ")
  }else{
    kna_YES = known_list[HGNC_symbol == known_list_ORA[i, SYMBOL] & is.na(Formula), KEGG_ID]
    kna_NO = paste(na.omit(unlist(lapply(f[-which(is.na(f))], function(x){as.character(ConvertNames(x, "Formula")[,"KEGG ID"])}))), collapse = "; ")
    k = paste(c(kna_YES, kna_NO), collapse = "; ")
  }
  if (nchar(k) == 8 & str_sub(k, 7,8) == "; "){
    k = str_remove(k, "; ")
  }
  known_list_ORA[i, Substrate := s]
  known_list_ORA[i, KEGG_ID := k]
}

bulk.overRepresentationAnalysis.known = function(known_list_ORA, up.to.rank, set.p.value = 0.05){
  # Calculate ORA
  known_list_ORA_result = vector("list", nrow(known_list_ORA))
  names(known_list_ORA_result) = known_list_ORA[, SYMBOL]
  for (i in 1:nrow(known_list_ORA)){
    etcs_atomic = etcs[SYMBOL == known_list_ORA[i, SYMBOL]][order(Score, A, decreasing = TRUE)][1:up.to.rank]
    etcs_atomic_feature = 
      lapply(etcs_atomic[,na.omit(Formula)], function(x){as.character(ConvertNames(x, "Formula")[,"KEGG ID"])}) |>
      unlist() |>
      str_split(";") |>
      unlist() |>
      str_trim()
    
    ora_reporting = overRepresentationAnalysis(features = unique(etcs_atomic_feature), 
                                               funCatList = human_metabolic_pathways_list, 
                                               universe = univ)
    ora_reporting = inner_join(x = data.table(ora_reporting)[pAdj <= set.p.value][order(pAdj)][,pValue := NULL], 
                               y = human_metabolic_pathways[, .(functionalCategory = `Pathway id`, `Pathway name`, Class, compound_no)], 
                               by = "functionalCategory")
    setcolorder(ora_reporting, c("functionalCategory", "Pathway name", "Class", "pAdj", "compound_no", "nOverlap", "overlap"))
    
    known_list_ORA_result[[i]] = ora_reporting
  }
  # Specify result
  known_list_ORA = data.table(known_list_ORA,
                              Status = vector("character", nrow(known_list_ORA)))
  for (i in 1:length(known_list_ORA_result)){
    pathway = inner_join(x = data.table("Pathway id" = known_list_ORA_result[[i]][,functionalCategory]),
                         y = human_metabolic_pathways[, .(`Pathway id`, linked_compound)],
                         by = "Pathway id")
    if (nrow(pathway) == 0){
      known_list_ORA[i, Status := "No Pathway Enriched"]
    }else{
      pathway_linked_compound = str_split(unlist(pathway[, linked_compound]), ";")[[1]]
      substrate_linked_compound = known_list_ORA[i, KEGG_ID]
      if (nchar(substrate_linked_compound) == 0){
        known_list_ORA[i, Status := "No Substrate Found"]
      }else{
        substrate_linked_compound = str_split(substrate_linked_compound, "; ")[[1]]
        known_list_ORA[i, Status := length(intersect(pathway_linked_compound, substrate_linked_compound))]
      }
    }
  }
  return(known_list_ORA)
}

plot_table = data.table(Rank = seq(10, 180, 10),
                        Number = vector("numeric", length(seq(10, 180, 10))),
                        Mean = vector("numeric", length(seq(10, 180, 10))))
for(k in seq(10, 180, 10)){
  a = bulk.overRepresentationAnalysis.known(known_list_ORA = known_list_ORA, up.to.rank = k)
  n = a[Status != "No Pathway Enriched" & Status != "No Substrate Found" & Status != "0", .N]
  m = mean(as.numeric(a[Status != "No Pathway Enriched" & Status != "No Substrate Found" & Status != "0", Status]))
  plot_table[Rank == k, Number := n]
  plot_table[Rank == k, Mean := m]
  print(k)
}

### plot ###
plot(x = plot_table[, Rank], y = plot_table[, Number])
ggplot(plot_table, aes(x = Rank, y = Number)) +
  geom_point() +
  theme(
    legend.position="none", plot.title = element_text(size=11),
    panel.grid.major = element_blank(), panel.grid.minor = element_blank(),
    panel.background = element_blank(), axis.line = element_line(colour = "black"))

## perform ORA on orphan list ##
orphan_list_ORA = data.table(SYMBOL = orphan_list[, unique(HGNC_symbol)],
                             Enriched_pathway = vector("character", length(orphan_list[, unique(HGNC_symbol)])))
orphan_list_ORA_result = vector("list", nrow(orphan_list_ORA))
names(orphan_list_ORA_result) = orphan_list_ORA[, SYMBOL]
for (i in 1:nrow(orphan_list_ORA)){
  etcs_atomic = etcs[SYMBOL == orphan_list_ORA[i, SYMBOL]][order(Score, A, decreasing = TRUE)][1:20]
  etcs_atomic_feature = 
    lapply(etcs_atomic[,na.omit(Formula)], function(x){as.character(ConvertNames(x, "Formula")[,"KEGG ID"])}) |>
    unlist() |>
    str_split(";") |>
    unlist() |>
    str_trim()
  
  ora_reporting = overRepresentationAnalysis(features = unique(etcs_atomic_feature), 
                                             funCatList = human_metabolic_pathways_list, 
                                             universe = univ)
  ora_reporting = inner_join(x = data.table(ora_reporting)[pAdj <= 0.05][order(pAdj)][,pValue := NULL], 
                             y = human_metabolic_pathways[, .(functionalCategory = `Pathway id`, `Pathway name`, Class, compound_no)], 
                             by = "functionalCategory")
  setcolorder(ora_reporting, c("functionalCategory", "Pathway name", "Class", "pAdj", "compound_no", "nOverlap", "overlap"))
  
  orphan_list_ORA_result[[i]] = ora_reporting
  print(i)
}

for (i in 1:length(orphan_list_ORA_result)){
  pathway = inner_join(x = data.table("Pathway id" = orphan_list_ORA_result[[i]][,functionalCategory]),
                       y = human_metabolic_pathways[, .(`Pathway id`, linked_compound)],
                       by = "Pathway id")
  if (nrow(pathway) == 0){
    orphan_list_ORA[i, Enriched_pathway := "No Pathway Enriched"]
  }else{
    ep = paste(unlist(pathway[, `Pathway id`]), collapse = ";")
    orphan_list_ORA[i, Enriched_pathway := ep]
  }
}
orphan_list_ORA = as.data.table(separate_longer_delim(orphan_list_ORA, Enriched_pathway, delim = ";"))
orphan_list_ORA = data.table(orphan_list_ORA,
                             Linked_Compound = vector("character", nrow(orphan_list_ORA)))
for (i in 1:nrow(orphan_list_ORA)){
  if(orphan_list_ORA[i, Enriched_pathway] != 	"No Pathway Enriched"){
    ep = orphan_list_ORA[i, Enriched_pathway]
    lp = human_metabolic_pathways[`Pathway id` == ep, linked_compound]
    orphan_list_ORA[i, Linked_Compound := lp]
  }
}

##### 4 Orphan prediction #####
orphan_prediction = data.table(SYMBOL = rep(orphan_list[, HGNC_symbol], each = 20),
                               PredictionRank = rep(1:20, nrow(orphan_list)),
                               Prediction = vector("character", nrow(orphan_list)*20),
                               A = vector("numeric", nrow(orphan_list)*20),
                               B = vector("numeric", nrow(orphan_list)*20),
                               C = vector("numeric", nrow(orphan_list)*20),
                               Score = vector("numeric", nrow(orphan_list)*20),
                               Pathway_enriched = vector("character", nrow(orphan_list)*20))
for(o in 1:nrow(orphan_list)){
  orphan_gene = orphan_list[o, HGNC_symbol]
  etcs_atomic = etcs[SYMBOL == orphan_gene][order(Score, A, decreasing = TRUE)][1:20]
  etcs_atomic_feature = 
    lapply(etcs_atomic[,na.omit(Formula)], function(x){as.character(ConvertNames(x, "Formula")[,"KEGG ID"])}) |>
    unlist() |>
    str_split(";") |>
    unlist() |>
    str_trim()
  orphan_prediction[SYMBOL == orphan_gene, c("Prediction", "A", "B", "C", "Score") := etcs_atomic[, c("Formula", "A", "B", "C", "Score")]]
  ep = paste(orphan_list_ORA[SYMBOL == orphan_gene, Enriched_pathway], collapse = ";")
  if (ep != "No Pathway Enriched"){
    orphan_list_ORA_atomic = orphan_list_ORA[SYMBOL == orphan_gene]
    ep_status = vector("list", nrow(etcs_atomic))
    for (j in 1:nrow(etcs_atomic)){
      kegg_to_test = str_trim(unlist(str_split(ConvertNames(etcs_atomic[j, Formula], "Formula")[,"KEGG ID"], ";")))
      if (check_NA(kegg_to_test)){
        ep_status[[j]] = ""
      }else{
        path_loc = c()
        for (k in 1:length(kegg_to_test)){
          path_loc = c(grep(kegg_to_test[k], orphan_list_ORA_atomic[, Linked_Compound]), path_loc)
        }
        if (length(path_loc) == 0){
          ep_status[[j]] = ""
        }else{
          ep_status[[j]] = paste("Enriched in pathway", paste(unique(orphan_list_ORA_atomic[path_loc, Enriched_pathway]), collapse = ", "))
        }
      }
    }
    orphan_prediction[SYMBOL == orphan_gene, Pathway_enriched := unlist(ep_status)]
  }
  print(o)
}

fwrite(orphan_prediction, "03 Results/05 Building ConfidenceScore/Orphan Prediction Result.csv")
```
