## Supplementary material for "Predicting substrates for orphan Solute Carrier Proteins using multi-omics datasets": Code for generating results: Step7_De-comfounding.R

```
library(data.table)
library(tidyverse)

orphan_prediction = fread("03 Results/05 Building ConfidenceScore/Orphan Prediction Result.csv")
load("~/Desktop/SLC ver3/03 Results/05 Building ConfidenceScore/Building ConfidenceScore ABC (clean start with generated random).RData")

a = table(str_trim(unlist(str_split(na.omit(unlist(tb[, `KEGG ID`])), ";"))))


translate(getAM("C20H32N6O12S2 (glutathione oxi)")[which(names(getAM("C20H32N6O12S2 (glutathione oxi)")) %in% c(1,2))])

et_deconfounding = et[Formula == "C4H7NO4 (aspartate)"]

ConfidenceScoreC = function(ki.row){
  # Factor A quantile
  ns_dist1 = na.omit(etki$NCI60) # NCI60 #
  ns_dist2 = na.omit(etki$CCLE2019) # CCLE2019 #
  ns_dist3 = na.omit(etki$CCL180) # CCL180 #
  q1 = quantile(ns_dist1[ns_dist1 > 0.0], seq(0, 1, by = 0.1))
  q2 = quantile(ns_dist2[ns_dist2 > 0.0], seq(0, 1, by = 0.1))
  q3 = quantile(ns_dist3[ns_dist3 > 0.8], seq(0, 1, by = 0.1))
  # Factor B quantile
  ns_dist4 = -log10(na.omit(etki$CRISPRDEP))
  q4 = quantile(ns_dist4[ns_dist4 > -log10(0.077)], seq(0, 1, by = 0.1))
  data_info = data.table(data_name = c("NCI60", "CCLE2019", "CCL180"),
                         data_dist = c("q1", "q2", "q3"),
                         Factor_A_threshold = c(0.0, 0.0, 0.8),
                         Factor_C_threshold = c(0.0, 0.0, 0.0))
  temp = getAM(ki.row$Formula)
  if (check_NA(temp)){
    Factor_C = 0
  }else if (!1 %in% names(temp) & !2 %in% names(temp)){
    Factor_C = 0
  }else{
    proximal = translate(temp[which(names(temp) %in% c(1,2))])
    if (check_NA(proximal)){
      Factor_C = 0
    }else{
      proximal_ns = getNR(ki.row[, SYMBOL], proximal)
      if (nrow(proximal_ns) != 0) {
        Factor_C = 0
        for (l in 1:nrow(data_info)){
          N = unique(proximal_ns[, get(data_info[l, data_name])])
          for (n in N){
            if (is.na(n) | n < data_info[l, Factor_C_threshold]){
              Factor_C = Factor_C + 0
            }else if (n > data_info[l, Factor_C_threshold] & n < data_info[l, Factor_A_threshold]){
              Factor_C = Factor_C + 0.9/2
            }else if (n > data_info[l, Factor_A_threshold]){
              Factor_C = Factor_C + findInterval(n, get(data_info[l, data_dist]))*0.9
            }
          }
        }
      }else{
        Factor_C = 0
      }
    }
  }
  return(Factor_C)
}

store = c()
for (i in 1:nrow(et_deconfounding)){
  store = c(store, ConfidenceScoreC(et_deconfounding[i,]))
  print(i)
}
which(store == max(store))


# Factor A quantile
ns_dist1 = na.omit(etki$NCI60) # NCI60 #
ns_dist2 = na.omit(etki$CCLE2019) # CCLE2019 #
ns_dist3 = na.omit(etki$CCL180) # CCL180 #
q1 = quantile(ns_dist1[ns_dist1 > 0.0], seq(0, 1, by = 0.1))
q2 = quantile(ns_dist2[ns_dist2 > 0.0], seq(0, 1, by = 0.1))
q3 = quantile(ns_dist3[ns_dist3 > 0.8], seq(0, 1, by = 0.1))
# Factor B quantile
ns_dist4 = -log10(na.omit(etki$CRISPRDEP))
q4 = quantile(ns_dist4[ns_dist4 > -log10(0.077)], seq(0, 1, by = 0.1))
data_info = data.table(data_name = c("NCI60", "CCLE2019", "CCL180"),
                       data_dist = c("q1", "q2", "q3"),
                       Factor_A_threshold = c(0.0, 0.0, 0.8),
                       Factor_C_threshold = c(0.0, 0.0, 0.0))
temp = getAM(ki.row$Formula)
proximal = translate(unique(temp[which(names(temp) %in% c(1,2))]))
proximal_ns = getNR(ki.row[, SYMBOL], proximal)
for (k in 1:nrow(proximal_ns)) {
  for (l in 1:nrow(data_info)){
    n = proximal_ns[k, get(data_info[l, data_name])]
    if (is.na(n) | n < data_info[l, Factor_C_threshold]){
      print("+0")
    }else if (n > data_info[l, Factor_C_threshold] & n < data_info[l, Factor_A_threshold]){
      print("+0.45")
    }else if (n > data_info[l, Factor_A_threshold]){
      print(paste("+", findInterval(n, get(data_info[l, data_dist]))*0.9, sep = ""))
    }
  }
}

for (l in 1:nrow(data_info)){
  N = unique(proximal_ns[, get(data_info[l, data_name])])
  for (n in N){
    if (is.na(n) | n < data_info[l, Factor_C_threshold]){
      print("+0")
    }else if (n > data_info[l, Factor_C_threshold] & n < data_info[l, Factor_A_threshold]){
      print("+0.45")
    }else if (n > data_info[l, Factor_A_threshold]){
      print(paste("+", findInterval(n, get(data_info[l, data_dist]))*0.9, sep = ""))
    }
  }
}
```
