## Supplementary material for "Predicting substrates for orphan Solute Carrier Proteins using multi-omics datasets": Code for generating results: 01 Generate SLC-drug interaction table.R

```
##### 0 load library #####
library(data.table)
library(tidyverse)
library(progress)

##### 1 read files #####
SLC_transcriptomics = fread("03 Results/01 Correlation Analysis/CCLE2019/CCLE2019 SLC normalised transcript counts.csv") # DepMap Public 23Q2 Omics
CCLE_metadata = read.delim("01 Raw Data/01 Correlation Analysis/01 CCLE2019/Cell_lines_annotations_20181226.txt")
Cell_line_metadata = fread("01 Raw Data/06 drug sensitivity/secondary-screen-cell-line-info.csv")
Drug_treatment_metadata = fread("01 Raw Data/06 drug sensitivity/secondary-screen-replicate-collapsed-treatment-info.csv")
Drug_screen = fread("01 Raw Data/06 drug sensitivity/secondary-screen-replicate-collapsed-logfold-change.csv")

##### 2 Primary treatment of file #####

# remove cell lines failed STR profiling #
Drug_screen = Drug_screen[-grep("FAILED_STR", V1),]

# translate the CCLE name to ACH code in SLC_transcriptomics #
CCLE_ach = unlist(lapply(colnames(SLC_transcriptomics)[2:ncol(SLC_transcriptomics)], function(x){CCLE_metadata$depMapID[which(CCLE_metadata$CCLE_ID == x)]}))
colnames(SLC_transcriptomics)[2:ncol(SLC_transcriptomics)] = CCLE_ach

# match cell lines in SLC_transcriptomics and drug_screen #
shared_cell_line = intersect(Drug_screen$V1, colnames(SLC_transcriptomics)[2:ncol(SLC_transcriptomics)])
SLC_transcriptomics = SLC_transcriptomics[, c(TRUE, colnames(SLC_transcriptomics)[2:ncol(SLC_transcriptomics)] %in% shared_cell_line), with = FALSE]
Drug_screen = Drug_screen[V1 %in% shared_cell_line, ]
setcolorder(SLC_transcriptomics, c("SYMBOL", Drug_screen$V1))

##### 3 Fit curve to SLC-drug pairs with loess #####
Curve = function(S, D){
  # get the metadata for desired drug, produce the screen profile for that drug #
  D_metadata = Drug_treatment_metadata[name == D]
  D_screen = Drug_screen[,c(TRUE, colnames(Drug_screen)[2:ncol(Drug_screen)] %in% D_metadata$column_name), with = FALSE]
  
  # get the expression profile of desired SLC, acquire cell lines of high and low expression #
  SLC_expression = as.numeric(unlist(SLC_transcriptomics[SYMBOL == S, 3:ncol(SLC_transcriptomics)]))
  # if the SLC is not found in the transcriptomics, terminate the function and return NA #
  if (length(SLC_expression) == 0){
    output = list(NaN, NaN, NaN)
    names(output) = c("Best.log.Dose", "Minimum.Wilcox.p", "Absolute.Median.Difference")
    return(output)
  }
  q = quantile(SLC_expression, c(0.2, 0.8))
  # if the lower quantile is not 0 #
  if (q[1] != 0){
    low_expression_cell = colnames(SLC_transcriptomics)[which(SLC_expression < q[1]) + 2]
    # if the lower quantile is 0 #
  }else{
    low_expression_cell = colnames(SLC_transcriptomics)[which(SLC_expression == q[1]) + 2]
  }
  high_expression_cell = colnames(SLC_transcriptomics)[which(SLC_expression > q[2]) + 2]
  
  # pool dose-response curve data for both high and low expression #
  cell_viability_for_D = melt(D_screen, id.vars = "V1")
  colnames(cell_viability_for_D) = c("cell", "dose", "viability")
  cell_viability_for_D$dose = as.numeric(str_match(cell_viability_for_D$dose, "::\\s*(.*?)\\s*::")[,2])
  cell_viability_for_D$log.dose = round(log10(as.numeric(cell_viability_for_D$dose)), 2)
  cell_viability_for_D$expression_type = vector("character", nrow(cell_viability_for_D))
  
  for (i in 1:nrow(cell_viability_for_D)){
    if (cell_viability_for_D$cell[i] %in% low_expression_cell){
      cell_viability_for_D[i, expression_type := "low expression"] # low expression
    }else if (cell_viability_for_D$cell[i] %in% high_expression_cell){
      cell_viability_for_D[i, expression_type := "high expression"] # high expression
    }else{
      cell_viability_for_D[i, expression_type := "NE"] # not examined #
    }
  }
  
  cell_viability_for_D = cell_viability_for_D[expression_type != "NE"][order(expression_type)]
  fit_high = loess(viability ~ log.dose, data = cell_viability_for_D[expression_type == "high expression"])
  fit_low = loess(viability ~ log.dose, data = cell_viability_for_D[expression_type == "low expression"])
  new_dose = expand.grid(log.dose = seq(-3.21, 1, 0.01))
  preds_high = predict(fit_high, newdata = new_dose)
  preds_low = predict(fit_low, newdata = new_dose)
  t_statistic = t.test(preds_high, preds_low)
  output = list(p_value = t_statistic$p.value,
                md = as.numeric(t_statistic$estimate[1] - t_statistic$estimate[2]))
  return(output)
}

### use examples ###
Curve("SLC26A2", "bis(maltolato)oxovanadium(IV)")
Curve("SLC35F2", "YM-155")
Curve("SLCO2B1", "atorvastatin")
Curve("SLC19A2", "thiamine")
Curve("PTCHD4", "idasanutlin")

##### 4 All interaction calculation #####
SLC_list = SLC_transcriptomics$SYMBOL
Compound_list = unique(Drug_treatment_metadata$name)

# SLC-Compound Interaction (SCI)
SCI = data.table(SLC = rep(SLC_list, each = length(Compound_list)),
                 Compound = rep(Compound_list, length(SLC_list)),
                 pvalue = vector("numeric", 747168),
                 mean.diff = vector("numeric", 747168))
for (i in 1:nrow(SCI)){
  ans = Curve(SCI[i, SLC], SCI[i, Compound])
  SCI[i, pvalue := ans$p_value]
  SCI[i, mean.diff := ans$md]
  print(i)
}

fwrite(SCI, "03 Results/06 Drug sensitivity/SCI.csv")
```
