## Supplementary material for "Predicting substrates for orphan Solute Carrier Proteins using multi-omics datasets": Code for generating results: 02 Pseudo-drug benchmark.R

```
library(data.table)
library(tidyverse)
library(hrbrthemes)
library(viridis)

##### 1 read files #####
SCI = fread("03 Results/06 Drug sensitivity/SCI.csv")
load("~/Desktop/SLC ver3/03 Results/06 Drug sensitivity/Using curve function.RData")

#### 2 Benchmarking ####
# Known drug target #
kdt = fread("01 Raw Data/00 SLC annotation 2024/known interaction/Known Drug Target.csv")

# eliminate drug compound only in primary screen #
kdt = kdt[Name %in% Drug_treatment_metadata$name]

# build kdt Curve data table #
kdt_curve = kdt
kdt_curve$pvalue = vector("numeric", nrow(kdt_curve)) # p-value of paired t-test
kdt_curve$mean.diff = vector("numeric", nrow(kdt_curve)) # absolute mean difference by paired t-test

for (i in 1:nrow(kdt_curve)){
  gene = kdt_curve[i, SYMBOL]
  drug = kdt_curve[i, Name]
  kdt_curve[i, pvalue := SCI[SLC == gene & Compound == drug, pvalue]]
  kdt_curve[i, mean.diff := SCI[SLC == gene & Compound == drug, mean.diff]]
  print(i)
}

# build Curve profile for simulated drug target #
fake_kdt_curve_collection = vector("list", 100)
kdt_curve_gene_pool = unique(kdt_curve$SYMBOL)
kdt_curve_drug_pool = unique(kdt_curve$Name)

for (sample_time in 1:100){
  # initialise fake kdt_curve data table #
  fake_kdt_curve = setNames(data.table(matrix(ncol = ncol(kdt_curve), nrow = nrow(kdt_curve))), colnames(kdt_curve))
  char.col = colnames(fake_kdt_curve)[1:5]
  num.col = colnames(fake_kdt_curve)[6:7]
  fake_kdt_curve[, (char.col) := lapply(.SD, as.character), .SDcols = char.col]
  fake_kdt_curve[, (num.col) := lapply(.SD, as.character), .SDcols = num.col]
  fake_kdt_curve[, Action := rep("Fake", nrow(fake_kdt_curve))]
  fake_kdt_curve[, c("SYMBOL", "Localisation", "Compartment") := as.list(kdt_curve[, c("SYMBOL", "Localisation", "Compartment")])]
  # for each SLC, shuffle fake substrate formula #
  for (j in 1:nrow(fake_kdt_curve)){
    true_drug = kdt_curve[SYMBOL == fake_kdt_curve$SYMBOL[j]][, Name]
    fake_drug_pool = kdt_curve_drug_pool[!kdt_curve_drug_pool %in% true_drug]
    fake_drug = sample(fake_drug_pool, 1)
    fake_kdt_curve[j, Name := fake_drug]
    fake_kdt_curve[j, c("pvalue", "mean.diff") := 
                     SCI[SLC == fake_kdt_curve$SYMBOL[j] & Compound == fake_drug, c("pvalue", "mean.diff")]]
  }
  fake_kdt_curve_collection[[sample_time]] = fake_kdt_curve
  print(paste(sample_time, "done"))
}

#### 3 Benchmark reporting ####
## absolute mean difference ##
fkdt_mean = unlist(lapply(1:100, function(x){fake_kdt_curve_collection[[x]][, mean(abs(as.numeric(mean.diff)))]}))
kdt_mean = kdt_curve[,mean(abs(as.numeric(mean.diff)))]
boxplot(fkdt_mean)
abline(h = kdt_mean, col = "blue")

length(which(fkdt_mean > kdt_mean))

## mean log p-value ##
fkdt_pmean = unlist(lapply(1:100, function(x){fake_kdt_curve_collection[[x]][, mean(-log10(as.numeric(pvalue)))]}))
kdt_pmean = kdt_curve[,mean(-log10(as.numeric(pvalue)))]
boxplot(fkdt_pmean)
abline(h = kdt_pmean, col = "blue")

benchmark = data.table(mean.diff = c(fkdt_mean, kdt_mean),
                   log.p = c(fkdt_pmean, kdt_pmean),
                   Status = c(rep("random", 100), "true"))
fwrite(benchmark, "03 Results/06 Drug sensitivity/t-test model prediction benchmark.csv")

ggplot() + # fill=name allow to automatically dedicate a color for each group
  geom_boxplot(data = benchmark[Status == "random"], aes(x = Status, y = log.p)) +
  scale_fill_viridis(discrete = TRUE, alpha=0.6, option="C") +
  theme_ipsum() +
  theme(
    legend.position="none",
    plot.title = element_text(size=11)
  ) +
  geom_hline(aes(yintercept=benchmark[Status == "true", log.p]), colour="#990000", linetype="dashed") +
  theme(panel.grid.major = element_blank(), panel.grid.minor = element_blank(),
        panel.background = element_blank(), axis.line = element_line(colour = "black"))
```
