## Supplementary material for "Predicting substrates for orphan Solute Carrier Proteins using multi-omics datasets": Code for generating results: 03 Check dose effect upon SLC-drug interaction.R

```
##### 0 load library #####
library(data.table)
library(tidyverse)
library(hrbrthemes)
library(viridis)
load("~/Desktop/SLC ver3/03 Results/06 Drug sensitivity/SCI.RData")

##### 1 Build new SLC-drug interaction table based on interaction results #####
kdt <- fread("01 Raw Data/00 SLC annotation 2024/known interaction/Known Drug Target.csv")[Name %in% Drug_treatment_metadata$name]

##### 2 Append dose effect to known drug table #####
CheckDoseEffect = function(S, D, original.data = FALSE){
  # get the metadata for desired drug, produce the screen profile for that drug #
  D_metadata = Drug_treatment_metadata[name == D]
  D_screen = Drug_screen[,c(TRUE, colnames(Drug_screen)[2:ncol(Drug_screen)] %in% D_metadata$column_name), with = FALSE]
  
  # get the expression profile of desired SLC, acquire cell lines of high and low expression #
  SLC_expression = as.numeric(unlist(SLC_transcriptomics[SYMBOL == S, 3:ncol(SLC_transcriptomics)]))
  # if the SLC is not found in the transcriptomics, terminate the function and return NA #
  if (length(SLC_expression) == 0){
    output = list(NaN, NaN)
    names(output) = c("p_value", "md")
    return(output)
  }
  q = quantile(SLC_expression, c(0.2, 0.8))
  # if the lower quantile is not 0 #
  if (q[1] != 0){
    low_expression_cell = colnames(SLC_transcriptomics)[which(SLC_expression < q[1]) + 2]
    # if the lower quantile is 0 #
  }else{
    low_expression_cell = colnames(SLC_transcriptomics)[which(SLC_expression == q[1]) + 2]
  }
  high_expression_cell = colnames(SLC_transcriptomics)[which(SLC_expression > q[2]) + 2]
  
  # pool dose-response curve data for both high and low expression #
  cell_viability_for_D = melt(D_screen, id.vars = "V1")
  colnames(cell_viability_for_D) = c("cell", "dose", "viability")
  cell_viability_for_D$dose = as.numeric(str_match(cell_viability_for_D$dose, "::\\s*(.*?)\\s*::")[,2])
  cell_viability_for_D$log.dose = round(log10(as.numeric(cell_viability_for_D$dose)), 2)
  cell_viability_for_D$expression_type = vector("character", nrow(cell_viability_for_D))
  
  for (i in 1:nrow(cell_viability_for_D)){
    if (cell_viability_for_D$cell[i] %in% low_expression_cell){
      cell_viability_for_D[i, expression_type := "low expression"] # low expression
    }else if (cell_viability_for_D$cell[i] %in% high_expression_cell){
      cell_viability_for_D[i, expression_type := "high expression"] # high expression
    }else{
      cell_viability_for_D[i, expression_type := "NE"] # not examined #
    }
  }
  
  cell_viability_for_D = cell_viability_for_D[expression_type != "NE"][order(expression_type)]
  fit_high = loess(viability ~ log.dose, data = cell_viability_for_D[expression_type == "high expression"])
  fit_low = loess(viability ~ log.dose, data = cell_viability_for_D[expression_type == "low expression"])
  if (isTRUE(original.data)){
    original.dose = c(-3.214, -2.612, -2.010, -1.408, -0.806, -0.204, 0.398, 1.000)
    new_dose = expand.grid(log.dose = original.dose)
    preds_high = predict(fit_high, newdata = new_dose)
    preds_low = predict(fit_low, newdata = new_dose)
    CorIn = cor.test(original.dose, abs(as.numeric(preds_high - preds_low)), method = "spearman")
  }else{
    new_dose = expand.grid(log.dose = seq(-3.21, 1, 0.01))
    preds_high = predict(fit_high, newdata = new_dose)
    preds_low = predict(fit_low, newdata = new_dose)
    CorIn = cor.test(seq(-3.21, 1, 0.01), abs(as.numeric(preds_high - preds_low)), method = "spearman")
  }
  output = list(p_value = as.numeric(CorIn$p.value),
                rho = as.numeric(CorIn$estimate))
  return(output)
}
kdt_CDE = kdt
kdt_CDE$pvalue = vector("numeric", nrow(kdt_CDE)) # p-value of paired t-test
kdt_CDE$rho = vector("numeric", nrow(kdt_CDE)) # absolute mean difference by paired t-test

for (i in 1:nrow(kdt_CDE)){
  gene = kdt_CDE[i, SYMBOL]
  drug = kdt_CDE[i, Name]
 #statistics = CheckDoseEffect(gene, drug, original.data = TRUE)
  statistics = CheckDoseEffect(gene, drug)
  kdt_CDE[i, pvalue := statistics$p_value]
  kdt_CDE[i, rho := statistics$rho]
  print(i)
}

# build CDE profile for simulated drug target #
fake_kdt_CDE_collection = vector("list", 100)
kdt_CDE_gene_pool = unique(kdt_CDE$SYMBOL)
kdt_CDE_drug_pool = unique(kdt_CDE$Name)

for (sample_time in 1:100){
  # initialise fake kdt_CDE data table #
  fake_kdt_CDE = setNames(data.table(matrix(ncol = ncol(kdt_CDE), nrow = nrow(kdt_CDE))), colnames(kdt_CDE))
  char.col = colnames(fake_kdt_CDE)[1:5]
  num.col = colnames(fake_kdt_CDE)[6:7]
  fake_kdt_CDE[, (char.col) := lapply(.SD, as.character), .SDcols = char.col]
  fake_kdt_CDE[, (num.col) := lapply(.SD, as.character), .SDcols = num.col]
  fake_kdt_CDE[, c("SYMBOL", "Localisation", "Compartment") := as.list(kdt_CDE[, c("SYMBOL", "Localisation", "Compartment")])]
  # for each SLC, shuffle fake substrate formula #
  for (j in 1:nrow(fake_kdt_CDE)){
    true_drug = kdt_CDE[SYMBOL == fake_kdt_CDE$SYMBOL[j]][, Name]
    fake_drug_pool = kdt_CDE_drug_pool[!kdt_CDE_drug_pool %in% true_drug]
    fake_drug = sample(fake_drug_pool, 1)
    fake_kdt_CDE[j, Name := fake_drug]
   #statistics = CheckDoseEffect(fake_kdt_CDE[j, SYMBOL], fake_drug, original.data = TRUE)
    statistics = CheckDoseEffect(fake_kdt_CDE[j, SYMBOL], fake_drug)
    fake_kdt_CDE[j, c("pvalue", "rho") := list(statistics$p_value, statistics$rho)]
    print(paste(sample_time, ":", round(j/nrow(fake_kdt_CDE),2)))
  }
  fake_kdt_CDE_collection[[sample_time]] = fake_kdt_CDE
  print(paste(sample_time, "done"))
}


##### 3 Benchmark reporting #####
## rho value reporting ##
fkdt_rho_mean = unlist(lapply(1:100, function(x){fake_kdt_CDE_collection[[x]][, mean(abs(as.numeric(rho)))]}))
kdt_rho_mean = kdt_CDE[,mean(abs(as.numeric(rho)))]
boxplot(fkdt_rho_mean, ylim = c(0, 0.7))
abline(h = kdt_rho_mean, col = "blue")

## mean p-value ##
fkdt_p_mean = unlist(lapply(1:100, function(x){fake_kdt_CDE_collection[[x]][, mean(as.numeric(pvalue))]}))
kdt_p_mean = kdt_CDE[,mean(as.numeric(pvalue))]
boxplot(fkdt_p_mean)
abline(h = kdt_p_mean, col = "blue")

## examples ##
plotCurve("SLC2A1", "oligomycin-a")
CheckDoseEffect("SLC2A1", "oligomycin-a")
CheckDoseEffect("SLC2A1", "oligomycin-a", original.data = TRUE)

plotCurve("SLC35F2", "YM-155")
CheckDoseEffect("SLC35F2", "YM-155")
CheckDoseEffect("SLC35F2", "YM-155", original.data = TRUE)

## produce benchmark plots
benchmark = data.table(rho.mean = c(fkdt_rho_mean, kdt_rho_mean),
                       log.p.mean = c(-log10(fkdt_p_mean), -log10(kdt_p_mean)),
                       Status = c(rep("random", 100), "true"))
fwrite(benchmark, "03 Results/06 Drug sensitivity/check dose effect benchmark.csv")

ggplot() + # fill=name allow to automatically dedicate a color for each group
  geom_boxplot(data = benchmark[Status == "random"], aes(x = Status, y = log.p.mean)) +
  scale_fill_viridis(discrete = TRUE, alpha=0.6, option="C") +
  theme_ipsum() +
  theme(
    legend.position="none",
    plot.title = element_text(size=11)
  ) +
  geom_hline(aes(yintercept=benchmark[Status == "true", log.p.mean]), colour="#990000", linetype="dashed") +
  theme(panel.grid.major = element_blank(), panel.grid.minor = element_blank(),
        panel.background = element_blank(), axis.line = element_line(colour = "black"))


##### 4 select the filter threshold for benchmark #####
## find the best threshold ##
p_cutoff = seq(0.05, 0, -0.001)
rho_cutoff = seq(0, 1, 0.01)
sweep = data.table(alpha = rep(p_cutoff, each = length(rho_cutoff)),
                   rho = rep(rho_cutoff, length(p_cutoff)),
                   discovery = vector("numeric", length(p_cutoff)*length(rho_cutoff)))
for(i in 1:nrow(sweep)){
  kdt_pnum = kdt_CDE[pvalue < sweep$alpha[i] & abs(as.numeric(rho)) > sweep$rho[i], .N]/kdt_CDE[, .N]
  fkdt_pnum = mean(unlist(lapply(1:100, function(x){fake_kdt_CDE_collection[[x]][pvalue < sweep$alpha[i] & abs(as.numeric(rho)) > sweep$rho[i], .N]/fake_kdt_CDE_collection[[x]][, .N]})))
  sweep[i, discovery := kdt_pnum - fkdt_pnum]
  print(i)
}

## when predicted data points is 8, best discovery is 0.3061224, best p is 0.05, best absolute rho is 0.05
## when predicted data points is 421, best discovery is 0.9591837, best p is 0.05, best absolute rho is 0.05


## calculate for every interaction ##
SCI$dose.pvalue = vector("numeric", nrow(SCI)) # p-value of paired t-test
SCI$dose.rho = vector("numeric", nrow(SCI)) # absolute mean difference by paired t-test

for (i in 1:nrow(SCI)){
  gene = SCI[i, SLC]
  drug = SCI[i, Compound]
  statistics = CheckDoseEffect(gene, drug)
  SCI[i, dose.pvalue := statistics$p_value]
  SCI[i, dose.rho := statistics$rho]
  print(i)
}

fwrite(SCI, "03 Results/06 Drug sensitivity/SCI_dose_correlation.csv")
```
