## Supplementary material for "Predicting substrates for orphan Solute Carrier Proteins using multi-omics datasets": Code for generating results: 04 Pseudo-SLC benchmark.R

```
##### 0 load library #####
library(data.table)
library(tidyverse)
library(progress)

##### 1 read files #####
SLC_transcriptomics = fread("03 Results/01 Correlation Analysis/CCLE2019/CCLE2019 SLC normalised transcript counts.csv") # DepMap Public 23Q2 Omics
CCLE_metadata = read.delim("01 Raw Data/01 Correlation Analysis/01 CCLE2019/Cell_lines_annotations_20181226.txt")
Cell_line_metadata = fread("01 Raw Data/06 drug sensitivity/secondary-screen-cell-line-info.csv")
Drug_treatment_metadata = fread("01 Raw Data/06 drug sensitivity/secondary-screen-replicate-collapsed-treatment-info.csv")
Drug_screen = fread("01 Raw Data/06 drug sensitivity/secondary-screen-replicate-collapsed-logfold-change.csv")

##### 2 Primary treatment of file #####

# remove cell lines failed STR profiling #
Drug_screen = Drug_screen[-grep("FAILED_STR", V1),]

# translate the CCLE name to ACH code in SLC_transcriptomics #
CCLE_ach = unlist(lapply(colnames(SLC_transcriptomics)[2:ncol(SLC_transcriptomics)], function(x){CCLE_metadata$depMapID[which(CCLE_metadata$CCLE_ID == x)]}))
colnames(SLC_transcriptomics)[2:ncol(SLC_transcriptomics)] = CCLE_ach

# match cell lines in SLC_transcriptomics and drug_screen #
shared_cell_line = intersect(Drug_screen$V1, colnames(SLC_transcriptomics)[2:ncol(SLC_transcriptomics)])
SLC_transcriptomics = SLC_transcriptomics[, c(TRUE, colnames(SLC_transcriptomics)[2:ncol(SLC_transcriptomics)] %in% shared_cell_line), with = FALSE]
Drug_screen = Drug_screen[V1 %in% shared_cell_line, ]
setcolorder(SLC_transcriptomics, c("SYMBOL", Drug_screen$V1))

##### 3 Fit curve to SLC-drug pairs with loess #####
Curve = function(D){
  # get the metadata for desired drug, produce the screen profile for that drug #
  D_metadata = Drug_treatment_metadata[name == D]
  D_screen = Drug_screen[,c(TRUE, colnames(Drug_screen)[2:ncol(Drug_screen)] %in% D_metadata$column_name), with = FALSE]
  
  # get the pseudo-expression profile of desired SLC, acquire cell lines of high and low expression #
  cell_pool = colnames(SLC_transcriptomics)
  cell_pool_index = 1:length(cell_pool)
  low_index = sample(cell_pool_index, 0.2*length(cell_pool))
  high_index = sample(cell_pool_index[!cell_pool_index %in% low_index], 0.2*length(cell_pool))
  low_expression_cell = colnames(SLC_transcriptomics)[low_index]
  high_expression_cell = colnames(SLC_transcriptomics)[high_index]
  
  # pool dose-response curve data for both high and low expression #
  cell_viability_for_D = melt(D_screen, id.vars = "V1")
  colnames(cell_viability_for_D) = c("cell", "dose", "viability")
  cell_viability_for_D$dose = as.numeric(str_match(cell_viability_for_D$dose, "::\\s*(.*?)\\s*::")[,2])
  cell_viability_for_D$log.dose = round(log10(as.numeric(cell_viability_for_D$dose)), 2)
  cell_viability_for_D$expression_type = vector("character", nrow(cell_viability_for_D))
  
  for (i in 1:nrow(cell_viability_for_D)){
    if (cell_viability_for_D$cell[i] %in% low_expression_cell){
      cell_viability_for_D[i, expression_type := "low expression"] # low expression
    }else if (cell_viability_for_D$cell[i] %in% high_expression_cell){
      cell_viability_for_D[i, expression_type := "high expression"] # high expression
    }else{
      cell_viability_for_D[i, expression_type := "NE"] # not examined #
    }
  }
  
  cell_viability_for_D = cell_viability_for_D[expression_type != "NE"][order(expression_type)]
  fit_high = loess(viability ~ log.dose, data = cell_viability_for_D[expression_type == "high expression"])
  fit_low = loess(viability ~ log.dose, data = cell_viability_for_D[expression_type == "low expression"])
  new_dose = expand.grid(log.dose = seq(-3.21, 1, 0.01))
  preds_high = predict(fit_high, newdata = new_dose)
  preds_low = predict(fit_low, newdata = new_dose)
  t_statistic = t.test(preds_high, preds_low)
  output = list(p_value = t_statistic$p.value,
                md = as.numeric(t_statistic$estimate[1] - t_statistic$estimate[2]))
  return(output)
}

drug_list = unique(Drug_treatment_metadata[, name])
pseudoSLC_benchmark = vector("list", length(drug_list))
names(pseudoSLC_benchmark) = drug_list
for (drug_index in 1:length(drug_list)){
  drug = drug_list[drug_index]
  benchmarking = vector("list", 100)
  for (sample_time in 1:100){
    benchmarking[[sample_time]] = Curve(drug)
    print(paste(drug, ":", sample_time))
  }
  pseudoSLC_benchmark[[drug_index]] = benchmarking
}

pseudo_benchmark = data.table(Drug = rep(drug_list, each = 100),
                              logp = vector("numeric", length(drug_list)*100),
                              amd = vector("numeric", length(drug_list)*100))
for (drug_index in 1:length(drug_list)){
  p = unlist(lapply(1:100, function(x){pseudoSLC_benchmark[[drug_index]][[x]][[1]]}))
  md = unlist(lapply(1:100, function(x){pseudoSLC_benchmark[[drug_index]][[x]][[2]]}))
  pseudo_benchmark[Drug == drug_list[drug_index], logp := -log10(p)]
  pseudo_benchmark[Drug == drug_list[drug_index], amd := abs(md)]
}

boxplot(pseudo_benchmark[, mean(logp), by = Drug][, V1])
boxplot(pseudo_benchmark[, mean(amd), by = Drug][, V1])

pseudo_significance = data.table(Drug = drug_list,
                                 logp_threshold = vector("numeric", length(drug_list)),
                                 amd_threshold = vector("numeric", length(drug_list)))
for (drug_index in 1:length(drug_list)){
  pM = pseudo_benchmark[Drug == drug_list[drug_index], mean(logp)]
  pSD = 2*pseudo_benchmark[Drug == drug_list[drug_index], sd(logp)]
  pseudo_significance[Drug == drug_list[drug_index], logp_threshold := pM + pSD]
  
  amdM = pseudo_benchmark[Drug == drug_list[drug_index], mean(amd)]
  amdSD = 2*pseudo_benchmark[Drug == drug_list[drug_index], sd(amd)]
  pseudo_significance[Drug == drug_list[drug_index], amd_threshold := amdM + amdSD]
}

fwrite(pseudo_significance, "03 Results/06 Drug sensitivity/pseudoSLC benchmarked significance.csv")
```
