## Supplementary material for "Predicting substrates for orphan Solute Carrier Proteins using multi-omics datasets": Code for generating results: 05 Generate drug predictions.R

```
library(data.table)
library(tidyverse)

SCI = fread("03 Results/06 Drug sensitivity/SCI_dose_correlation.csv")
pseudoSLC_significance = fread("03 Results/06 Drug sensitivity/pseudoSLC benchmarked significance.csv")

# remove interactions that are likely to fall in pseudo significance #
SCI_filtered = setNames(data.table(matrix(ncol = ncol(SCI), nrow = 0)), colnames(SCI))
for (drug in pseudoSLC_significance[, Drug]){
  logpTh = pseudoSLC_significance[Drug == drug, logp_threshold]
  amdTh = pseudoSLC_significance[Drug == drug, amd_threshold]
  SCI_atomic = SCI[Compound == drug][-log10(pvalue) > logpTh & abs(mean.diff) > amdTh]
  SCI_filtered = rbind(SCI_filtered, SCI_atomic)
}

# taking into account of dose-viability correlation
SCI_filtered = SCI_filtered[dose.pvalue < 0.05][order(-abs(mean.diff), -abs(dose.rho))]

# remove mutation specific drugs
SCI_filtered = SCI_filtered[Compound != "poziotinib"]

# generate prediction
drug_prediction = SCI_filtered[1:50]
fwrite(drug_prediction, "03 Results/06 Drug sensitivity/Top predicted drug SLC interaction.csv")
fwrite(SCI_filtered, "03 Results/06 Drug sensitivity/Filtered predicted drug SLC interaction.csv")

# plotCurve
load("~/Desktop/SLC ver3/03 Results/06 Drug sensitivity/Using plotCurve function.RData")

for (i in 11:50){
  print(plotCurve(drug_prediction[i, SLC], drug_prediction[i, Compound]))
  print(paste(drug_prediction[i, SLC], drug_prediction[i, Compound]))
  Sys.sleep(5.0)
}


plotCurve(drug_prediction[i, SLC], drug_prediction[i, Compound])
```
