## Supplementary material for "Predicting substrates for orphan Solute Carrier Proteins using multi-omics datasets": Code for generating results: 06 Interaction with RITA.R

```
##### 0 libraries #####
library(data.table)
library(tidyverse)

##### 1 read files #####
setwd("03 Results/06 Drug sensitivity/Interaction with RITA/")
temp = list.files(pattern="*.csv")
sensitising = temp[!temp %in% "SLC3A1.csv"]
attenuating = "SLC3A1.csv"
SLCs = lapply(sensitising, fread) # read through all the rand.csv files

SLCs_high_expression = lapply(SLCs, function(x){unlist(x[expression_type == "high expression", cell, by = cell][,1])})
SLCs_low_expression = lapply(SLCs, function(x){unlist(x[expression_type == "low expression", cell, by = cell][,1])})

a = as.data.table(table(unlist(SLCs_high_expression)))
b = as.data.table(table(unlist(SLCs_low_expression)))

draw = function(g){
  ggplot(g, aes(x=log10(dose), y=viability, fill =expression_type, color = expression_type)) + 
    scale_fill_manual(values=c("#63BBB0", "#AC667E")) +
    scale_color_manual(values=c("#63BBB0", "#AC667E")) +
    geom_point(position = position_dodge(0.2), alpha = 0.5) + 
    geom_smooth(method = 'loess') + 
    scale_y_continuous(limits = c(-7.5, 2.5), breaks = seq(-7.5, 2.5, 2.5)) +
    theme(
      legend.position="none", plot.title = element_text(size=11),
      panel.grid.major = element_blank(), panel.grid.minor = element_blank(),
      panel.background = element_blank(), axis.line = element_line(colour = "black"))
}
glist = lapply(SLCs, draw)
for (i in 1:length(glist)){
  print(glist[[i]])
}

ggplot(a, aes(x = N)) +
  geom_histogram(fill = "#63BBB0", color="#e9ecef", binwidth = 1) +
  scale_x_continuous(breaks = seq(1,8,1)) +
  scale_y_continuous(limits = c(0, 150)) +
  theme(
    legend.position="none", plot.title = element_text(size=11),
    panel.grid.major = element_blank(), panel.grid.minor = element_blank(),
    panel.background = element_blank(), axis.line = element_line(colour = "black"))

ggplot(b, aes(x = N)) +
  geom_histogram(fill = "#AC667E", color="#e9ecef", binwidth = 1) +
  scale_x_continuous(breaks = seq(1,8,1)) +
  scale_y_continuous(limits = c(0, 150)) +
  theme(
    legend.position="none", plot.title = element_text(size=11),
    panel.grid.major = element_blank(), panel.grid.minor = element_blank(),
    panel.background = element_blank(), axis.line = element_line(colour = "black"))

CCLE_metadata = fread("01 Raw Data/02 CRISPR dependency/Model.csv")

colnames(a) = c("ModelID", "Number")
colnames(b) = c("ModelID", "Number")

a = inner_join(a[Number >= 5], CCLE_metadata)
b = inner_join(b[Number >= 5], CCLE_metadata)
```
