## Supplementary material for "Predicting substrates for orphan Solute Carrier Proteins using multi-omics datasets": Code for generating results: Girardi association match.R

```
library(data.table)
library(readxl)

association = as.data.table(read_excel("01 Raw Data/07 Literature match/41589_2020_483_MOESM3_ESM.xlsx"))
Drug_treatment_metadata = fread("01 Raw Data/06 drug sensitivity/secondary-screen-replicate-collapsed-treatment-info.csv")
prediction = fread("03 Results/06 Drug sensitivity/Filtered predicted drug SLC interaction.csv")

compound_name = c(rep("azacitidine",        6),
                  rep("mercaptopurine",     1),
                  rep("artesunate",         2),
                  rep("belinostat",         3),
                  rep(NA,                   3),
                  rep("chidamide",          3),
                  rep(NA,                   1),
                  rep("cisplatin",          8),
                  rep("clofarabine",        5),
                  rep("cytarabine",         9),
                  rep("decitabine",        10),
                  rep("digitoxin",          2),
                  rep("dihydroartemisinin", 7),
                  rep("disulfiram",         3),
                  rep("docetaxel",          9),
                  rep("dronedarone",        1),
                  rep("entinostat",         1),
                  rep("epirubicin",         4),
                  rep("etoposide",          4),
                  rep("gemcitabine",        2),
                  rep("homoharringtonine",  2),
                  rep("idarubicin",         1),
                  rep(NA,                   2),
                  rep("methotrexate",       3),
                  rep(NA,                   2),
                  rep("mitomycin-c",       14),
                  rep("mitoxantrone",       7),
                  rep("mycophenolic-acid",  5),
                  rep(NA,                   7),
                  rep("paclitaxel",         2),
                  rep("panobinostat",       4),
                  rep("pentamidine",        5),
                  rep("ponatinib",          8),
                  rep(NA,                   2),
                  rep("pralatrexate",       1),
                  rep("raltitrexed",        4),
                  rep("resminostat",        6),
                  rep(NA,                   1),
                  rep("sunitinib",          9),
                  rep(NA,                   4),
                  rep("temozolomide",       1),
                  rep("topotecan",          2),
                  rep("triptolide",         9),
                  rep("vinblastine",        6),
                  rep("vincristine",        1),
                  rep("vindesine",          4),
                  rep("vinorelbine",        5))

association = cbind(compound_name, association)
association = association[!is.na(compound_name)]

overlap = c() # number of overlapping SLC for a specific drug between association and prediction
rank = c() # rank of how high is this overlapped SLC ranked in our prediction, if multiple, mean is calculated
rank_per = c() # rank percentile

for (drug in association[, unique(compound_name)]){
  associated_SLC = association[compound_name == drug, GeneName]
  predicted_SLC = prediction[Compound == drug, SLC]
  overlapped_SLC = intersect(associated_SLC, predicted_SLC)
  overlap = c(overlap, length(overlapped_SLC))
  if (length(overlapped_SLC) == 0){
    rank = c(rank, NA)
    rank_per = c(rank_per, NaN)
  }else if (length(overlapped_SLC) == 1){
    rank = c(rank, paste(which(predicted_SLC == overlapped_SLC), "/", length(predicted_SLC)))
    rank_per = c(rank_per, which(predicted_SLC == overlapped_SLC)/length(predicted_SLC))
  }else{
    rank_ls = vector("numeric", length = length(overlapped_SLC))
    for (i in 1:length(rank_ls)){
      rank_ls[i] = which(predicted_SLC == overlapped_SLC[i])
    }
    rank = c(rank, paste(mean(rank_ls), "/", length(predicted_SLC)))
    rank_per = c(rank_per, mean(rank_ls)/length(predicted_SLC))
  }
}

match_report = data.table(Drug = association[, unique(compound_name)],
                          Overlap_Num = overlap,
                          Rank = rank,
                          Rank_percent = rank_per)

all_prediction = fread("03 Results/06 Drug sensitivity/SCI_dose_correlation.csv")

for (drug in match_report[Overlap_Num == 0, Drug]){
  all_prediction_atomic = all_prediction[Compound == drug][1:52]
  associated_SLC = association[compound_name == drug, GeneName]
  predicted_SLC = all_prediction_atomic[, SLC]
  print(intersect(associated_SLC, predicted_SLC))
}
```
