## Supplementary material for "Predicting substrates for orphan Solute Carrier Proteins using multi-omics datasets": Code for generating results: Meixner predict match 01.R

```
library(imager)
library(data.table)

# Read the heatmap image
heatmap = load.image("01 Raw Data/07 Literature match/cells.png")
channel1 = heatmap[,,1]
channel2 = heatmap[,,2]
channel3 = heatmap[,,3]
channel4 = heatmap[,,4]

# Function to extract color of each cell
extract_cell_color = function(x, y) {
  # Define cell dimensions and grid line width
  cell_width = 78
  cell_height = 18
  grid_line_width = 3.37
  # Calculate coordinates of the cell
  cell_x = (x-1) * (cell_height + grid_line_width) + grid_line_width
  cell_y = (y-1) * (cell_width + grid_line_width) + grid_line_width
  # Extract color of the cell
  cell_color = rgb(red = median(channel1[cell_y:(cell_y + cell_width), cell_x:(cell_x + cell_height)]),
                   green = median(channel2[cell_y:(cell_y + cell_width),cell_x:(cell_x + cell_height)]),
                   blue = median(channel3[cell_y:(cell_y + cell_width),cell_x:(cell_x + cell_height)]),
                   alpha = median(channel4[cell_y:(cell_y + cell_width),cell_x:(cell_x + cell_height)]))
  return(cell_color)
}

# Convert colors to values
Spectrum_value = seq(1,0,-0.04)
Spectrum_color = c("#D40A00","#DB3826","#E24B32","#EC5D3B","#F57747",
                   "#F78C51","#F8A45C","#FCB768","#FBC879","#FCDC8C",
                   "#FCE69A","#FEF2AA","#FDF7BB","#F9F7CA","#F1F5D5",
                   "#E8F4EA","#DAF0F6","#C9E8EF","#B6DEEA","#A6D3E4",
                   "#92C3DB","#80B6D5","#6AA1C9","#5C91C3","#4C7DB8",
                   "#3670B3")

# Function to calculate color distance
color_distance <- function(color1, color2) {
  sum((as.integer(grDevices::col2rgb(color1)) - as.integer(grDevices::col2rgb(color2)))^2)
}

# Function to map cell color to spectrum value
map_color_to_spectrum <- function(cell_color) {
  distances <- sapply(Spectrum_color, function(spectrum_color) color_distance(cell_color, spectrum_color))
  closest_index <- which.min(distances)
  return(Spectrum_value[closest_index])
}

# For each cell of table, record a number
cells_matrix = matrix(NaN, nrow = 115, ncol = 18)

for (x in 1:115) {
  for (y in 1:18) {
    cell_color = extract_cell_color(x, y)
    cells_matrix[x, y] = map_color_to_spectrum(cell_color)
  }
}

y_name = c("MFSD9","SLC35A5","NIPAL3","SLC35F5","SLC35F1",
           "SLC35F2","MFSD11","SLC35C2","MFSD5","SLC35G2",
           "CLN3","SLC35G4","SLC39A11","SLC35F6","TMEM165",
           "SLC35E1","SLC10A4","SLC26A10","SLC10A3","LETMD1",
           "SLC25A51","MFSD14A","MFSD6L","SLC35E2A","SLC35G5",
           "SLC35G6","SLC25A52","SLC25A34","SLC25A35","SFXN5",
           "SLC25A40","SLC23A3","MFSD13A","UNC93B1","SLC35E4",
           "SLC50A1","MFSD3","SLC35A4","SLC25A46","SLC27A3",
           "SLC44A3","ANKH","LETM2","SLC38A10","NIPAL2",
           "SLC10A5","SLC10A7","SLC44A5","SLC22A23","SLC35F4",
           "MFSD1","SLC22A10","SLC36A3","SLC38A11","MFSD8",
           "SLC38A6","SLC9B1","SLC16A6","SPNS3","SLC45A3",
           "MFSD14B","SLC35G1","SLC16A4","SLC38A8","SVOP",
           "SPNS1","DIRC2","SLC16A14","SLC7A4","MFSD4B",
           "SLC22A15","SFXN4","SLC25A39","SLC25A48","SLC35D3",
           "MFSD12","SLC22A17","SLC22A31","SLC35G3","TMEM241",
           "SLC16A13","MFSD4A","SLC46A2","SLC2A6","SLC22A18",
           "SLC35E3","SLC16A5","SLC49A3","SLC2A8","SLC7A13",
           "SLC25A14","SLC25A43","UNC93A","SLC22A25","SLC25A45",
           "SLC35E2B","SLCO5A1","SLCO6A1","SLC6A16","SLC6A17",
           "MFSD6","SLC9C2","OCA2","SLC9A4","SLC45A4",
           "SLC12A9","SV2A","SLC12A8","SV2B","SV2C",
           "SLC7A14","SVOPL","TMEM104","SLC15A5","SLC22A14")
x_name = c("Metal cation","Sodium(+)","Alkali metal cation","L-alpha-amino acid","Elemental halogen",
           "Inorganic anion","Alkaline earth cation","Magnesium(2+)","Potassium(+)","Carbohydrate",
           "Calcium(2+)","Carboxylic acid","Lipid","Steroid","Nucleobase-containing",
           "Divalent metal cation","Transition element cation","Zinc(2+)")

heatmap = data.table(Orphan_SLC = y_name,
                     cells_matrix)
colnames(heatmap) = c("Orphan SLC", x_name)
fwrite(heatmap, "03 Results/07 Heatmap prediction/Meixner Prediction.csv")
```
