## Supplementary material for "Predicting substrates for orphan Solute Carrier Proteins using multi-omics datasets": Code for generating results: Meixner predict match 02.R

```
library(data.table)

review = as.data.table(SLC_ontology_annotation_20240321)
old_review = as.data.table(old_annotation)
prediction = as.data.table(Meixner_Prediction)

orphan_set = c()
for (i in 1:nrow(prediction)){
  find = grep(prediction[i, `Orphan SLC`], review[, HGNC_symbol])
  if (length(find) == 1){
    orphan_set = c(orphan_set, find)
  }else if (prediction[i, `Orphan SLC`] == "MFSD1"){
    find = which(review[, HGNC_symbol] == "MFSD1")
    orphan_set = c(orphan_set, find)
  }else if (prediction[i, `Orphan SLC`] == "SVOP"){
    find = which(review[, HGNC_symbol] == "SVOP (SLC22B4)")
    orphan_set = c(orphan_set, find)
  }else if (prediction[i, `Orphan SLC`] == "MFSD6"){
    find = which(review[, HGNC_symbol] == "MFSD6")
    orphan_set = c(orphan_set, find)
  }
  print(paste(prediction[i, `Orphan SLC`], ":", find))
}

orphan_set = review[orphan_set]

orphan_set_old = c()
for (i in 1:nrow(prediction)){
  find = which(old_review[, HGNC_symbol] == prediction[i, `Orphan SLC`])
  orphan_set_old = c(orphan_set_old, find)
}

old_orphan_set = old_review[orphan_set_old]

term_predict = c()
for (i in 1:nrow(prediction)){
    atomic_data = paste(colnames(prediction[,2:ncol(prediction)])[prediction[i,2:ncol(prediction)]>0.5], collapse = ";")
    term_predict = c(term_predict, atomic_data)
}

prediction_table = data.table(`Orphan SLC` = prediction[, `Orphan SLC`],
                              Prediction = term_predict)
orphan_set[, Prediction := term_predict]
fwrite(orphan_set, "Desktop/Matching predictions.csv")
```
